# Precision fMRI reveals densely interdigitated network patches with conserved motifs in the lateral prefrontal cortex

**DOI:** 10.1101/2025.07.24.666468

**Authors:** Zach Ladwig, Kian Z. Kermani, Youngeun Park, Elena Housteau, Ally Dworetsky, Nathan Labora, Joanna J. Hernandez, Megan Dorn, Derek M. Smith, Derek Evan Nee, Steven E. Petersen, Rodrigo M. Braga, Caterina Gratton

## Abstract

Dominant models of human lateral prefrontal cortex (LPFC) organization emphasize broad domain-general zones and smooth functional gradients. However, these models rely heavily on group-averaged neuroimaging, which can obscure fine-scale cortical features - especially in highly inter-individually variable regions like the LPFC. To address this limitation, we collected a new precision fMRI dataset from 10 individuals, each with approximately 2 hours of resting-state and 6 hours of task data. We mapped individual-specific LPFC networks using resting-state fMRI and tested network-level functional preferences using task fMRI. We found that individual LPFC organization differed markedly from group-averaged estimates. Individual maps showed more fragmented and interdigitated networks - especially in anterior LPFC - including novel conserved motifs present across individuals. Task fMRI revealed that distinct but adjacent networks support domain-specific processes (i.e., language, social cognition, episodic projection) versus domain-general control. Sharp functional boundaries were visible at the individual level that could not be observed in group data. These findings uncover previously hidden organizational principles in the LPFC and offer a framework for understanding how the LPFC supports flexible, complex cognition through a finely organized architecture.

## INTRODUCTION

The lateral prefrontal cortex (LPFC) plays a central role in high-level cognitive processes like planning, reasoning, and problem-solving (Fuster, 1989; Stuss & Benson, 1986). Damage to the LPFC disrupts many aspects of goal-directed behavior (Lhermitte et al., 1986; Milner, 1963; Shallice & Burgess, 1991), and abnormalities in LPFC circuitry have been linked to a wide range of neurological and psychiatric disorders (Barbalat et al., 2009; McTeague et al., 2017).

Despite its central role in cognition, there is little consensus on how the LPFC is functionally organized. A major point of debate concerns the degree and topography of functional specialization within the LPFC. Several dominant models portray the LPFC as a broad, multifunctional region with gradual gradients of specialization. For instance, large-scale neuroimaging studies have reported overlapping LPFC activations across diverse tasks (Cabeza & Nyberg, 2000; Duncan & Owen, 2000; Niendam et al., 2012), supporting unitary models of flexible cognitive control (Duncan, 2010; Miller & Cohen, 2001). Others have described smooth topographic gradients, such as a rostral–caudal axis of control-related abstraction (Badre, 2008; Koechlin et al., 2003; Nee & D’Esposito, 2016) and a dorsal–ventral domain shift from spatial to verbal content (Abdallah et al., 2022; Blumenfeld et al., 2013; Rahm et al., 2013). Functional network-based frameworks have identified broad network territories spanning the LPFC (Power et al., 2011; Yeo et al., 2011), dominated by a large frontoparietal network region that is ascribed much of the high-level planning and control functions for which the LPFC is known (Vincent et al., 2008). These networks too have been interpreted along a slowly moving cortex-wide gradient from sensorimotor to default mode regions (Margulies et al., 2016). However, these smooth patterns stand in contrast with studies based on neuronal recordings or tract tracing of neuroanatomical circuits, which have identified adjacent regions with different cytoarchitectonics and neuronal preferences (Barbas & Pandya, 1989; Funahashi et al., 1993; Petrides, 2005).

We contend that this discrepancy may arise because most neuroimaging studies of human LPFC organization have relied on group-averaged data, aligning individual brains to a common anatomical template and assuming functional regions occur in similar locations across individuals. Recent work has shown that group averaging can obscure fine-scale details and overestimate overlap between distinct functional regions, particularly in areas with high inter-individual variability (Braga & Buckner, 2017; Du et al., 2024; Fedorenko, 2021; Gordon et al., 2017; Petersen et al., 2024). The LPFC has emerged as one of the most inter-individually variable brain regions (Dworetsky et al., 2024; Mueller et al., 2013; Seitzman et al., 2019), raising the possibility that the LPFC may contain fine-scale organization not reflected in current models.

To address the limitations of group-averaged methods, several studies have adopted a “precision neuroimaging” approach—collecting large amounts of data from individual participants and analyzing functional organization on a subject-by-subject basis (Fedorenko, 2021; Gordon et al., 2017; Laumann et al., 2015; Michon et al., 2022). In the LPFC, several such studies have identified examples of fine-scale functional specialization not visible in group-averaged models (DiNicola et al., 2020; Fedorenko et al., 2012; Fedorenko & Thompson-Schill, 2014; Michalka et al., 2015; Noyce et al., 2017) and found that functional differentiation often corresponds with distinct functional networks (Blank et al., 2014; Braga et al., 2020; DiNicola et al., 2023; Du et al., 2024; Salvo et al., 2025; Tobyne et al., 2017).

Here, we sought to extend these observations and map the fine-scale organization of the individual LPFC. To do this, we collected a new high-resolution precision fMRI dataset (PAN: Precision targeting of Association Networks) from ten individuals (18-33 years; 5F), each with two hours of resting-state fMRI and 6 hours of task fMRI. The task battery spanned 11 different cognitive demands known to engage the LPFC including theory of mind, episodic projection, language processing, and eight distinct cognitive control tasks. Using resting-state fMRI, we derived high-resolution, individual-specific networks and characterized fine-scale features present in the individual-specific compared to group-averaged LPFC. Using task fMRI, we evaluated how individual-specific LPFC networks supported diverse cognitive demands. Finally, we used a separate publicly available precision fMRI dataset (Natural Scenes Dataset; Allen et al., 2022), to replicate the fine-scale network features identified in our dataset.

We found that individual-specific LPFC network organization systematically differed from group-averaged models. While group-averaged data consisted of large contiguous network regions, individuals exhibited fragmented and interdigitated networks. In individual-specific data, the frontoparietal network was systematically smaller and borders between association networks were denser – especially in anterior LPFC. Despite variability in the exact positioning of networks, we observed conserved spatial motifs across individuals that could not be seen viewing group data alone, as well as clearly idiosyncratic features. Using task fMRI, we found that some LPFC networks supported different domain-specific processes, while others supported domain-general control. Individual-specific task activations corresponded closely with individual-specific networks and revealed sharp functional boundaries that would not have been visible in group-level data. Together, these findings refine coarse and gradient models of the LPFC, emphasizing fine-scale architecture and underscoring the importance of individual-level analysis in the LPFC for research and clinical applications.

## RESULTS

### Individual-specific LPFC network and task maps are reliable and unique

Using at least 100 minutes of motion-censored resting-state fMRI data per individual, we generated individual-specific network parcellations for 10 individuals (Figure 1A, whole-cortex in Supplemental Figure S1, S2) using a state-of-the-art network identification protocol (Lynch et al., 2024). A group-averaged estimate was also derived for comparison from a reference set of 37 individuals (Lynch et al., 2024, see Supplemental Figure S3 for comparison with a group-averaged map derived from the 10 individuals in this study). For most analyses reported in this manuscript, network maps were masked to focus on the lateral prefrontal cortex (LPFC). To establish the reliability and individual specificity of individual-level LPFC network parcellations, we calculated split-half reliability for two individuals (PAN01 and PAN02) who completed additional resting-state runs to facilitate reliability analyses (totaling 147 and 159 motion-censored minutes respectively). As shown in Supplemental Figure S4, their individual-specific LPFC parcellations were reliable across split halves derived from independent scanning sessions (mean network Dice overlap = 0.79 ± 0.05 between split halves of the same individual) and unique (mean network Dice overlap = 0.34 ± 0.13 between split halves of different individuals). A paired-samples t-test revealed that within-individual similarity was significantly higher than between-individual similarity (t(6) = 9.8, p < 0.0001).

**Figure 1:**
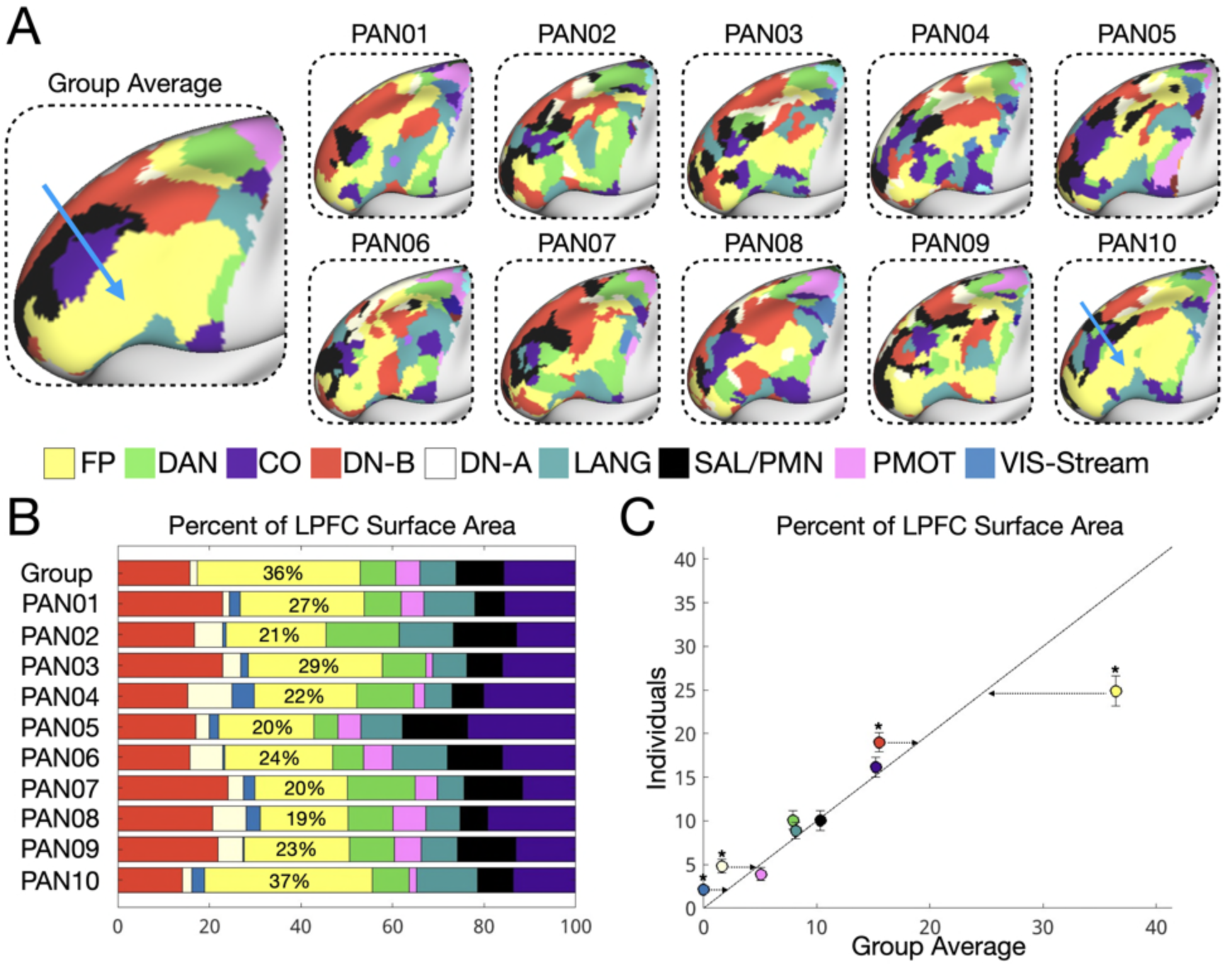
LPFC network organization differs systematically between individuals and the group average, especially in the frontoparietal network. (A) Individual-specific network parcellations are shown, alongside the group average. While the group-averaged map consisted of broad contiguous regions, individual maps were fragmented and interdigitated (see Supplemental Figure S6 for replication with an alternative group-averaged parcellation). This difference was especially pronounced in the frontoparietal network, which was large and contiguous in the group-average data (blue arrow) but was smaller and fragmented in individuals (except for PAN10, blue arrow). (B,C) On average, frontoparietal network size in the LPFC was overestimated in the group-averaged networks (t(9) = 6.7, p = 0.00009) while default B (t(9) = -3.1, p = 0.013), default A (t(9) = -4.1, p = 0.003), and visual-stream (t(9) = -4.3, p = 0.0008) size were underestimated.

In addition, we collected ∼6 hours of task fMRI data per individual (30-60 minutes per task) and generated individual-specific z-statistic maps for three cognitive domains of interest (theory of mind, language processing, and episodic projection) as well as for eight distinct cognitive control tasks (visual n-back, auditory n-back, spatial memory span, verbal memory span, numeric multi-source interference, verbal multi-source interference, visual attention, auditory attention). To validate the reliability of individual-level cognitive control maps, we examined two individuals (PAN01 and PAN02) who completed approximately twice as many runs per task (50-100 minutes per contrast) as the eight other individuals. Shown in Supplemental Figure S5, their z-statistic maps were reliable across split halves (r = 0.85 ± 0.06 within individuals) and unique (r = 0.39 ± 0.09 between individuals). A paired-samples t-test revealed that within-individual similarity was significantly higher than between-individual similarity across tasks (t(7) = 21.6, p < 0.000001), suggesting that LPFC task responses were reliable and individual specific.

In the next sections, we characterized the topographical features of LPFC organization visible at the individual level relative to the group average. Specifically, we: (1) characterized the areal composition and spatial density of distinct networks in the individual LPFC (Figures 1,2), (2) described the architecture of conserved network motifs not visible in group-level data (Figures 3,4), and (3) described how this network architecture supports a variety of high-level cognitive demands (Figures 5–8).

**Figure 2.**
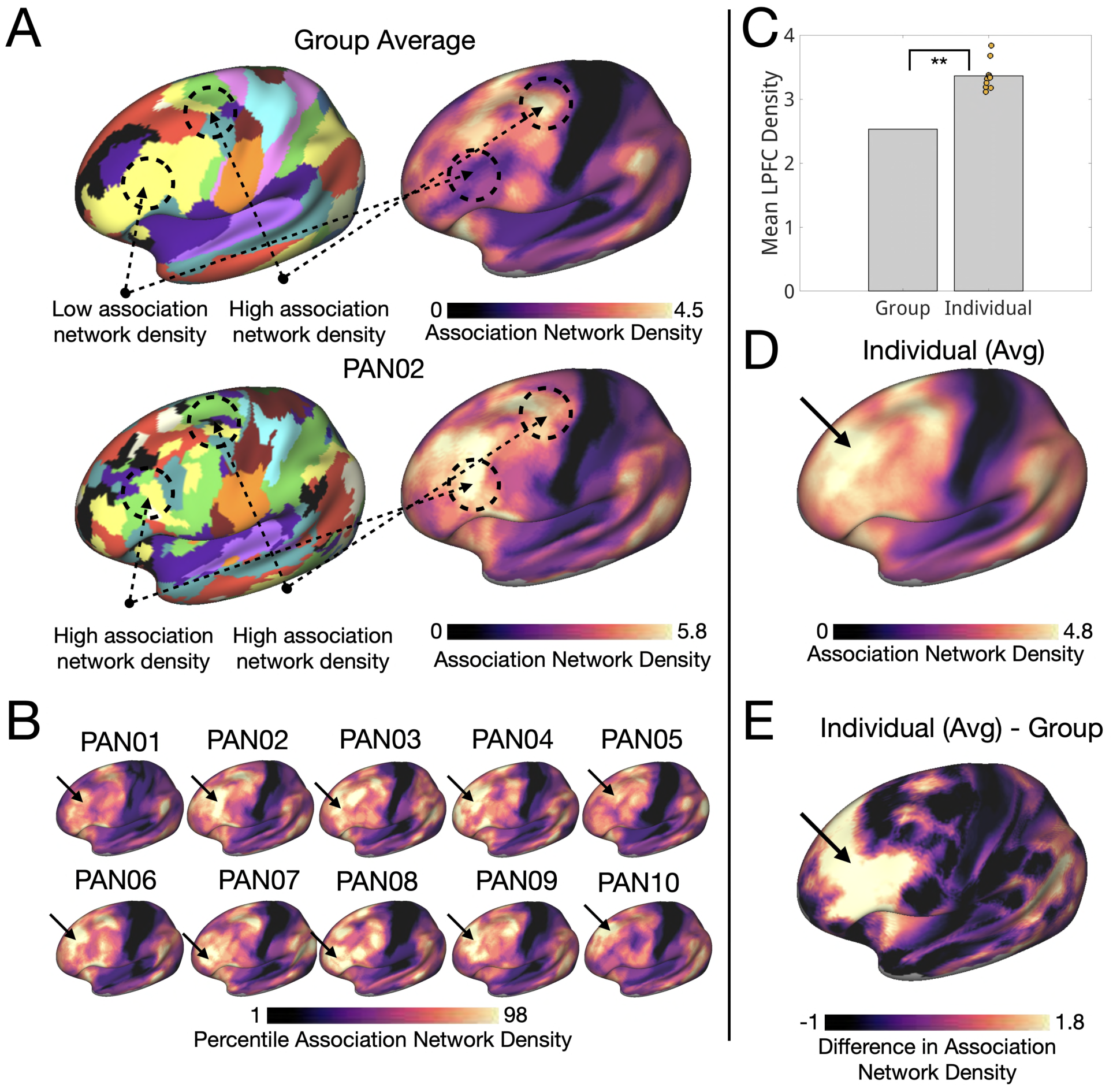
Association network density is underestimated in LPFC by group averaging, especially in anterior LPFC. (A) Association network density was computed for each cortical vertex as the number of unique association networks within a 6–14 mm radius (averaged across distances of 6, 8, 10, 12, and 14 mm). (B) Individual maps of association network density are shown with an arrow highlighting the anterior high-density region. (C) Mean LPFC association network density was significantly higher in individuals compared to the group-averaged networks (One-sample t-test: t(9) = 11.5, p < 0.000001). (D) On average, individuals exhibited a distinct spatial pattern, including a region of high density in anterior LPFC. (E) This anterior pattern was absent in the group-averaged map, suggesting that association network density is particularly underestimated in anterior LPFC. (See also whole cortex estimates in Supplemental Figure S8 and all-network density estimates in Supplemental Figure S11.) These results were replicated using a different group-averaged prior, suggesting they are not specific to this parcellation (Yeo 17 networks, Supplemental Figure S10).

**Figure 3.**
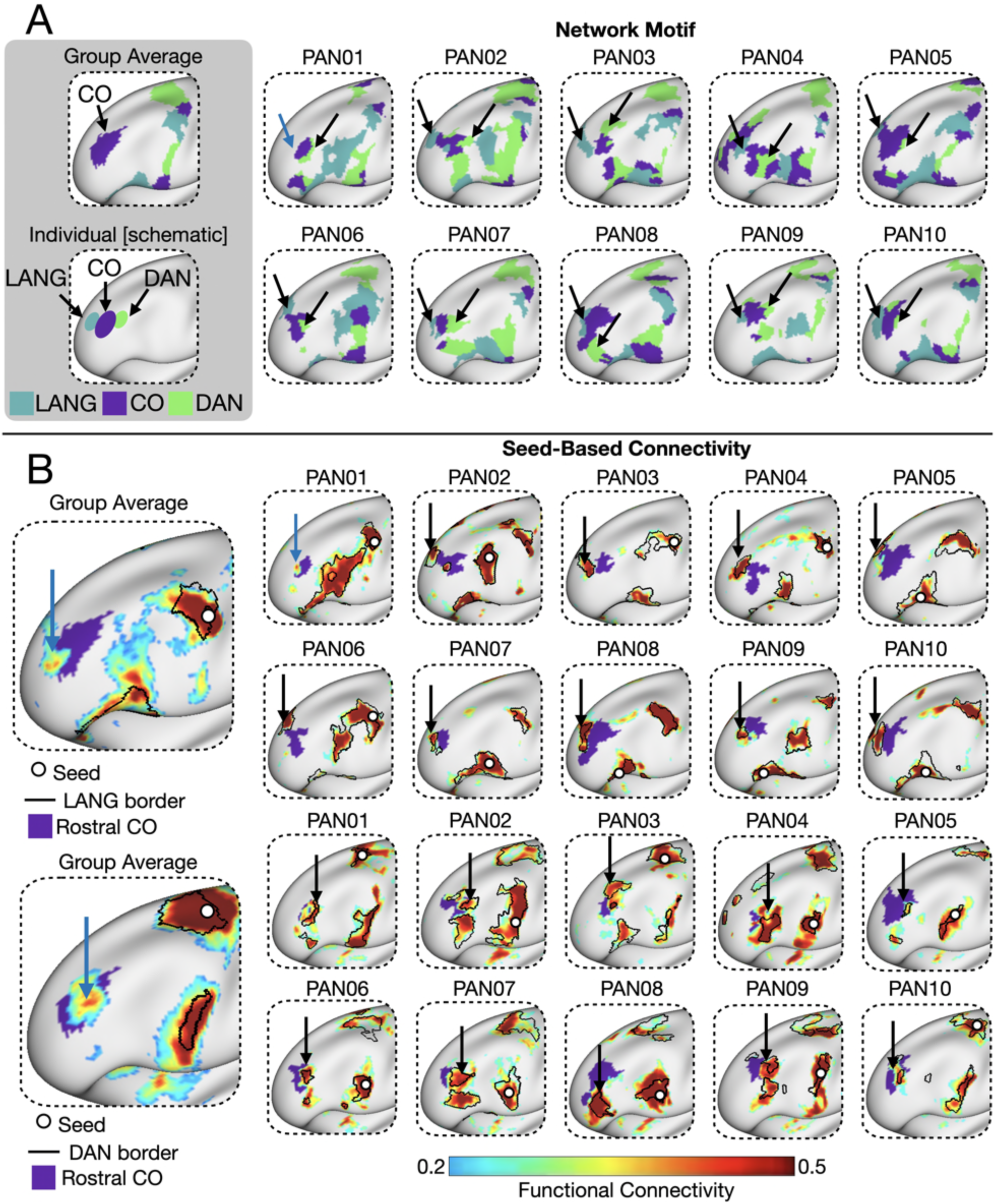
A consistent anterior LPFC motif involving language, cingulo-opercular, and dorsal attention networks is present across individuals but absent in the group average. (A) Individual-specific parcellations revealed a consistent spatial motif in 9 out of 10 individuals: a language network region anterior to rostral cingulo-opercular network and dorsal attention network region posterior to it. This motif was not observed in the group-averaged parcellation. (B) Seed-based connectivity from other frontal regions of language and dorsal attention showed connectivity with these rostral regions in individuals, supporting their network identity. Weak but present connectivity was also observed in the group-averaged map (blue arrows). The weakness may be explained by the variability in location/orientation of strong connectivity across people, hence why these features were not identified in the group-averaged parcellation.

**Figure 4:**
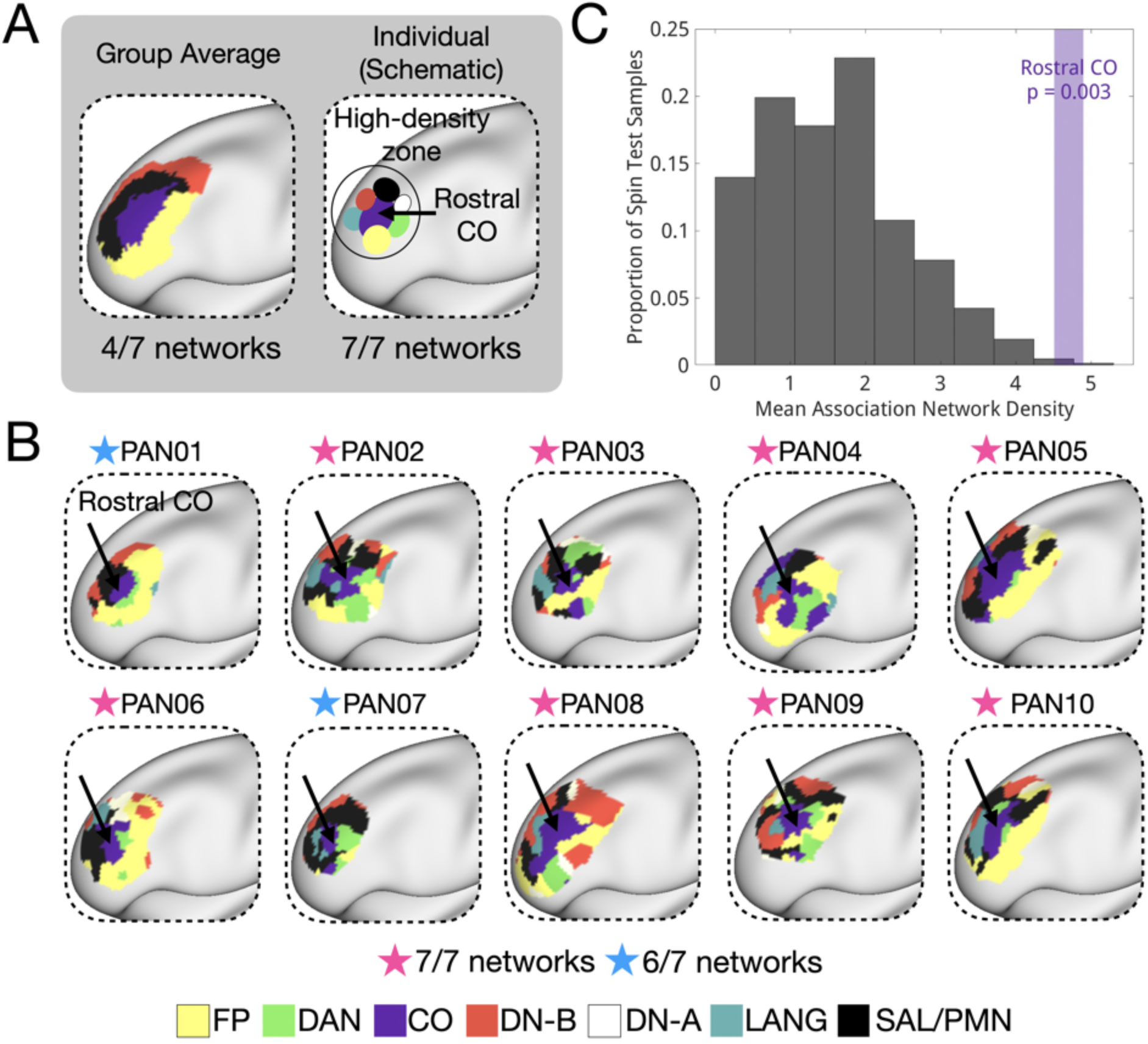
An anterior PFC high-density zone including all 7 association networks is consistently observed across individuals. (A) In the group average, the anterior LPFC contains a CO network region bordered by FP, DN-B, and SAL/PMN. In individuals, we often observed regions of all 7 association networks here, underlying the high-density zone observed in Figure 2. (B) This pattern was observed in 8/10 subjects. PAN01 and PAN07 did not exhibit rostral DN-A regions but showed seed-based DN-A connectivity to this location (Supplemental Figure S14). All individuals are masked to show networks within 15mm from the rostral CO region. (C) The rostral CO network region exhibited significantly higher association network density versus a matched randomly rotated null distribution (p = 0.003).

**Figure 5:**
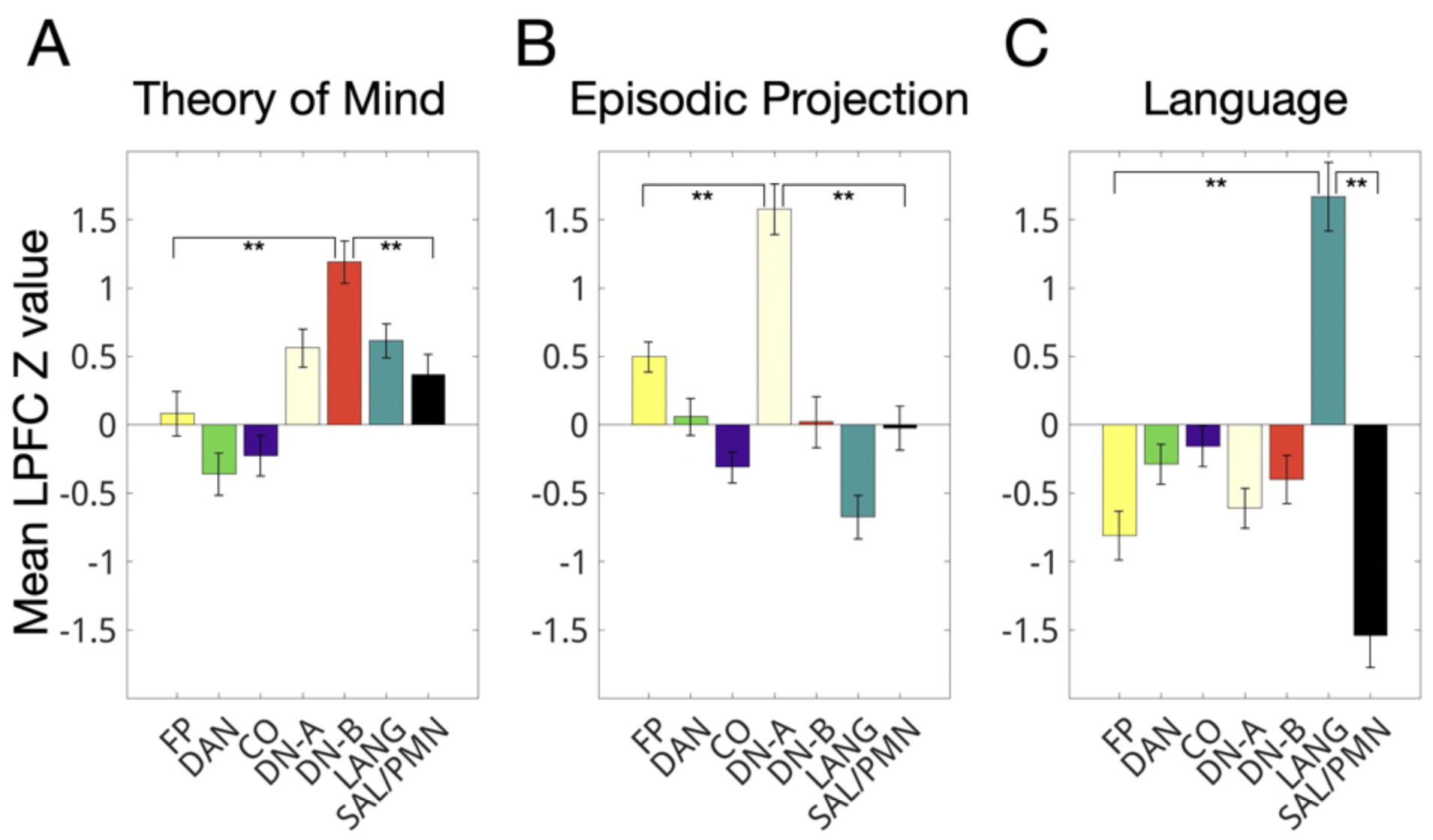
Theory of mind, episodic projection, and language demands preferentially recruit distinct individual-specific LPFC networks. Average z-statistics were computed for LPFC regions of each individual-specific network for theory of mind, episodic projection, and language processing tasks. Paired t-tests comparing the target network to each of the other LPFC networks revealed that: (A) theory of mind preferentially activated default B (all comparisons, corrected p < 0.01), (B) episodic projection preferentially activated default A (all comparisons, corrected p < 0.00005), and (C) language processing preferentially activated the language network (all comparisons, corrected p < 0.001).

**Figure 6:**
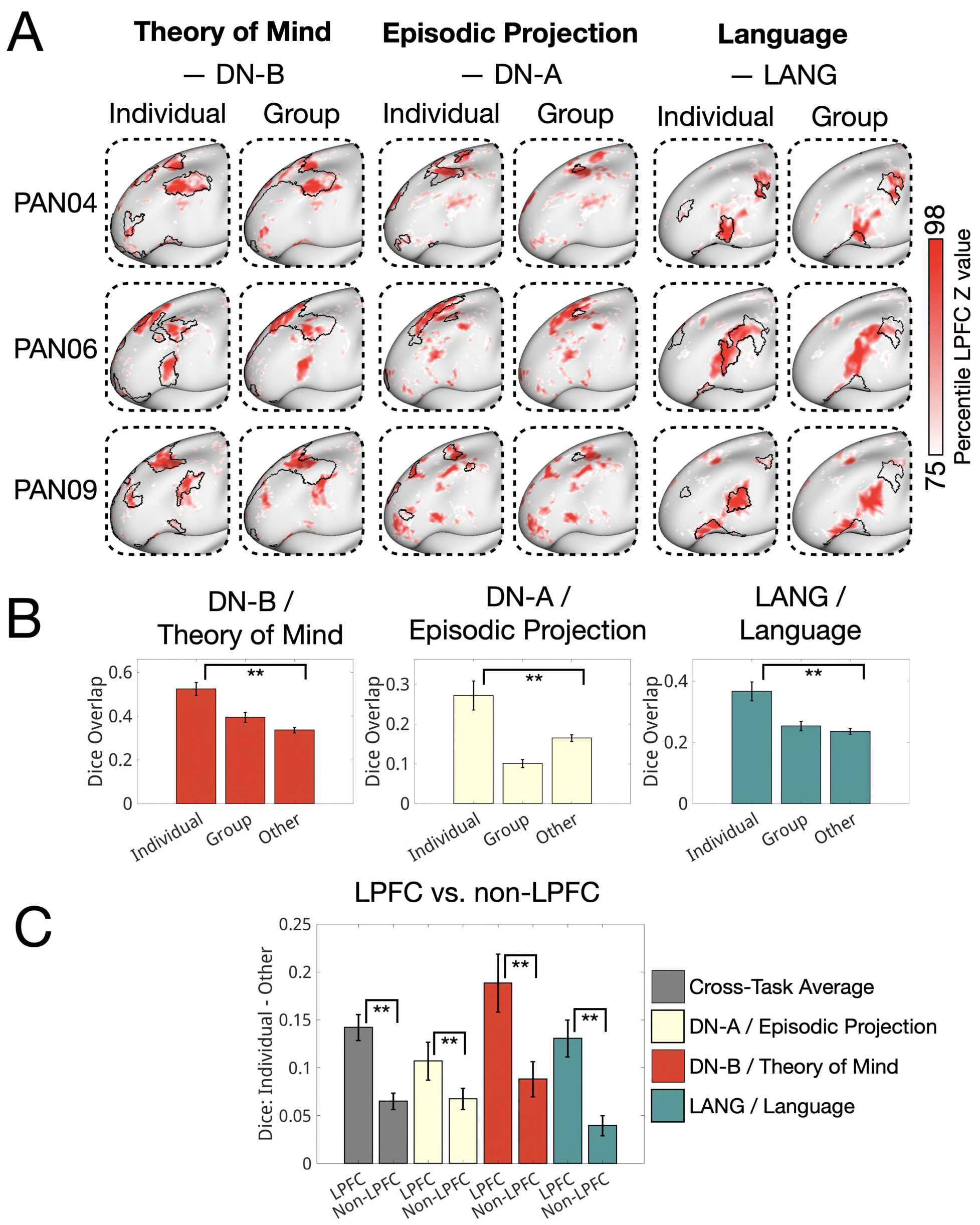
LPFC domain-specific task activations are patchy and correspond with individual-specific network boundaries. (A) The top 25% most active LPFC vertices (ranked by z-statistic) are shown for three exemplar individuals for theory of mind, episodic projection and language demands, overlaid with both individual-specific (left column) and group-averaged (right column) network boundaries. All individuals are shown in Supplemental Figure S17. (B) Average dice overlap was calculated between thresholded (top 15%, 20%, 25%) LPFC task activation maps and individual-specific, group-average, and non-specific individual parcellations. Task activations showed significantly greater overlap with individual-specific LPFC networks than with either group-averaged networks or non-specific individual networks (corrected p < 0.05 for all comparisons). (C) This effect was larger in the LPFC than in non-LPFC cortical regions (p < 0.0001).

**Figure 7:**
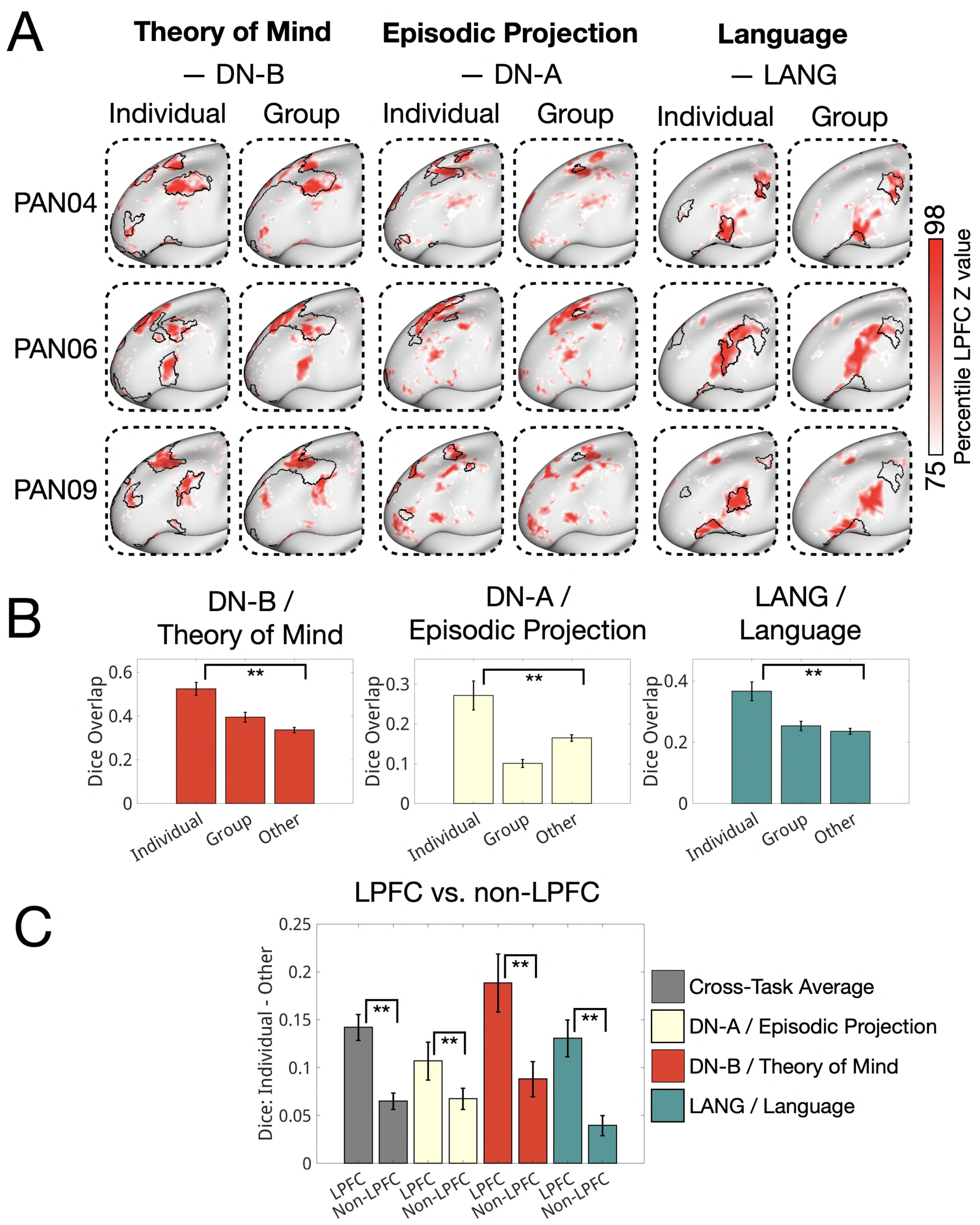
Language regions are embedded unexpectedly within mid-LPFC canonical frontoparietal territory in a subset of individuals. (A) Language network parcellations for 10 individuals and the group-averaged data are shown, overlaid with the group-averaged frontoparietal network. Four individuals (PAN01, PAN02, PAN06, PAN09) exhibited large language regions in mid-LPFC, well outside typical language territory. (B) These variant language regions were interdigitated with fragmented individual-specific FP regions. (C) Variant regions showed strong seed-based connectivity with other LPFC language network regions, supporting their network identity. (D) Language processing task activations showed positive responses in the variant language regions but not in adjacent FP regions. (E) Spatial working memory activations showed the opposite pattern: activation in adjacent FP regions but not in the variant Language regions. (F) Language > spatial working memory *z*-values are shown for the variant individuals and for the same ROI locations averaged across the six non-variant individuals. Variant-region ROIs in non-variant individuals did not show language > spatial working memory activity, validating the individual-specificity of variants.

**Figure 8:**
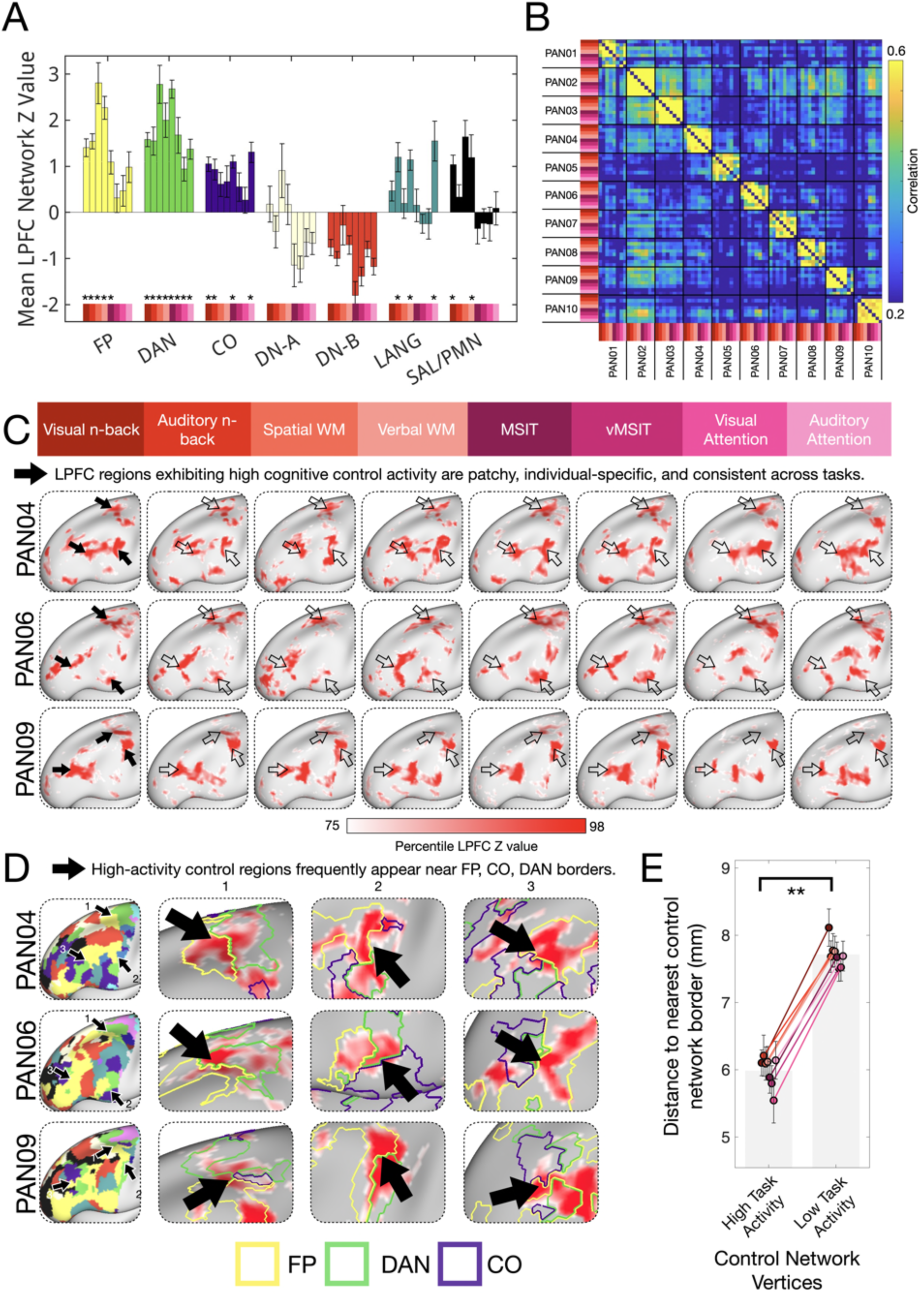
Cognitive control tasks strongly recruit a set of distributed, individual-specific regions concentrated along the borders of frontoparietal, cingulo-opercular, and dorsal attention networks in the LPFC. (A) Mean network-level LPFC activations (z value) are shown for all eight cognitive control tasks. Bars are grouped by network and colored accordingly. Statistically significant activations (corrected p < 0.05) are marked with asterisks. The dorsal attention (DAN), frontoparietal (FP), and cingulo-opercular (CO) networks were the most consistently engaged across tasks (8/8, 5/8, and 4/8 tasks, respectively). Language and salience/parietal memory networks showed more selective engagement (3/8 and 2/8 tasks). (B) Vertex-wise activation patterns were individual-specific (r = 0.27 ± 0.03 between individuals for the same task) and relatively more consistent within individuals across tasks (r = 0.58 ± 0.03), indicating a stable individual-specific activation motif across domains. (C) The top 25% most active LPFC vertices are shown for each task in three exemplar individuals (same individuals as Figure 6, all individuals shown in Supplemental Figures S23 and S24). Task responses consistently involved a distributed set of regions which was similar across tasks. Individual arrow locations are consistent across tasks. (D) High cognitive control task activity tended to appear near the borders of control networks. Zoom-ins of control task activity (red) on top of network borders are shown for the vMSIT task for three exemplar individuals (all individuals shown in Supplemental Fig. 23 & 24). (E) To quantify this border proximity effect, we measured the geodesic distance from each LPFC vertex in FP, DAN, or CO to the nearest vertex belonging to a different control network. Highly active vertices (z > 2) were significantly closer to other control networks than less active vertices (z < 2) in all eight tasks (all p < 0.001). This was replicated at two other thresholds (Supplemental Figure S26). A reverse effect was often seen for domain-specific tasks (Supplemental Figure S27).

### LPFC network organization is systematically different in individuals, especially the frontoparietal network which is overestimated in group-averaged data

Shown in Figure 1A, we observed systematic differences between individual LPFC networks and the group average. While a similar set of association networks were present in individuals and the group-averaged data (frontoparietal, default A, default B, cingulo-opercular, dorsal attention, salience/parietal memory, language), both the relative size and spatial patterning of LPFC network regions were different. In the group-averaged data, network regions tended to be large and contiguous, while in individuals, network regions were often smaller and fragmented.

This difference was especially stark in the frontoparietal network. In the group-averaged data, the frontoparietal network made up 36% of LPFC surface area and contained a large contiguous network region (Figure 1A). In individuals, the frontoparietal network was significantly smaller (25% ± 5.4% of LPFC surface area; one-sample t-test: t(9) = 6.7, p = 0.00009, see Figure 1B,C) and was frequently fragmented into multiple distinct regions. Only 1/10 individuals had a large and contiguous frontoparietal network as was seen in the group-averaged data (PAN10, see blue arrow in Figure 1A). This result was replicated using an alternative group atlas which included two frontoparietal networks (Yeo 17 network parcellation, Supplemental Figure S6). By contrast, default B (t(9) = -3.1, p = 0.013), default A (t(9) = -4.1, p = 0.003), and visual-stream (t(9) = -4.3, p = 0.0008) networks were larger in individuals than in the group-averaged data (Figure 1B). These findings suggest the LPFC exhibits a more distributed and fragmented network topography than was previously assumed, with more cortical surface area dedicated to networks (e.g., default A/B) not typically associated with cognitive control – a topic we will return to later. We further validated network assignments using seed connectivity for PAN01 and PAN02 to ensure the observed organization was not an artifact of thresholding or misassignment in the network parcellation (Supplemental Figure S7).

### The individual LPFC is densely packed with association network borders, especially in the anterior LPFC

As described above, LPFC organization in individuals exhibited interdigitated networks compared with the relatively contiguous networks in the group-averaged data. Prior work has speculated that network interdigitation could have functional consequences (Assem et al., 2024; Power et al., 2013). To compare interdigitation in the LPFC of individuals relative to the group average, we defined “association network density” as the average number of distinct association networks within 6-14 mm of each vertex (Figure 2A,B based on “community density” from Power et al., 2013). This metric included canonical association networks (Dorsal Attention [DAN], Frontoparietal [FP], Cingulo-Opercular [CO], Default-B [DN-B], Default-A [DN-A], and Salience/Parietal Memory [SAL/PMN] networks), and excluded sensory-motor processing networks (Somatomotor, Premotor, Visual, Auditory, and SCAN networks). Notably, when defining networks, we removed implausibly small regions (< 50 mm^2^) to ensure this metric was not inflated by spurious network assignments.

As shown in Figure 2C, association network density was consistently higher in individual-specific LPFC networks (mean density = 3.36 ± 0.22) than in the group-averaged networks (mean density = 2.53, one-sample t-test: t(9) = 11.5, p < 0.000001). In addition to an overall increase, individuals exhibited a distinct spatial pattern: all individuals showed a hotspot of high density in anterior LPFC that was not observed in the group-averaged map (Figure 2B,D,E). This was particularly apparent in the difference map between the individual and group-averaged density maps (Figure 2E) and in the whole-cortex view (Supplemental Figure S8). To ensure that this difference was not driven by BOLD signal differences within the LPFC, we examined the spatial BOLD signal pattern across individuals in the LPFC and found that the correlation between BOLD signal and association network density was near zero (r = 0.018, 95% CI: [–0.002, 0.039]), suggesting no meaningful relationship (Supplemental Figure S9). As with network composition, these results were replicated using an alternative group network prior (Yeo 17, Supplemental Figure S10). Network density maps were also computed across all functional networks (not only association networks) and yielded similar results (Supplemental Figure S11). These findings indicate that network interdigitation is systematically underestimated in LPFC group-averaged maps, particularly in the anterior LPFC.

Examining whole-cortex maps (Supplemental Figure S8), we observed that the LPFC appeared to be uniquely interdigitated compared to other association regions of cortex. To formally test this observation, we defined ROIs for four major association cortex regions—lateral temporal, lateral parietal, medial prefrontal, and lateral prefrontal cortex—and compared their association network densities. Consistent with our observation, the LPFC exhibited significantly higher association network density than each of the other three regions (Supplemental Figure S12; all paired t-tests, p < 0.001), though locally elevated association network density was also observed in the temporoparietal junction (Supplemental Figure S8). Together, these results indicate that although network interdigitation is not unique to the LPFC, it is most pronounced and spatially extensive within the LPFC.

### Individual-specific maps reveal conserved network motifs not present in group-averaged maps

To better understand the spatial patterns underlying the increased association network density in the LPFC, we manually examined network organization in each individual. We asked whether the patterns were completely idiosyncratic or if there were consistent features—conserved network motifs—that appeared across people. Prior work using precision fMRI has shown that such motifs can be found across individuals but are too fine-grained or variable in location to show up in group-averaged data (Braga & Buckner, 2017; Gordon et al., 2023; Noyce et al., 2017). In line with this, we identified and validated two novel spatial motifs in the LPFC that were shared across most individuals but not visible in the group average. We also identified some individual-specific network features, described later.

First, we identified a novel conserved network motif surrounding the rostral cingulo-opercular network region. In group-averaged data, the rostral cingulo-opercular network region was bordered by the frontoparietal network posteriorly and by the salience/parietal memory and default B networks anteriorly. However, in individuals, additional small regions of the language and dorsal attention networks were present (Figure 3A), forming a three-network motif. Although the shape and extent of these regions varied across individuals, they were visible in 9/10 and 10/10 individuals, respectively. These findings were validated with seed-based functional connectivity analyses (Figure 3B), suggesting they were real features of functional organization rather than artifacts of our parcellation method. We also found additional instances of this three-network motif in the caudal strip of the LPFC (Figure 1A), the temporoparietal junction (TPJ) and pre-supplementary motor area (pre-SMA) (Supplemental Figure S13), aligning with previous observations that functional networks often border the same neighbors in multiple cortical regions (Braga & Buckner, 2017).

There were two exceptions to this three-network-motif among the ten individuals. PAN01 exhibited the anterior dorsal attention region but not the anterior language region. However, their seed connectivity suggested the presence of an anterior language region directly adjacent to CO not captured by the parcellation (Figure 3B, blue arrow). PAN04 displayed two anterior CO regions. The more posterior CO region followed the motif—language anterior, dorsal attention posterior—whereas the more anterior region showed the reverse (Figure 3A).

Second, we observed that this dorsal attention – cingulo-opercular – language motif was part of a larger organizational feature in the same location. In 8 out of 10 individuals, the anterior PFC contained a high-density zone where regions of all seven association networks (dorsal attention, cingulo-opercular, frontoparietal, salience/parietal memory, default A, default B, language) appeared in close proximity, with the cingulo-opercular region near its center (Figure 4A). Two individuals (PAN01, PAN07) did not exhibit a rostral DN-A region in their network parcellations but did show seed-based connectivity to this location from other DN-A regions (Supplemental Figure S14).

This location aligned with the hotspot of association network density in Figure 2D and was the only region in the LPFC where the salience/parietal memory network consistently appeared across individuals (Figure 1A). To assess the unique properties of the rostral CO location, we compared the association network density of the rostral CO region to a cortex-wide rotational null model. On average across individuals, this region had significantly higher association network density compared to the matched randomly rotated null (p = 0.003, Figure 4C). A whole-brain map of association network density further highlights the prominence of this location, which stands out relative to most other brain regions (Supplemental Figure S8; note that another high-density region was also observed in the temporoparietal junction). The gathering of many association networks at specific sites may have important functional implications, as we later discuss.

### Fine-scale network structure is replicated in a publicly available high-resolution 7T precision fMRI dataset

To confirm that the fine-scale network structure we observed in the PAN dataset was not specific to this set of individuals, we replicated the primary findings from Figures 1–4 in an independent, publicly available, 7T precision fMRI dataset (Natural Scenes Dataset; Allen et al., 2022). As can be seen in Supplemental Figure S15, NSD subjects also exhibited a smaller frontoparietal network versus the group average (Group Average: 35.5%, Individual Average = 23.9% ± 2.5%, one-sample t-test t(5) = 11.6, p < 0.0001), higher LPFC association network density versus the group average (Group Average = 2.53, Individual Average = 3.08 ± 0.02, one-sample t-test t(5) = 19.1, p < 0.00001), the DAN-CO-LANG motif (5/6 individuals) and a rostral CO location of higher-than-expected association network density (p = 0.004). This replication provides support that fine-scale network structure in the LPFC is not specific to our sample but is likely a common feature of brain networks that can be observed in high resolution precision fMRI data.

### Language, theory of mind, and episodic projection tasks preferentially activate distinct LPFC networks adjacent to – but separate from – cognitive control networks

Up to this point, we have characterized the individual-specific LPFC based on the spatial topography of large-scale networks defined by resting-state functional connectivity. We now turn to task fMRI to examine how these networks support distinct high-level cognitive demands. While the LPFC is classically associated with domain-general cognitive control, recent studies have shown that domain-specific processes —including theory of mind, episodic projection, and language processing—preferentially engage networks which include LPFC regions: default B, default A, and language networks (Braga et al., 2020; DiNicola et al., 2020, 2023; Du et al., 2024; Salvo et al., 2025). Close examinations of task activations with individuals has often shown these functionally specialized regions lay adjacent to, but not overlapping with, nearby cognitive control-related regions (DiNicola et al., 2023; Fedorenko et al., 2012). Notably, these sharp anatomical distinctions are obscured in group-averaged data.

Here, we examined individual-specific LPFC responses to theory of mind, episodic projection, and language processing using well-established localizer tasks (see Methods for task details). We first assessed network-level functional preferences within the LPFC, aiming to replicate prior findings that theory of mind, episodic projection, and language processing preferentially engage the default B, default A, and language networks, respectively. Using paired t-tests, we compared the mean z-values of LPFC regions in each target network to those of all six other association networks. The results replicated prior network-level dissociations: theory of mind preferentially activated LPFC default B regions (corrected p < 0.01 for all comparisons), episodic projection preferentially activated LPFC default A regions (corrected p < 0.00005), and language processing preferentially activated LPFC language network regions (corrected p < 0.001; Figure 5A). Full statistical details—including corrected and uncorrected *p*-values, *t*-statistics, and Cohen’s *d*—are provided in Supplemental Table S1. To test whether network preferences could be due to the spatial properties of the networks, we validated these results with a whole-brain spatial permutation test. In all cases, target network activation was significant relative to the spatial null distribution (all p < 0.05, Supplemental Figure S16).

We next assessed how individual-specific LPFC task activations corresponded with individual-specific versus group-averaged network boundaries. As described earlier, individual-specific LPFC networks differed systematically from group-averaged maps—they were more fragmented, interdigitated, and featured a smaller frontoparietal region. Figure 6A shows three example individuals, with the top 25% of activated LPFC vertices for each task overlaid on both individual-specific and group network boundaries (see Supplemental Figure S17 for all individual-specific maps). We observed that task activations were patchy and idiosyncratic, and aligned more closely with individual-specific than group-averaged network boundaries.

To quantify these observations, we calculated the average Dice overlap between thresholded task activations (top 15%, 20%, 25% of LPFC vertices by z-statistic) and network boundaries for individual-specific, group-average, and individual-non-specific networks. We compared thresholded task activations to the boundaries of the single network each task activated most strongly (Language → LANG; Theory of Mind → DN-B; Episodic Projection → DN-A; note that this choice is a simplification, as tasks, including Theory of Mind for example, engage multiple networks at different levels, as shown in Figure 5). As shown in Figure 6B, task activations overlapped more with individual-specific networks than with either group-averaged or individual-non-specific networks across all three tasks (corrected p < 0.05 for all comparisons; see Supplemental Table S2 for full statistics). This replicated prior findings that task activations align with individual-specific networks (Braga et al., 2020; DiNicola et al., 2020; Gordon et al., 2017; Salvo et al., 2021). Notably, although this pattern was evident across the cortex, the effect was significantly stronger in the LPFC than in other regions (Figure 6C), consistent with the idea that the LPFC exhibits particularly high individual variability and contains fine-scale organization not captured in group-averaged data. As with the network topography results, we replicated these task results using the Yeo 17 network prior to ensure they were not specific to our parcellation (Supplemental Figure S18).

Finally, we observed that individual-specific activations for these three domains closely matched individual network boundaries even when these networks appeared in unexpected locations. For example, a subset of individuals displayed extensive, unexpected segments of the language and default B networks interdigitated within discrete FP territories in the mid-LPFC (Figure 7A, Supplemental Figure S19) far from their expected group-averaged locations.

Language network regions were found within the canonical group-averaged frontoparietal territory of the LPFC in 4 out of 10 individuals (Figure 7A). No comparable regions could be observed in the other 6 individuals, suggesting this is not a conserved motif. In all 4 cases, the variant regions were adjacent to individual-specific FP regions (Figure 7B) and showed strong seed-based functional connectivity with other LPFC language network territories (Figure 7C), indicating they were not artifacts of the parcellation. Functionally, these variant language network regions responded robustly to language processing demands (Figure 7D) and did not respond to spatial working memory demands. By contrast, nearby individual FP regions showed the opposite pattern—activation during spatial working memory but not language processing (Figure 7E). To test that the other six individuals did not have an analogous functional region that was missed in our parcellation, we created regions of interest (ROIs) matched to the size and location of the mid-LPFC language region of these 4 individuals. We tested the functional activity of these ROIs in the 6 individuals without mid-LPFC language regions. We found that these ROIs did not show stronger responses to language than to spatial working memory, further supporting the idea that mid-LPFC language-preferring regions only existed in a subset of individuals (Figure 7F). Similar effects were seen for default B variants in 2/10 individuals (Supplemental Figure S19).

These findings provide functional evidence supporting the highly interdigitated organization of the LPFC and challenge models suggesting gradual, continuous functional gradients. These cases also vividly illustrate how averaging across individuals can produce misleading impressions of blurred boundaries between functionally specialized and domain-general control regions due to their variable location across individuals.

### Cognitive control demands recruit the dorsal attention, frontoparietal, and cingulo-opercular networks with strong activations at their borders

Having observed that some LPFC networks were preferentially activated by domain-specific non-control processes, we next examined how individual-specific LPFC networks were engaged by cognitive control demands in a wide variety of contexts. We used eight well-validated tasks spanning multiple stimulus modalities, domains, and control demands (N-back: Visual and Auditory, Memory Span: Spatial and Verbal, Multi-Source Interference: Numeric and Verbal, Attention: Visual and Auditory). Each task map was calculated as the difference between high-demand and low-demand conditions (N-back: 2-back versus 0-back, Memory Span: 8 items versus 4 items, Multi-Source Interference: conflict versus no-conflict, Attention: sustained attention versus sensorimotor control).

As shown in Figure 8A, the eight cognitive control tasks frequently engaged the same three networks—the frontoparietal, dorsal attention, and cingulo-opercular networks (corrected p < 0.05 dorsal attention: 8/8 tasks; frontoparietal: 5/8 tasks; cingulo-opercular: 4/8 tasks). Other networks, including salience/parietal memory (2/8 tasks) and language (3/8 tasks), were engaged in specific task contexts (see Supplemental Table S3 for full statistics). For example, the language network was activated during verbal working memory, auditory working memory, and auditory attention tasks, all of which involved additional reading or listening in the high-versus low-demand conditions. These network-level preferences were replicated using an alternative parcellation (Yeo-17), yielding largely comparable results, aside from subtle distinctions between FP-A (activated in 8/8 tasks) and FP-B (activated in 4/8 tasks; Supplemental Figure S20).

Consistent with observations of domain-specific demands, cognitive control activations were highly individual-specific. As quantified in Figure 8B and illustrated for three individuals in Figure 8C, LPFC activation maps showed only modest similarity across individuals performing the same task (cross-individual r = 0.27 ± 0.03). As shown in Supplemental Figure S21, interindividual differences were larger when considered at the vertex-level versus the network-level, suggesting they were more driven by interindividual variability in network topography versus interindividual differences in network engagement.

In contrast, within individuals, activation patterns across the eight control tasks were strikingly similar (r = 0.58 ± 0.03, Figure 8B, Supplemental Figure S22), suggesting a common control-related activation pattern in the LPFC. This was supported by several comparisons: within-individual similarity for different control tasks was significantly greater than similarity of the same task across individuals (unpaired two-sample t-test t(16) = 6.94, p < 0.00001, Supplemental Figure S22) and greater than the similarity of cognitive control activations to domain-specific activations within individuals (r = -0.02 ± 0.02, paired t-test t(9) = 27.2, p < 0.00001, Supplemental Figure S22). Further, unlike control task activations, different domain-specific activations showed little similarity to each other within individuals (r = 0.00 ± 0.03, paired t-test t(9) = 11.4, p < 0.00001, Supplemental Figure S22). The similarity of control activations within individuals is visible for three example individuals in Figure 8C, where a consistent set of distributed regions (marked with arrows) tended to be active across all 8 control tasks, despite large task-specific differences in stimuli and control demands (see Supplemental Figures S23 and S24 for all individual cognitive control maps and Supplemental Figure S25 for composite control maps combining control activations for each individual).

Notably, unlike LPFC activations for theory of mind, episodic projection, and language demands, which tended to fill the borders of a single network, cognitive control activations often appeared to sit at the borders between FP, DAN and CO. This is shown for three individuals in Figure 8D. To quantify this tendency, we measured the geodesic distance between active regions and the nearest other control network (e.g., if an FP vertex, the distance to the nearest CO or DAN vertex) for all individuals. We found that high-activation vertices (z > 2) were significantly closer to other control network borders than were lower-activation vertices (z < 2; Figure 8E, p < 0.001 across all 8 tasks). This effect was replicated across multiple z-thresholds (Supplemental Figure S26). To test that this border activation effect was specific to cognitive control, we quantified the distance of language, theory of mind, and episodic projection activations to network borders. We found a reverse trend - more active vertices tended to be further from network borders than less active vertices (Supplemental Figure S27), consistent with our observation that domain-specific activations appeared within the target network rather than near its borders.

Finally, to determine whether the FP, CO, and DAN border vertices that were co-active across control tasks could constitute a distinct, previously un-identified, network, we evaluated whether task co-active vertices from different control networks exhibited higher resting-state functional connectivity than not co-active vertices within the same network (Supplemental Figure S28). We found that functional connectivity was always highest within network, regardless of whether those regions were commonly co-activated during cognitive control demands (within-network not co-active FC > across-network co-active FC, paired t-test t(9) = 14.9, p < 0.000001). This suggests that these cognitive control co-active vertices do not form an independent network. However, across control networks, co-active vertices were more highly correlated with one another than not co-active vertices, providing preliminary evidence that task co-active vertices may participate in cross-network communication (across-network co-active FC > across-network not co-active FC, paired t-test t(9) = 9.2, p < 0.00001).

## DISCUSSION

Neuroimaging studies have frequently reported overlapping activations across large portions of the LPFC, supporting the idea that it is a multifunctional, flexible region (Cabeza & Nyberg, 2000; Duncan & Owen, 2000; Niendam et al., 2012). However, these reports may have missed fine-scale organizational structure, as group-averaged data can blur functionally distinct regions across individuals—particularly in the highly inter-individually variable LPFC (Dworetsky et al., 2024; Mueller et al., 2013; Seitzman et al., 2019). In this study, we collected a new precision fMRI dataset (PAN) with both resting-state and task data to examine the fine-scale organization of LPFC networks in 10 deeply sampled individuals. We found that LPFC network architecture is systematically different at the individual level compared to group averaged data: while group-level data show large, contiguous swaths of networks, individual data reveal dense interdigitation of distinct networks. This difference was most striking in anterior LPFC where a high-density zone of association networks was consistently observed across individuals but was not present in group-level data. We also validated that distinct LPFC network regions participate in distinct cognitive processes using task fMRI. This echoed prior findings that some LPFC networks are functionally specialized for domain-specific processes (e.g., LANG, DN-B, DN-A), while others support cognitive control across diverse task contexts (e.g., FP, DAN, CO). Below, we explore the implications of this organizational structure for prior and future work.

### Within-individual analysis reveals interdigitated functionally specialized and domain-general control networks in the LPFC

Our findings build on a growing body of work that has identified functionally specialized regions in the LPFC using methods that preserve individual-specific organization (Assem et al., 2025; Braga et al., 2020; DiNicola et al., 2020; Fedorenko et al., 2011, 2012; Michalka et al., 2015). In several cases, these studies showed — as we do here — that functionally specialized regions are distinct from, but often adjacent to, domain-general control regions (Assem et al., 2025; DiNicola et al., 2023; Fedorenko et al., 2012; Noyce et al., 2017). Here, we extend this work with a more comprehensive description of the patchy and interdigitated nature of LPFC organization, delineating fine-scale individual networks and dissociating their functional roles across domain-specific and domain-general control demands. We propose that fine-scale interdigitation may reflect a general organizational principle of the LPFC and suggest that further high-resolution, within-individual studies (e.g., Allen et al., 2022; Braga et al., 2020) will uncover additional examples of functional specialization.

Conceptually, this interdigitated organization contrasts with influential unitary and gradient-based accounts of LPFC organization. Such accounts emphasize smooth functional transitions along axes of control-related abstraction (Badre, 2008; Koechlin et al., 2003; Nee & D’Esposito, 2016), informational domain (Abdallah et al., 2022; Blumenfeld et al., 2013; Rahm et al., 2013) and a principal gradient of representational abstraction (Margulies et al., 2016). In contrast, we observed sharp boundaries in both task activation and functional connectivity throughout the individual LPFC, often with multiple noncontiguous patches of the same network appearing in different LPFC subregions. This was observed most dramatically in a subset of individuals where language and default B regions were idiosyncratically embedded within the canonical frontoparietal mid-LPFC region (Figure 7, Supplemental Figure S19). In these cases, functional preference shifted abruptly and in individual-specific ways. These cases highlight how group averaging can obscure sharp functional boundaries and give the illusion of broad multifunctionality, or smooth gradients. Consistent with recent theoretical arguments, (Petersen et al., 2024), we suggest that while gradients may emerge when data are smoothed—either analytically or through group averaging—they are substantially reduced in high-resolution, within-individual analyses.

### Functional implications of network fragmentation and interdigitation in the LPFC

As discussed above, individual-level analyses revealed a densely interdigitated organization in the LPFC, with high association network density, especially in the anterior LPFC (Figures 1,2). We also frequently observed fragmentation of individual network regions into multiple spatially distinct patches within the LPFC. These organizational principles were not visible in group-averaged data and have not been comprehensively described in canonical network models (e.g., Power et al., 2011; Yeo et al., 2011). The functional role of this organizational structure is not known, here we explore several different possible functional implications of a dense and fragmented architecture in the LPFC.

One possibility, for which we introduced initial evidence in this manuscript, is that network fragmentation and interdigitation may increase opportunities for information integration at the borders between networks. Several influential theories propose that cognitive control emerges from coordination between multiple large-scale networks (Assem et al., 2024; Cole et al., 2013; Dosenbach et al., 2008; Petersen & Sporns, 2015). Consistent with these ideas, we found that LPFC regions belonging to the dorsal attention, frontoparietal, and cingulo-opercular networks were frequently recruited across a range of cognitive control tasks (Figure 8A). Using within-individual network maps, it was possible to observe that these networks frequently bordered one another in individual-specific topographies (Figure 1A) and that cognitive control task activations frequently peaked near the borders between them (Figure 8D). This result echoes a recent report showing that cognitive control-related activations tend to be localized near the borders between “core” and “non-core” multiple demand regions (Assem et al., 2024) with activations shifting depending on task-specific demands. It also aligns with spatial descriptions of “connector hubs” — regions exhibiting strong functional connectivity to multiple distinct networks — which are often found near network borders or “articulation points.” (Power et al., 2013, Figure 7; Gratton et al., 2018 Figure 3). Importantly, while prior group-level findings could have been driven by individual variability and smoothing, the current findings provide corroboration of the importance of border regions within individuals. Further, we note that anatomical tracing studies in nonhuman primates have shown strong short-range connectivity across borders between distinct cytoarchitectonic areas in visual cortex (Kennedy & Bullier, 1985; Rockland & Lund, 1983), and LPFC (Barbas & Pandya, 1989; Petrides & Pandya, 2002), providing a plausible structural basis for cross-network interactions. To test this hypothesis, future work which directly manipulates cross-network integration demands will be important.

A second possibility is that the fragmentation of LPFC networks into separate small regions represents functional distinctions between those regions rather than functional utility specifically at the borders between them. While over the past decade, imaging neuroscience has shifted from a localist to a more distributed network-oriented perspective (Bassett & Sporns, 2017; Petersen & Sporns, 2015; Sporns, 2013), questions remain regarding how spatially distinct nodes within a network contribute to that network’s function. Classic models of language processing propose distinct roles for different subcomponents of a distributed language system—such as comprehension in posterior STG and production in inferior frontal gyrus (Geschwind, 1970; Hickok & Poeppel, 2007). Similarly, a seminal account of the distributed spatial attention network identified specialized roles for posterior parietal cortex (spatial location), frontal eye fields (top-down control of saccades), and cingulate cortex (motivational relevance) (Mesulam, 1981, 1990). It could likewise be the case that the multiple patches of a single network in LPFC support different functional roles. While the present study focused on network-level task activity, future work could examine specific parcels to determine whether spatially distinct regions within the same network exhibit forms of functional specialization.

A third, not mutually exclusive, possibility is that the fragmented organization of the LPFC observed here reflects a byproduct of the evolution and development of association networks in this region. A recent framework (the Expansion-Fractionation-Specialization hypothesis) by DiNicola and colleagues proposed that the evolutionary expansion of association cortex in humans enabled its fractionation into multiple parallel segregated systems specialized for different cognitive functions (DiNicola & Buckner, 2021). This framework further proposed that a similar process of segregation and specialization within association cortex occurs during the human lifespan. The lateral prefrontal cortex is notable for both its pronounced evolutionary expansion (Krubitzer 2007) and its particularly protracted developmental trajectory (Gogtay et al., 2004). These characteristics may make the LPFC more particularly likely to show high degrees of fragmentation and interdigitation, though future work using developmental data are required to evaluate this directly.

In summary, our findings provide initial evidence suggesting that border regions may play a role in control-related activity. However, more definitive interpretation will require future experiments that directly probe functional specialization, cross-network integration, and developmental changes in prefrontal cortex organization to distinguish this possibility from the alternative hypotheses described above.

### Anterior LPFC density and relationship to its special LPFC properties

From an integration perspective, regions where many distinct networks converge may play a unique role in cognition. In this study, we identified a consistent high-density zone in the anterior LPFC where many association networks were frequently represented. We speculate that such regions could have unique functions – perhaps acting as “diverse club” hubs that facilitate communication across many association networks (Bertolero et al., 2017, 2018) or supporting conjunctive coding - integrating stimulus, context, and response at the intersection of multiple networks (Badre, 2025; Fusi et al., 2016; Rigotti et al., 2013). Such ideas are especially compelling given early studies identifying the anterior LPFC at the apex of a cognitive control hierarchy, specialized for highly abstract or integrative functions (Badre, 2008; Koechlin et al., 2003). It is possible that highly abstract or complex cognitive functions may require more cross-network interactions. In this way, we speculate that studies of network density could reconcile with gradient frameworks.

Further, the anterior LPFC has been identified as the effective site for transcranial magnetic stimulation (TMS) in major depression (Siddiqi et al., 2020). One possible consequence of applying stimulation here is the potential to influence multiple networks simultaneously. While most studies of the LPFC TMS target have focused on its connectivity with the subgenual cingulate (Fox et al., 2012), we propose that future studies may examine how the uniquely dense topography may contribute to treatment effects. Dense interdigitation may also help explain the broad cognitive deficits observed following LPFC lesions, which are often nonspecific and span multiple domains (Luria, 1966; Roca et al., 2010; Shallice & Burgess, 1991; though see Stuss, 2011). Lesions in this densely packed area may more easily cross network boundaries, resulting in less selective functional deficits.

These findings offer a new framework for considering the LPFC’s functional role in comparison to other brain regions. The LPFC has been historically considered a special area with strong integrative qualities distinct from posterior brain regions (Miller, 2000). However, later network analyses revealed that the LPFC contains the same association networks also present in the parietal, temporal, and cingulate cortices (Power et al., 2011; Yeo et al., 2011), raising the possibility that unique qualities ascribed to the LPFC may be better attributed to specific highly integrative networks (Cole & Schneider, 2007). Here, we propose a unifying hypothesis: the LPFC is not unique in network identity but rather in network density.

### Precision fMRI reveals common cortical features obscured by group averaging

This study uncovered fine-scale features of LPFC network organization that were consistent across individuals but not visible in group-averaged data. Notably, we identified a reproducible DAN–CO–LANG motif in anterior LPFC (Figure 3) which was absent from group-level data. In contrast, a prominent feature of the group-averaged map—a continuous frontoparietal region in dorsolateral PFC (Figure 1)—was present in only 1 of 10 individuals. This highlights a key limitation of group averaging: it can produce features that do not reflect the organization of any individual.

These results parallel recent precision fMRI studies that have revealed similarly fine-scale motifs obscured in group data, including parallel default networks (Braga & Buckner, 2017; DiNicola et al., 2020), a novel somato-cognitive-action network (Gordon et al., 2023), and a unified salience/parietal memory network (Kwon et al., 2025) among others.

Although the three-network motif identified here has not been the central focus of prior work, converging evidence supports its existence. Assem et al., 2024 reported strong connectivity between the dorsal attention network and a rostral LPFC region in group-averaged data and hypothesized the existence of a dorsal attention region in that location (their Supplemental Figure 4). Braga et al., 2020 identified an anterior LPFC region assigned to the language network across individuals, not captured in group-level atlases. Du et al., 2024 observed a similar motif to the one we observed in multiple individuals (Figures 2, 5, 14–16), though it was not a focal point of their study and was detected using a different parcellation method (MS-HBM). We also found this motif repeated in other cortical zones—including pre-SMA, TPJ, and caudal LPFC—consistent with prior work showing that networks often border the same neighbors in multiple locations (Braga & Buckner, 2017; DiNicola et al., 2020). This repeating spatial structure aligns with classic proposals of parallel, functionally specialized processing streams (Goldman-Rakic, 1988).

Identifying common fine-scale features is essential for grounding theoretical models of brain organization. In the present study, detecting small anterior regions of the dorsal attention and language networks in individuals was critical for revealing the high association network density region in the anterior LPFC and for localizing the CO region as its spatial anchor (Figure 4). Further, while influential models propose that an abstraction hierarchy in the LPFC may map to a sequence of distinct LPFC networks (Badre & Nee, 2018; Nee, 2021), these models do not account for the repeated interdigitation of network patches within the LPFC. Future models can incorporate the patchy, interdigitated organization revealed by precision fMRI.

### Precision fMRI can identify individual-specific cortical features

In addition to common cortical features, precision fMRI enables the identification of idiosyncratic network regions that do not conform to a shared cortical template and are present in only a subset, or even a single individual (Dworetsky et al., 2024; Glasser et al., 2016; Gordon et al., 2017; Laumann et al., 2015; Seitzman et al., 2019). In this study, we characterized several such regions of the language and default B networks embedded in the mid-LPFC frontoparietal region in a subset of individuals. We validated these regions using task activations and observed they were particularly strong examples of adjacent specialized and domain-general control activity in the LPFC.

Prior work has differentiated distinct types of idiosyncratic features: “ectopic intrusions” --islands of altered connectivity and function, and “border shifts”– local expansions or contractions of commonly shared network regions. It was found that ectopic intrusions are most prevalent in the LPFC while border shifts are most prevalent in the TPJ (Seitzman et al., 2019, Dworetsky et al., 2024). In this study, we characterized both ectopic (both DN-B variants, LANG variants PAN02 and PAN09) and border shifts (LANG variants PAN01 and PAN06). Important questions remain about why ectopic intrusions are so common in the LPFC and how they develop. Prior work showed that these regions are less heritable than border shifts and so may relate more to experience-dependent processes. The LPFC undergoes particularly protracted development (Gogtay et al., 2004), which may render it especially plastic to experience-dependent processes (Johnson, 2001; Petanjek et al., 2011). Future work may investigate how ectopic intrusions in LPFC arise and vary across individuals, potentially offering new insight into how life experience shapes large-scale cortical organization.

### Limitations

This study has several methodological limitations, which we have attempted to address where possible. First, in deriving individual-specific networks, we chose one specific parcellation strategy (vertex-level Infomap with a template matching manual consensus procedure following Lynch et al., 2024) based on an 18-network group prior. We selected this prior because it allowed us to test a priori hypotheses about the functional preferences of specific networks (i.e., language, default A/B) and selected vertex-level resolution to identify small network regions. However, this raises the possibility that our findings may be specific to the parcellation method or group prior. To mitigate this risk, we validated the parcellation using seed maps (Supplemental Figure S7) and replicated network composition, interdigitation, and task preference findings using an additional group atlas for template matching (Yeo 17-network atlas; Supplemental Figures S6, S10, S20).

Second, to examine the relationship between individual task activation maps and individual-specific networks, we required highly reliable individual-level task data. We therefore used block designs across all tasks, as these have been shown to produce more robust and reliable individual z-statistic maps than mixed or event-related designs (Mumford et al., 2014; Noble et al., 2021). However, block designs limit the ability to resolve temporally distinct cognitive processes within tasks—an important caveat in cognitive control paradigms, where different networks may operate at different timescales (Dosenbach et al., 2008). Future work may be able to produce temporally sensitive, individual-specific maps of these control-related dynamics.

Third, our analyses were generally constrained by our relatively small sample size (N = 10). While this is typical for precision fMRI studies (e.g., Allen et al., 2022; Braga & Buckner, 2017; DiNicola et al., 2023; Du et al., 2024; Gordon et al., 2017; Kwon et al., 2025; Salvo et al., 2025), it limits our ability to assess the prevalence of these patterns in the broader population or to detect variability not represented in our sample. To mitigate this limitation, we replicated our primary findings in the independent Natural Scenes Dataset (N = 6; Allen et al., 2022) which was collected at 7T with substantially different imaging parameters. Future work can further extend this to assess the prevalence of the specific network variants we identified in larger populations as well as assess their possible relationships with behavior.

Finally, all estimates of network topography and task activation were limited by the inherent smoothness of fMRI data. To reduce this risk, we used small voxel sizes (2.5 mm³) and minimal smoothing (1 mm Gaussian kernel). This high-resolution approach enabled detection of small, spatially reliable network regions critical for identifying LPFC motifs which were not visible in group-averaged data (Figure 3, see PAN05 dorsal attention network), while still preserving reliable measures within individuals. We further mitigated this risk by replicating network findings in a high resolution 7T dataset (Natural Scenes Dataset, 1.8 mm^3^ voxels, Allen et al., 2022). In the future, it will be important to replicate task results at 7T as well. In particular, the border activations observed between control-related networks (Figure 8D) could partly reflect signal spillover between adjacent networks. However, this account does not fully explain our findings. Some control-network regions showed minimal or no activation during control tasks, inconsistent with border peaks resulting from widespread activation combined with smoothing. Nevertheless, replication of task-activation effects with higher-resolution methods will be important to confirm these findings and evaluate residual smoothing artifacts.

## Conclusion

In this study, we collected a new precision fMRI dataset (PAN) to map the fine-scale network organization of the lateral prefrontal cortex (LPFC) in 10 deeply sampled individuals. While the LPFC is often portrayed as a broad, multifunctional region, our data revealed a patchy, interdigitated architecture in which distinct networks support domain-general control and domain-specific processing. We identified both conserved motifs—such as a consistent high-density zone in anterior LPFC—and individual-specific variants. These features were validated using both task and resting-state data and replicated in an independent dataset.

Together, these findings offer a more detailed picture of LPFC organization. They highlight the value of individual-specific approaches for revealing fine-scale functional architecture and suggest that spatial patterns of network interdigitation may play a role in how the LPFC supports flexible and complex cognition.

## STAR METHODS

## KEY RESOURCES TABLE

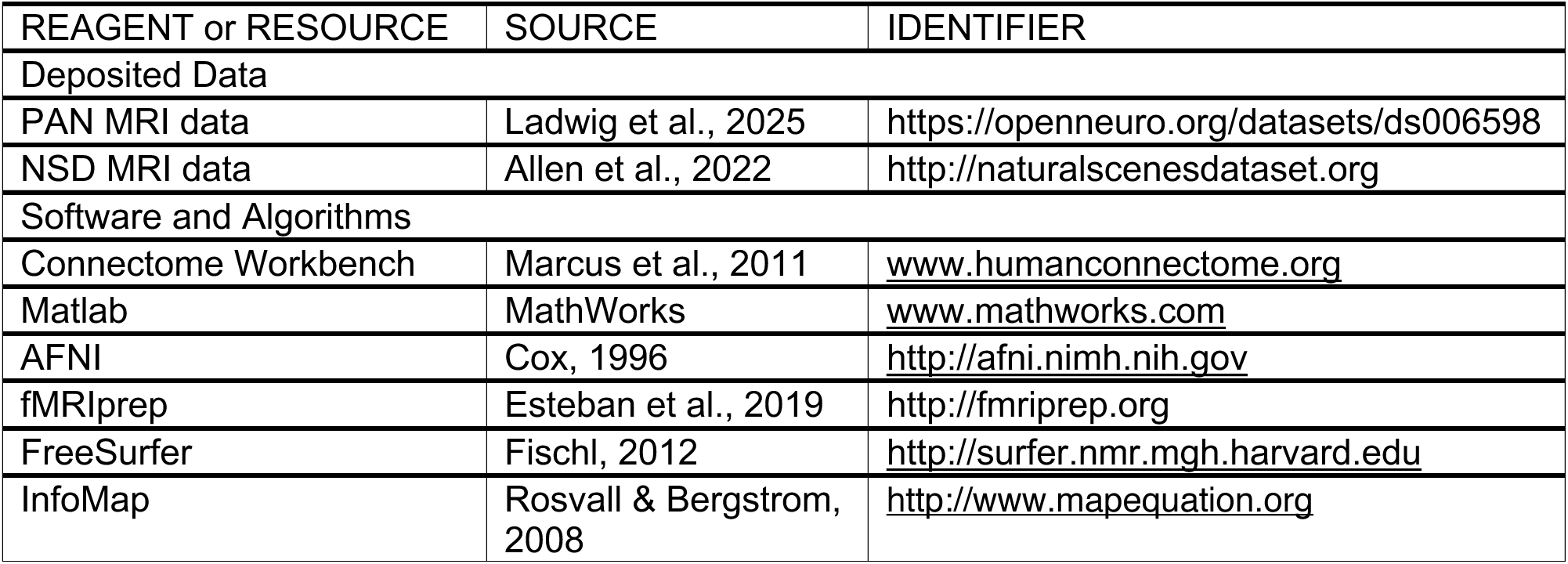

## EXPERIMENTAL MODEL AND STUDY PARTICIPANT DETAILS

### Participants

Our primary analyses rely on the PAN dataset. In this dataset, ten paid participants (ages 18-33 years, mean age = 26.4, SD = 4.8, 5F) completed one behavioral training session and 7-10 fMRI sessions. Each fMRI session lasted ∼1.5 hours, comprising 8-11 task runs (5-10 minutes each) and 4-5 resting-state runs (5 minutes each). Participants were required to have corrected-to-normal vision, be native/fluent English speakers, be right-handed, and have no history of neurological disorders. All participants provided written informed consent, and the study was approved by the Northwestern University and Florida State Institutional Review Boards.

Our replication analyses rely on the publicly available Natural Scenes Dataset (NSD; Allen et al., 2022). The preprocessed NSD data was generously shared by the Braga Lab. The processing methods are described here in brief, with additional details available in Kwon et al., 2025. Two individuals out of eight (S5, S8) were excluded due to head motion (ages 23-30 years, mean age = 26.8 ± 2.8 years). Individuals completed 30-40 weekly fMRI sessions (approximately 2.0 hours of resting-state and 38.5 hours of task fMRI data per individual); only the resting-state data was used in this study.

## METHOD DETAILS

### MRI Acquisition

PAN data for one individual (PAN01) was acquired at the Northwestern University Center for Translational Imaging (CTI) on a 3T Siemens Prisma MRI scanner with a 64-channel head coil (Siemens Healthcare, Erlangen, Germany). PAN data for the remaining 9 individuals was acquired at the Florida State University MRI Facility, also on a 3T Siemens Prisma MRI scanner with a 64-channel head coil. A multi-echo multiband sequence was used for all functional runs (voxel size: isotropic 2.4mm, TR = 1.355s, TE = 12.0ms, 32.4ms, 52.0ms, 71.6ms, 91.2ms, flip angle = 59°, multi-band factor = 6, in-plane acceleration factor = 2, AP phase encoding direction, FOV=216mm, Slices = 54, see Lynch et al., 2020). Two structural (T1) scans were acquired per individual on separate days [voxel size: isotropic 1mm, TR = 2.3s, TE = 1.86ms, 3.78ms, TI = 1180ms, flip angle = 7°, FOV = 256mm, Slices = 208]. A gradient-echo fieldmap with the same geometric parameters as the functional sequence was acquired in each fMRI session. Motion was monitored in real-time using Framewise Integrated Real-time MRI Monitoring (Dosenbach et al., 2017) and individuals’ eyes were video monitored for signs of sleepiness.

NSD data was collected at the Center for Magnetic Resonance Research at the University of Minnesota using a 7T Siemens Magnetom MR scanner. A single-echo multiband sequence was used for functional runs (1.8-mm isotropic resolution, whole-brain coverage, TR = 1,600 ms, TE = 22.0 ms, flip angle 62°, FOV = 216 mm (FE) × 216 mm (PE), slice thickness 1.8 mm, slice gap 0 mm, matrix size 120 × 120, echo spacing 0.66 ms, bandwidth 1,736 Hz per pixel, partial Fourier 7/8, iPAT 2, multiband slice acceleration factor = 3, and 84 slices acquired in the axial plane). Dual-echo fieldmaps were collected.

### Task Paradigms

All individuals completed at least 30 minutes per demand across 11 different cognitive demands targeting language, theory of mind, episodic projection, and cognitive control. Task paradigms were selected from the literature based on their ability to elicit reliable LPFC activity within single individuals and their suitability for repeat testing (i.e., being resilient to habituation or allowing for easy generation of novel stimuli). Tasks were adapted with minimal modifications from the original publications, with any relevant changes described below. In several cases, domain-level (theory of mind, episodic projection, and language) activation maps were derived by averaging activation maps from multiple tasks together (ex: retrospection and prospection for episodic projection). This was done to directly replicate prior literature that created domain-level composites using these specific tasks and showed they yield similar activation patterns (DiNicola et al., 2020). Here we extend this approach by examining these domain-level activations specifically within LPFC network regions. All tasks were implemented using Psychtoolbox-3.

### Language

Two language tasks were administered: a visual language task and an auditory language task (described below). Recent work has shown that visual and auditory language tasks elicit highly similar activation maps which selectively engage the transmodal language network (Salvo et al., 2025). These two tasks were averaged to generate individual-level language activation maps.

#### Visual Language Task

The visual language task was adapted from Fedorenko et al., 2011 using publicly available code and stimuli (https://www.evlab.mit.edu/resources-all/download-localizer-tasks). Individuals passively read either real sentences (Sentences condition) or pronounceable nonword sequences (Nonword condition), presented one (non)word at a time. Each experimental block lasted 18 seconds and included three trials. Each trial lasted 6 seconds, beginning and ending with 100 ms of blank screen. During the trial, 12 (non)words were presented every 450 ms, followed by a 400 ms response cue prompting participants to press a button to mark the end of the trial.

Individuals completed 16 experimental blocks, intermixed with five 14-second fixation blocks placed at the start of each run and after every four experimental blocks. Each person completed at least six runs of the visual language task (5 minutes 58 seconds per run; 35 minutes 48 seconds total). Condition order was counterbalanced across runs and held constant across individuals. Evidence suggests that contrasting Sentence > Nonwords blocks reliably identifies high-level language processing regions, including in the LPFC (Fedorenko et al., 2011, 2012).

#### Auditory Language Task

The auditory language task was adapted from Scott et al., 2017 using publicly available code and stimuli (https://www.evlab.mit.edu/resources-all/download-localizer-tasks), along with additional stimuli created in-house. Individuals listened to 18-second clips of either intact speech (Intact condition) or acoustically degraded speech (Degraded condition). Degraded clips retained auditory properties of speech but were not intelligible. To expand the stimulus set beyond the 32 publicly available clips, we created 64 additional stimuli by extracting 18-second segments from episodes of *The Moth* podcast and degrading them using the procedure described below. Each run included 16 experimental blocks (one 18-second audio clip per block) and five 14-second fixation blocks, placed at the beginning of each run and after every four experimental blocks. All individuals completed six runs of the auditory language task (5 minutes 58 seconds per run; 35 minutes 48 seconds total). Condition order was counterbalanced across runs and held constant across individuals. Evidence suggests that contrasting Intact > Degraded blocks reliably identifies high-level language-processing regions, including regions in the LPFC (Salvo et al., 2025; Scott et al., 2017).

##### Auditory Degradation

To degrade novel audio clips, we followed the procedure outlined in Scott et al. (2017) using their publicly available code. Briefly, a low-pass version of the intact clip was created with a passband cutoff at 500 Hz. A noise track was generated by temporally scrambling the intact audio and modulating it by the original amplitude envelope to preserve naturalistic intensity variation. This noise signal was then bandpass filtered between 8000–10,000 Hz and added to the low-pass version of the clip. All degraded clips were manually checked to ensure they were unintelligible while preserving general acoustic similarity to the original speech.

### Episodic Projection

The episodic projection task was adapted from Andrews-Hanna et al., 2010 and DiNicola et al., 2020. Code and stimuli for the task were generously provided by DiNicola and colleagues. This task probes remembering and prospection demands by asking participants to respond to scenarios regarding past, future, or present events. While participants were presented with scenarios in six possible categories (Past Self, Present Self, Future Self, Past Non-Self, Present Non-Self, Future Non-Self), only self-related scenarios were used in this manuscript. Following DiNicola et al., 2020, a general episodic projection contrast was derived by averaging together contrasts targeting Retrospection (Past Self > Present Self) and Prospection (Future Self > Present Self). Episodic projection runs were 624 seconds long, consisting of 30 20-second experimental blocks (5 per category) and 12-second fixation blocks beginning and ending the run. Each experimental block consisted of a 5-second fixation period, a 10-second trial period, and another 5-second fixation period. During the trial period, participants viewed a scenario and related question and were asked to select from three possible responses with a button press. The order of scenarios was randomized and kept consistent across participants. All participants completed six runs of the episodic projection task (10 minutes 24 seconds each; 62 minutes 24 seconds total). Evidence suggests that the general episodic projection contrast dissociates regions of default A from regions of default B, including regions in the LPFC (DiNicola et al., 2020).

### Theory of Mind

Two theory of mind tasks were administered: False Belief and Emotional/Physical Pain, both replicated from DiNicola et al. (2020). As in prior literature, high-level theory of mind processing contrast was created by averaging the contrasts from these two tasks (Belief > Photo and Emotional Pain > Physical Pain) (DiNicola et al., 2020; Du et al., 2024). Evidence suggests that this contrast dissociates regions of default B from regions of default A, including regions in the LPFC (DiNicola et al., 2020).

#### False Belief/Photo

The False Belief task was generously shared by the Braga Lab, adapted from publicly available materials developed by Dodell-Feder et al., 2011 (https://saxelab.mit.edu/use-our-efficient-false-belief-localizer/). In this task, participants read short stories in which either a character held a potentially false belief (False Belief condition) or an object (e.g., a photograph or map) contained potentially false information (False Photo condition). They then answered a true/false question via button press. Each run was 324 seconds long, with ten 30-second trials (five per condition) and 12-second fixation periods at the beginning and end. Each trial included a 10-second story, 5-second response, and 15-second fixation. Story order was randomized and consistent across participants. All individuals completed four runs (5 minutes 24 seconds each; 21 minutes 36 seconds total).

#### Emotional/Physical Pain

The Emotional/Physical Pain task was generously shared by the Braga Lab, adapted from publicly available materials developed by Jacoby et al., 2016 (https://saxelab.mit.edu/theory-mind-and-pain-matrix-localizer-narratives/). In this task, participants read stories describing a character experiencing either emotional or physical pain and rated the character’s pain on a 1–4 scale via button press. Each run lasted 324 seconds, with ten 30-second trials (five per condition) and 12-second fixation periods at the beginning and end. Each trial included a 10-second story, 5-second response, and 15-second fixation. Story order was randomized and consistent across participants. All individuals completed four runs (5 minutes 24 seconds each; 21 minutes 36 seconds total).

### Cognitive Control

We report on eight contrasts derived from six distinct cognitive control tasks. Four tasks (MSIT, VMSIT, Spatial Memory Span, Verbal Memory Span) were adapted from Fedorenko et al. (2013), which identified “domain-general control” regions based on greater activation for “hard” versus “easy” conditions across diverse tasks. Two additional tasks—Visual vs. Auditory Attention and Visual vs. Auditory N-back—were adapted from Noyce et al., 2017 and Michalka et al., 2015. We hypothesized these control tasks would show distinct recruitment from the language, episodic projection, and theory of mind contrasts described above.

#### Visual and Auditory Attention

The visual and auditory sustained spatial attention task was adapted from Michalka et al., 2015 with minimal modification. Participants were simultaneously presented with four rapid serial streams of stimuli (two auditory, two visual), each containing either numeric or alphabetical characters. They were instructed to monitor one stream for target digits (1–4) while ignoring distractor letters in that stream (‘A’, ‘F’, ‘G’, ‘H’, ‘J’, ‘K’, ‘L’, ‘M’, ‘N’, ‘P’, ‘R’, ‘X’, ‘Y’) and the other three streams, which contained only digits (1–9, excluding 7). Each visual stream was flanked by three additional digit-only distractor streams. Participants responded via button press (1–4) to target digits in the attended stream, which occurred three times per block. In sensorimotor control (“passive”) blocks, participants viewed the same stream layout, but all streams contained only digits (1–9), and participants pressed each response button once at a relaxed pace. Fixation control blocks showed a white cross on a black screen. Each run lasted 363.2 seconds and included an 8-second initial and 12-second final fixation, ten 26-second experimental blocks (two per stream type and two passive), two 26-second fixation blocks, and 2.6-second audiovisual cue periods announcing upcoming block types. Each experimental block contained 40 stimuli, presented audiovisually for 300 ms with a 350 ms inter-stimulus interval. Block order was counterbalanced across runs and consistent across individuals. A Visual Attention contrast was defined as Visual > Passive; an Auditory Attention contrast was defined as Auditory > Passive. All individuals completed at least six runs (7 minutes 3.2 seconds each; 36 minutes 19 seconds total).

#### Visual and Auditory N-Back

The visual and auditory working memory task was adapted from Noyce et al., 2017. Participants performed either a 2-back working memory task or a 0-back sensorimotor control task on visual (faces) or auditory (animal sounds) stimuli, presented in separate blocks. Visual stimuli were black-and-white photographs of young adult faces from the Chicago Face Database (Ma et al., 2015) with male and female faces presented in separate blocks to increase difficulty. Auditory stimuli consisted of cat and dog sounds, also presented in separate blocks. Each run lasted 372 seconds and included an initial 8-second fixation period, a mid-run 8-second fixation break (after four blocks), and a final 12-second fixation. Runs included eight 43-second blocks: one per combination of stimulus category (Male Faces, Female Faces, Cat Sounds, Dog Sounds) and demand level (2-back or sensorimotor). Each block began with a 3-second cue indicating the upcoming category, followed by a 40-second trial period. Visual stimuli were displayed for 1000 ms with a 250 ms ISI; auditory stimuli lasted ∼500 ms and were spaced 1250 ms apart to match timing. In 2-back blocks, participants responded to each stimulus as “new” or a “2-back repeat” (25% of trials). In sensorimotor blocks, no repeats occurred, and participants made random button presses to each stimulus. Block order was counterbalanced across runs and consistent across individuals. A Visual n-back contrast was defined as Visual 2-back > Visual Sensorimotor; an Auditory n-back contrast as Auditory 2-back > Auditory Sensorimotor. All participants completed at least six runs (6 minutes 12 seconds each; 37 minutes 12 seconds total).

#### Spatial Memory Span

The spatial memory span task was adapted from Fedorenko et al., 2011. Participants viewed a 3×4 grid and were sequentially shown spatial locations to remember, with either four or eight total locations depending on difficulty (easy vs. hard conditions). At the end of each trial, they selected the correct grid from two options via button press; the incorrect grid contained one or two wrong locations. In easy blocks, locations were presented one at a time (four total); in hard blocks, locations were shown in pairs (eight total). Each run lasted 436 seconds and included ten 34-second experimental blocks (five easy, five hard) and six 16-second fixation blocks. Each experimental block contained four trials, with each trial consisting of a 0.5-second fixation, four 1-second stimulus presentations, and a 4-second response window. The order of easy and hard blocks was counterbalanced across runs and kept consistent across individuals. All participants completed at least four runs (7 minutes 52 seconds each; 31 minutes 28 seconds total).

#### Verbal Memory Span

The verbal memory span task was adapted from Fedorenko et al., 2011. Participants were sequentially presented with spoken digit names and asked to remember either four or eight digits depending on condition difficulty (easy vs. hard). At the end of each trial, they selected the correct sequence from two options via button press; incorrect options contained one or two wrong digits. In hard blocks, digit names were presented in pairs (eight total); in easy blocks, they were presented one at a time (four total). Each run lasted 436 seconds and included ten 34-second experimental blocks (five easy, five hard) and six 16-second fixation blocks. Each block contained four trials, with each trial consisting of a 0.5-second fixation, four 1-second digit presentations, and a 4-second response window. Block order was counterbalanced across runs and consistent across individuals. All participants completed at least four runs (7 minutes 52 seconds each; 31 minutes 28 seconds total).

#### Numeric Multi-Source Interference Task (MSIT)

The numeric Multi-Source Interference Task was adapted from Fedorenko et al., 2011, which was originally adapted from Bush et al., 2003. Participants were shown digit triplets (e.g., “123”) and asked to respond based on the identity—rather than the position—of the unique (non-repeated) digit. In the easy condition, the identity of the target digit matched its position, and the distractors were not valid response options (e.g., “003”). In the hard condition, the identity of the target digit did not match its position, and the distractors were valid response options (e.g., “131”). Each run lasted 396 seconds and included two 30-second fixation blocks (at the beginning and end) and eight 42-second experimental blocks. Each experimental block contained 24 trials with a 1.5-second stimulus presentation and a 0.25-second interstimulus interval (ISI). The order of hard and easy blocks was counterbalanced across runs and consistent across participants. All individuals completed at least four runs (6 minutes 36 seconds each; 26 minutes 24 seconds total).

#### Verbal Multi-Source Interference Task (VMSIT)

The Verbal Multi-Source Interference Task (VMSIT) was adapted from Fedorenko et al., 2011, which was originally adapted from Bush et al., 2003. Participants were shown triplets of words (e.g., “none,” “left,” “middle,” “right”) and asked to respond based on the meaning of the unique (non-repeated) word, rather than its spatial position. In the easy condition, the position of the target word matched its meaning, and the distractor words were not valid response options (e.g., “none none right”). In the hard condition, the target word’s position conflicted with its meaning, and the distractor word(s) were valid response options (e.g., “left right left”). Each run lasted 396 seconds and included two 30-second fixation blocks (at the beginning and end) and eight 42-second experimental blocks.

Each block contained 24 trials, with 1.5-second stimulus presentation and a 0.25-second interstimulus interval (ISI). The order of easy and hard blocks was counterbalanced across runs and consistent across individuals. All participants completed at least four runs (6 minutes 36 seconds each; 26 minutes 24 seconds total).

## QUANTIFICATION AND STATISTICAL ANALYSES

### fMRI Processing

PAN fMRI data was processed using fMRIprep 23.0.2 – a standardized open-source pipeline for minimal fMRI preprocessing. This included the creation of a reference T1w, skull-stripping, head motion correction (rigid body correction using 6 parameters), segmentation of white matter, gray matter, and CSF, susceptibility distortion correction, registration to T1w, spatial normalization to MNI152NLin6Asym 2mm isotropic resolution template space, the optimal combination of echoes, and the generation of a cortical surface with FreeSurfer (Dale et al., 1999; Esteban et al., 2019). To maximize comparability with prior work, we processed multi-echo data in this study using the standardized preprocessing pipeline implemented in fMRIprep 23.0.2. This approach includes the optimal combination of echoes (OC-ME), but does not apply ME-ICA, which has been shown to further improve the reliability of individual-level functional connectivity estimates (Lynch et al., 2020). The resulting surfaces were registered into fs_LR_32k surface space as described in Glasser et al., 2013. For resting-state fixation runs, after running fMRIprep 23.0.2, data was further processed as in Power et al., 2014 to reduce the effect of artifacts on functional connectivity estimation. Data was demeaned and detrended, nuisance signals (motion parameters, white matter, gray matter, CSF, global signal) were regressed, high motion frames (fFD > 0.1mm, as in Gratton et al., 2020) were censored and their data interpolated, the residual data were band-pass filtered (0.08-0.009 Hz). After preprocessing in the volume, cortical functional data were registered to the fslr32k surface. Cortical surface and volumetric subcortical and cerebellar data were combined into CIFTI format using Connectome Workbench (D. S. Marcus et al., 2011) and data were (minimally) smoothed (Gaussian kernel, sigma = 1mm) using 2D geodesic smoothing on the surface and 3-D Euclidean smoothing for subcortical volumetric data.

NSD data was downloaded (http://naturalscenesdataset.org) after slice time correction, head motion correction, alignment across sessions and correction for EPI distortion. Additional functional connectivity processing was done including nuisance signal regression (six parameters for head motion, whole-brain, ventricular, white matter signal, and temporal derivatives) and bandpass filtering (0.01-0.1 Hz). Data was projected on to a standardized cortical surface (fs_LR_32k) and smoothed using a 1mm FWHM kernel.

### fMRI Data Quality Control

For the PAN dataset, motion-contaminated frames were identified by filtered framewise displacement (fFD) (Gratton et al., 2020). Frames with fFD > 0.1 were flagged as high motion frames. For task data, runs were excluded if > 20% of frames were flagged as high motion frames or if task performance was below chance. In total, 791 task runs were completed in this study, and 11 runs were excluded. PAN02 had 1 Auditory Attention run excluded for task performance, PAN05 had one Auditory Attention run excluded for task performance, PAN06 had one Episodic Projection run excluded for motion, PAN07 had two runs (Auditory Attention, Theory of Mind - False Belief) excluded for task performance, and one run (Visual Language) removed for motion. PAN08 had two runs (Auditory Attention, Theory of Mind) excluded for task performance, and one run (Episodic Projection) excluded for motion. PAN09 had one run (Episodic Projection) excluded for motion. PAN10 had one run (Visual Working Memory) excluded for task performance. For functional connectivity analyses using resting-state data, all runs were included but high motion frames were censored to minimize bias from motion-related artifacts (Gratton et al., 2020; Power et al., 2014b). On average across individuals, 95.5% ± 2.1% of frames were retained, leaving a minimum of 101 motion-censored minutes of data per individual (range = [101 minutes – 159 minutes], average = 118 minutes). Prior work suggests at least 40 minutes of data are necessary for precision mapping of functional networks (Gordon et al., 2017; Laumann et al., 2015), though possibly less with multi-echo data (Lynch et al., 2020).

Following Kwon et al., for the NSD data, high motion runs were identified and discarded. High motion runs had > 0.4 mm maximum framewise displacement (FD), > 2.0 mm maximum absolute motion or visible motion along with maximum FD > 0.2mm or maximum absolute motion > 1.0 mm. After discarding high motion runs, individuals retained between 6 and 35 low motion runs (NSD01: 35, NSD02: 6, NSD03: 16, NSD04: 12, NSD06: 19, NSD07: 18).

### Precision Functional Network Mapping

Individual-specific functional networks were derived using the InfoMap community detection algorithm (Rosvall & Bergstrom, 2008b), following the approach described in Lynch et al., 2024. For each participant, we built a functional connectivity matrix by correlating time series between all cortical and subcortical vertices across all resting-state runs and sessions. To reduce the influence of spatially local correlations, connections between nodes ≤ 20 mm apart were set to zero—using geodesic distance for cortical-cortical connections and Euclidean distance for cortical-subcortical ones. These matrices were thresholded at a range of densities (0.01%, 0.02%, 0.05%, 0.1%, 0.2%, 0.5%, 1%, 2%, and 5%) to retain only the strongest edges per vertex. Each thresholded matrix was input to InfoMap to identify community structure. We focused on the 0.1% threshold, as prior work has shown it produces networks with the highest size-weighted homogeneity compared to randomly rotated null models (Gordon et al., 2020; Lynch et al., 2024). At this threshold, individuals showed an average of 62.2 ± 5.0 functional communities.

Network identities were assigned to each community by matching them to a reference set of 18 independently derived functional networks (default B, default A, visual-lateral, visual-stream, visual-V1, visual-V5, frontoparietal, dorsal attention, premotor, language, salience, parietal memory, cingulo-opercular, auditory, somatomotor-hand, somatomotor-face, somatomotor-foot, somato-cognitive-action), based on both functional connectivity and spatial location. These priors were adapted from a precision mapping dataset of 37 healthy adults (Lynch et al., 2024) The priors were kept identical, with the exception that the default-dorsolateral, default-anterolateral, and default-parietal networks were combined into a single default-B network. This was done to match prior literature dissociating two distinct default networks (DiNicola et al., 2020, 2023) and to maintain consistency with prior network parcellations (Gordon et al., 2017; Power et al., 2011). A confidence score was automatically generated for each match, equally weighting functional connectivity and spatial topography. Communities were manually reviewed, and in ambiguous cases, their labels were adjusted by Z.L and verified by K.K. When a community didn’t resemble any known network—often in low-signal regions—it was labeled “noise.” On average, 3.3% ± 0.7% of cortical vertices were reassigned to a different network after review, and 1.7% ± 1.0% were labeled as noise. Following Gordon et al., 2017, we removed contiguous network patches smaller than 50 mm² and filled them in by dilating neighboring labels. This threshold was determined manually in the first two participants through an analysis of the size of regions which were replicable across split halves and was applied equally to all individuals. Although networks were estimated for subcortical structures, only cortical assignments were used in subsequent analyses. Finally, based on recent work suggesting that the salience and parietal memory networks are part of a single large-scale system (Du et al., 2024; Kwon et al., 2025) and consistent with our own seed-based maps, we combined them into one network. To ensure our results didn’t depend on a single parcellation approach, we repeated the network assignment step using the Yeo 17-network atlas (Yeo et al., 2011). Because the Yeo 17 prior lacked connectivity templates, assignments were based on spatial overlap alone. For the NSD data, the same protocol was followed but only used cortical data and retained smaller regions (40 mm^2^) due to the increased resolution of NSD data. On average across the 6 individuals, 4.9% ± 0.1% of cortical vertices were reassigned to a different network after review and 5.6% ± 1.0% labeled as noise.

### Group-Averaged Network Parcellation

Comparisons were made between individual network parcellations and a group-averaged parcellation. The primary group parcellation was defined as the mode network assignment across the 37 healthy controls used as priors here and in Lynch et al. (2024). For comparison, we also generated a group-averaged parcellation from the 10 individuals included in the main analyses. This was done by averaging their dense connectivity matrices and running them through the same precision functional mapping procedure. The resulting parcellation was highly similar to the Lynch et al. (2024) group-averaged map (Supplemental Figure S3), confirming its validity for our sample. As described above, individual network maps were also generated using the Yeo et al. (2011) 17-network parcellation as a spatial prior (Supplemental Figure S6). These were compared to the publicly available group-averaged 17 network parcellation (https://github.com/ThomasYeoLab/CBIG/tree/master/stable_projects/brain_parcellation/Yeo2011_fcMRI_clustering). For more straightforward comparison, LPFC networks in the Yeo 17-network parcellation were manually renamed to match the network names used in the current study where possible (their control A = frontoparietal A, control B = frontoparietal B, control C = parietal memory, temporal parietal = language, default C = default A, default A = default B1, default B = default B2, salience/ventral attention A = cingulo-opercular, salience/ventral attention B = salience, dorsal attention A = visual-stream, dorsal attention B = dorsal attention).

### Cortical Masks

An average LPFC mask was created and used across all individuals. This mask was generated by taking the union of Freesurfer aparc.a2009s parcellations from all 10 individuals, including the following parcels: L_G_and_S_frontomargin, L_G_and_S_transv_frontopol, L_G_front_inf-Opercular, L_G_front_inf-Triangul, L_G_front_middle, L_Lat_Fis-ant-Vertical, L_S_front_inf, L_S_front_middle, L_S_front_sup, L_S_orbital_lateral, L_S_precentral-inf-part, and L_S_precentral-sup-part.

Masks for the lateral parietal, lateral temporal, and medial prefrontal cortex were created manually using the Schaefer 300 parcellation and applied identically across all individuals.

### Split Half Functional Network Reliability

For two individuals (PAN01 and PAN02), approximately 2.5 hours of motion-censored resting-state data were collected (160 minutes and 146 minutes, respectively). These individuals were used as pilot cases to assess the reliability of the precision functional mapping approach. Their data were split into odd and even sessions (always collected on different days), and precision functional mapping was run independently on each half, starting from the construction of functional connectivity matrices as described in “Precision Functional Mapping.”

The reliability of individual LPFC networks (Supplemental Figure S4) was assessed by computing the Dice overlap for each LPFC network between the two halves. To quantify whether within-individual reliability was greater than between-individual similarity, the average dice overlap value per network was calculated over the two individuals, and a one-sided paired t-test was used to compare the overlap of networks within individuals across split halves to the overlap of networks across individuals within the same split half.

### Surface Area Analyses

For each individual, the percent of total surface area for each network was calculated using the final network estimates, masked to include only the LPFC. Surface area per network was defined as the sum of the surface area of each vertex assigned to that network (Figure 1). Vertex-wise surface areas were calculated using the Conte69 32k fs_LR template surface.

### Association Network Density

For each individual, association network density was calculated at each vertex as the number of unique association networks present within a fixed-radius neighborhood (Figure 4). This approach was adapted from Power et al., 2013, where it was termed “community density” and was calculated on all communities across different thresholds. Here, we refer to it as association network density, as the calculation was based on the final labeled network identities and limited to the 7 association networks (dorsal attention, frontoparietal, default A, default B, salience/parietal memory, cingulo-opercular, language), excluding sensory-motor processing networks (visual, auditory, somatomotor, premotor, somato-cognitive-action). For the Yeo 17-networks parcellation, 10 association networks were included (dorsal attention, cingulo-opercular, salience, frontoparietal A, frontoparietal B, default A, default B1, default B2, parietal memory, language). Supplemental analyses also assessed network density across all functional networks (not only association networks).

A geodesic distance matrix was computed using the Conte69 32k fs_LR group template surface and applied uniformly across individuals. Density was calculated at five radii (6 mm, 8 mm, 10 mm, 12 mm, 14 mm), and we reported the average value across these radii per vertex. Additionally, canonically low signal regions (the inferior temporal cortex and orbitofrontal cortex) were excluded from density calculations via a custom mask. This is visible in the whole brain view (see Supplemental Figure S8). The same mask was used for all subjects and was based on mode 1000 BOLD signal values from an independent set of 120 subjects from a previous study (Power et al., 2011).

### Density Rotation Analysis

In each individual, the left hemisphere rostral cingulo-opercular network region was defined as a region of interest (ROI) using Connectome Workbench’s cifti-cluster function, followed by manual selection of the rostral subregion from the resulting network clusters. Association network density was computed by averaging the density values of all vertices within this ROI. To assess whether the observed density was higher than expected by chance, a surface-based rotation analysis was performed. The ROI was randomly rotated across the individuals’ cortical surface 10000 times, preserving its shape and size. Rotated instances were retained only if all ROI vertices remained within a single association network in the individual-specific parcellation. For each valid rotation, the mean association network density was calculated. These values were aggregated across individuals to generate a null distribution, against which the true ROI value was compared (Figure 4C).

### Task Analysis

All tasks were analyzed using the general linear model (GLM) implemented via the 3dDeconvolve function in AFNI. All task conditions were included in the block design model. Surface-projected data were analyzed within individuals on a per-run basis. For each contrast, t-statistic maps were generated and converted to z-statistic maps in AFNI, then averaged across runs to produce a mean z-statistic map for each individual and contrast.

### Within Individual Task fMRI Reliability

While there is growing literature on the amount of resting-state fMRI data needed for reliable functional connectivity measures (Birn et al., 2013; Elliott et al., 2020; Gordon et al., 2017; Laumann et al., 2015; Noble et al., 2017), it is less clear how much data is required for reliable within-individual task fMRI. The answer likely depends on the task contrast and the design of the paradigm (e.g., block, event-related, or mixed). To ensure the robustness of our study, we first collected task data in two pilot individuals (PAN01 and PAN02), acquiring approximately twice as much data as in previous studies. This allowed us to assess test-retest reliability for each task. As shown in Supplemental Figure S5, we found that most tasks demonstrated strong test-retest reliability with 30 minutes of data per split half. We evaluated this based on the correlation of LPFC z-statistic maps. Based on these results, we proceeded with collecting at least 30 minutes of data per task per individual in the main study.

### Network Task Activity Comparison

To compare across cognitive domains, contrasts were grouped as described in the Task Paradigms section. For the language processing map, z-statistic maps from the [Sentence – Nonword] and [Intact Speech – Degraded Speech] contrasts were averaged. For the theory of mind map, [Emotional Pain – Physical Pain] and [False Belief – Photo] maps were averaged. For the episodic projection map, [Past Self – Present Self] and [Future Self – Present Self] maps were averaged. For each of 8 cognitive control tasks, a z-statistic map was created from the relevant high demand – low demand contrast (N-back: 2-back versus 0-back, Memory Span: 8 items versus 4 items, Multi-Source Interference: conflict versus no-conflict, Attention: sustained attention versus sensorimotor control).

To compare the activity of LPFC network regions, all individual network maps were masked using the LPFC mask described above. Mean z-values for each network were extracted from each of the contrast maps per individual, using their own LPFC individual-specific network maps, resulting in one z-value per network per individual per task.

For the specialized cognitive domains (Language Processing, Theory of Mind, Episodic Projection), one-tailed paired t-tests were used to compare the activation of the hypothesized target LPFC network—language, default B, and default A, respectively—against each of the other six LPFC association networks (language, default A, default B, frontoparietal, dorsal attention, cingulo-opercular). To correct for multiple comparisons, p-values were Bonferroni corrected for six comparisons (p × 6).

For the cognitive control tasks, one-tailed one-sample t-tests were used to identify LPFC association networks with significantly greater-than-zero activation. These p-values were Bonferroni corrected for seven comparisons (p × 7). All statistical tests were conducted using the *ttest* function in MATLAB.

For the domain-specific demands (language processing, theory of mind, episodic projection), we also used a whole-brain spatial permutation test to compare the network activations to a spatially aware chance-level baseline. Unthresholded task activation maps for each demand were randomly rotated around the surface 10,000 times and their network-level activations calculated to create a null distribution of network activity based on the shape and size of the networks. Significance was determined by a comparison of the real value to the null distribution.

### Task Visualization

The activation maps for all tasks were visualized relative to the boundaries of the relevant LPFC networks. In all figures, activity was thresholded to display the top 25% of LPFC vertices per hemisphere, based on z-statistic values, excluding any vertices with z < 0.

### Task/Network Overlap

The spatial overlap between thresholded task activation maps and different network parcellations—individual-specific, group-average, and other-individuals’ maps—was quantified using the Dice similarity coefficient (Dice, 1945). Individual-specific task activation maps were thresholded at three levels (top 15%, 20%, and 25% of LPFC vertices) and compared to each LPFC network parcellation. For each individual and task, the average Dice overlap values (across thresholds) were compared between individual networks and (1) group-averaged networks or (2) other-individual networks using one-sided paired t-tests. P-values were corrected for 2 comparisons (p × 2). To test for regional specificity, the difference in Dice overlap between individual-specific and other-individual networks was compared for LPFC versus non-LPFC regions. The same procedure described above was repeated for non-LPFC task activations and networks. LPFC and non-LPFC values were compared using one-sided paired t-tests for each task and a composite score averaged across the three tasks.

### Border Activation Analysis

It was observed that cognitive control activations appeared near the borders of multiple control networks (frontoparietal, dorsal attention, cingulo-opercular). This was quantified by comparing the mean distance of activated versus non-activated FP, DAN, and CO network vertices (at z > 1, z > 2, z > 3) to the nearest vertex of either of the other two control networks (for FP, the nearest DAN or CO vertex, for example). Minimum distances were measured from each vertex to each network on the Conte32k_fsLR template surface which was used across all individuals. A similar procedure was followed for language, theory of mind, and episodic projection activations except that we compared the mean distance of activated versus non-activated vertices for each target network (LANG, DN-B, DN-A, respectively) to any other network.

### Task Activation Similarity Analysis

To compare the similarity of vertex-wise activation patterns across different demands within and across individuals (Figure 8B), unthresholded task activation maps masked to the LPFC were correlated within and across individuals. For network-level comparisons, mean LPFC network activation z-values were calculated for each network, and these values were correlated within and across individuals.

### Seed Connectivity Analyses

Seed-based connectivity was used to validate network parcellations throughout the manuscript. In each case, the seed was manually selected from within network boundaries to best represent the network’s characteristic connectivity pattern.

### Variant ROI Task Analyses

Language and default B variant regions described in the manuscript were defined as ROIs by clustering the relevant network using Connectome Workbench’s cifti-cluster function and manually selecting the cluster corresponding to the variant. In cases where the variant partially overlapped with the canonical group-averaged network, the overlapping portion of the group-averaged network cluster was subtracted from the manually defined ROI. Task activation z-statistics were averaged across all vertices within each ROI for both the variant individual and all individuals without a variant in that location. A variant task activation metric was computed as the difference between the relevant specialized demand (language processing or theory of mind) and the expected cognitive control demand (spatial memory). This metric was then compared between variant individuals and non-variant individuals.

### Task Co-activation and Functional Connectivity

The relationship between resting-state functional connectivity, network membership, and task co-activity during cognitive control demands was examined by first classifying vertices within control networks (FP, CO, DAN) as either active or inactive for each cognitive control task based on a z-statistic threshold (z > 2). Using resting-state data, we then computed the average functional connectivity between vertices belonging to each activity-defined group.

## Data and Software Availability

Preprocessed CIFTI resting-state and task timecourses have been deposited in the OpenNeuro data repository (https://openneuro.org/datasets/ds006598/versions/1.0.0). Code to perform analysis is available at https://github.com/GrattonLab/Ladwig-LPFC.

## Declaration of generative AI and AI-assisted technologies in the writing process

During the preparation of this work the author(s) used chatGPT in order to improve the readability of the text. After using this tool/service, the author(s) reviewed and edited the content as needed and take(s) full responsibility for the content of the publication.

## Declaration of competing interests

The authors declare no competing interests.

## CONTRIBUTIONS

Z.L., J.H., M.D., and C.G. designed the study. Z.L., J.H., M.D., and N.L. collected the data. Z.L., K.Z.K., Y.P., E.H., A.D., D.M.S., D.E.N., S.E.P., R.M.B., and C.G. analyzed and interpreted the data. Z.L. conducted formal analyses and visualized results. Z.L. and C.G. wrote the manuscript.

## ACKNOWLEDGEMENTS

This work was supported with funds from NIH grant R01MH118370 (CG), NIH grant R01NS124738 (CG), NIH grant R01MH121509 (DEN), NSF grant NSFCAREER 2305698 (CG), NIH training grant T32 AG020506 (ZL), NIH grant P30AG013854 (RMB), and the Therapeutic Cognitive Neuroscience Fund (DMS). This research was supported in part through the computational resources and staff contributions provided by the Center for Translational Imaging and Quest high performance computing facility at Northwestern University, which is jointly supported by the Office of the Provost, the Office for Research, and Northwestern University Information Technology.

## Notes

### Competing Interest Statement

The authors have declared no competing interest.

### Summary of Updates

The manuscript has been updated to include new supplementary analyses as well as a revised version of Figure 8.

