## Supplemental Text and Figures for "Precision fMRI reveals densely interdigitated network patches with conserved motifs in the lateral prefrontal cortex"

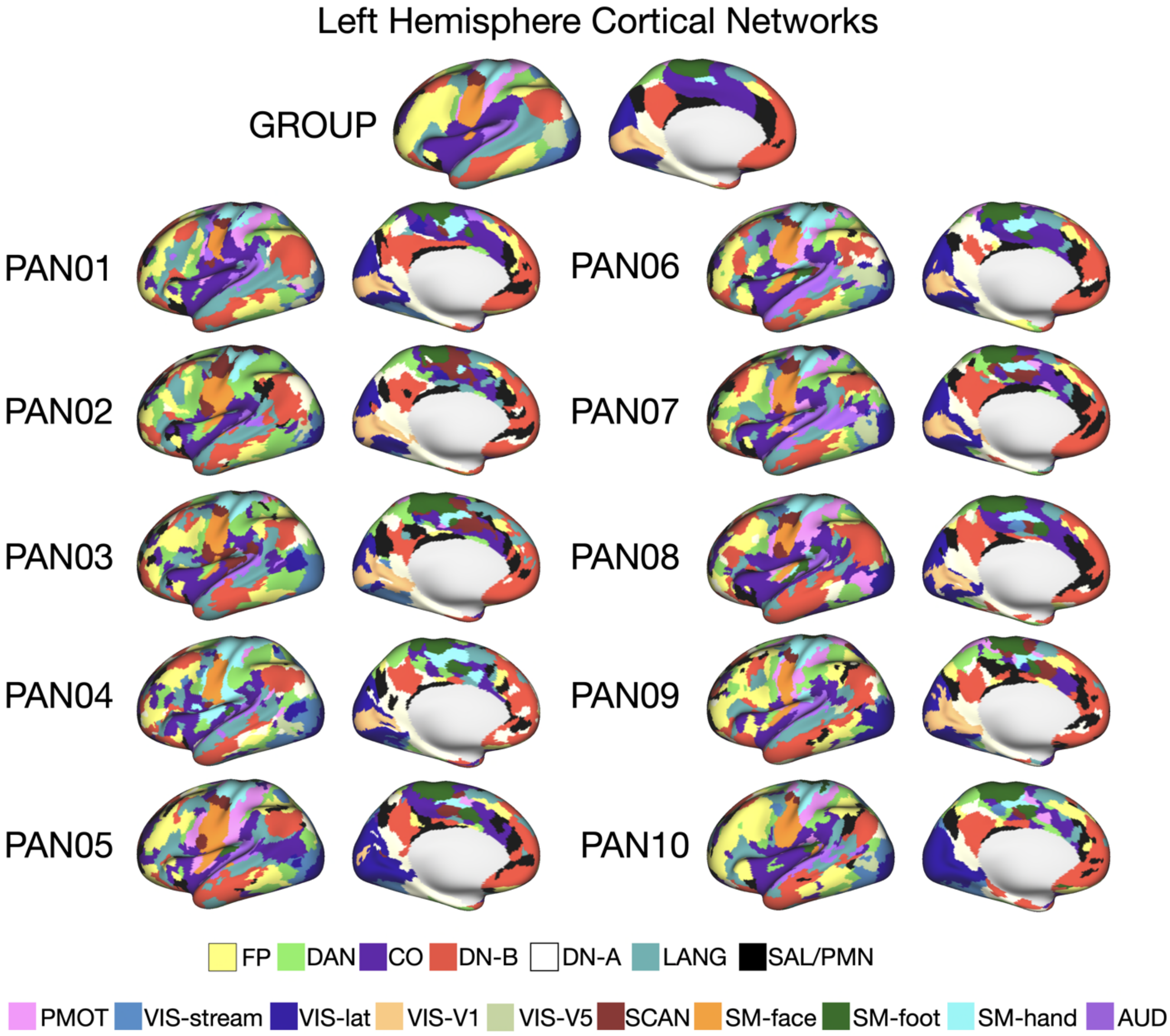


**Supplemental Figure S1. Individual-specific cortex-wide network parcellations (left hemisphere).** Network parcellations were generated for all ten individuals using a cortex-wide network identification protocol adapted from Lynch et al. (2024). While the main text focuses on the lateral prefrontal cortex (LPFC), parcellations were defined across the entire cortical surface to ensure comprehensive coverage. Substantial individual variability in network topography is evident across the cortex.

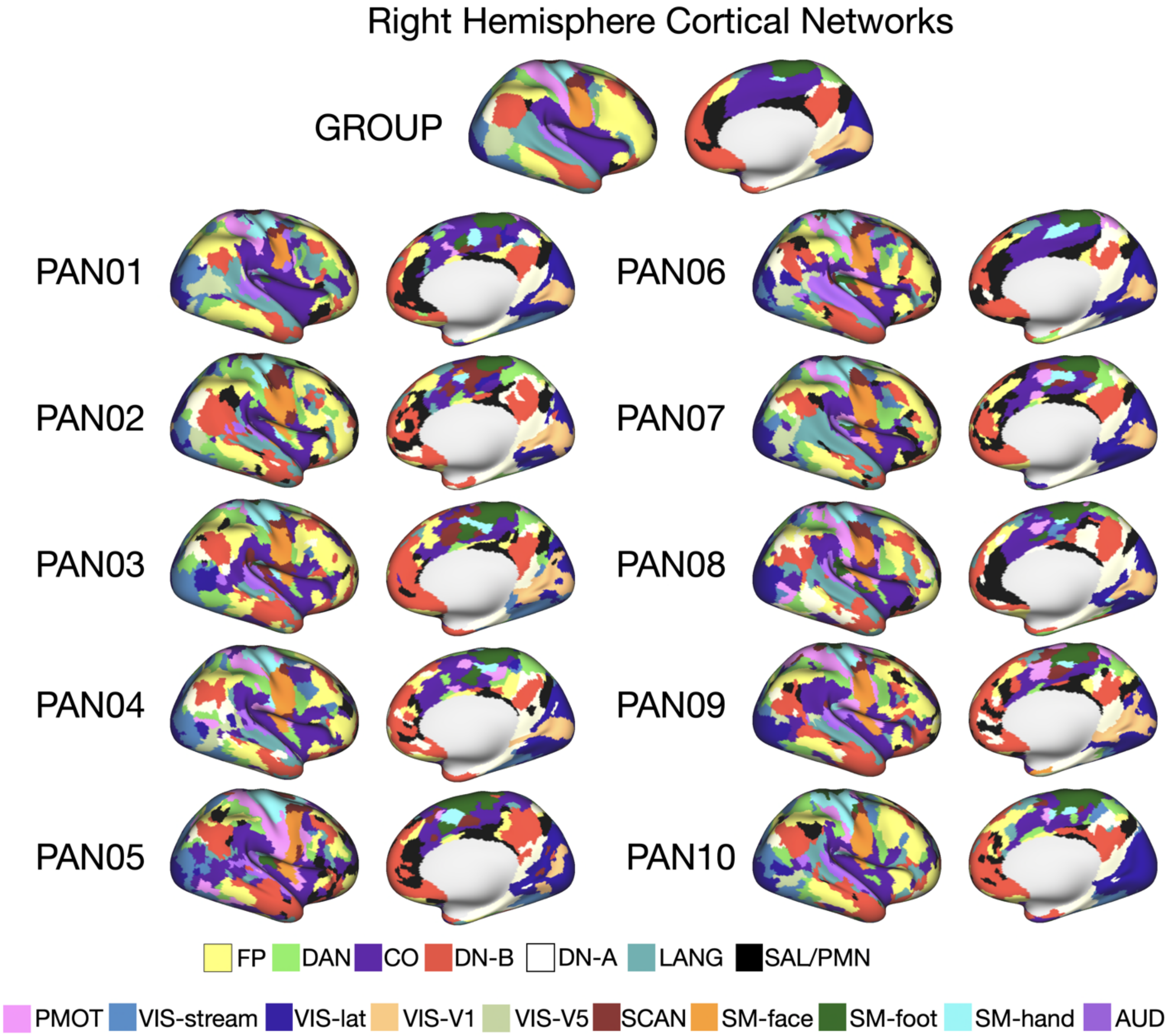


**Supplemental Figure S2. Individual-specific cortex-wide network parcellations (right hemisphere).** Network parcellations were generated for all ten individuals using a cortex-wide network identification protocol adapted from Lynch et al. (2024). While the main text focuses on the lateral prefrontal cortex (LPFC), parcellations were defined across the entire cortical surface to ensure comprehensive coverage. Substantial individual variability in network topography is evident across the cortex.

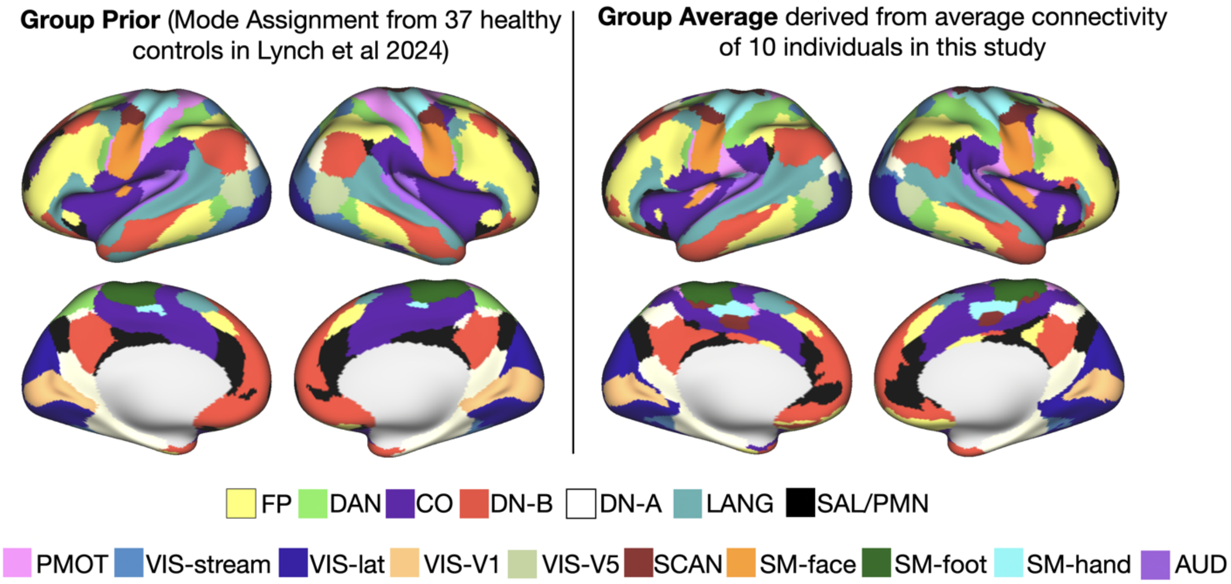


**Supplemental Figure S3. Group-average network priors are consistent across samples***.* To assess the robustness of group-average network structure, an alternative parcellation was generated using the same network identification as for individuals but using the average functional connectivity matrix from the ten individuals in the present study. This was visually compared to the mode parcellation derived from 37 healthy adults reported in Lynch et al. (2024). The resulting network maps were similar, indicating that the group-average is stable across independent samples and derivation methods.

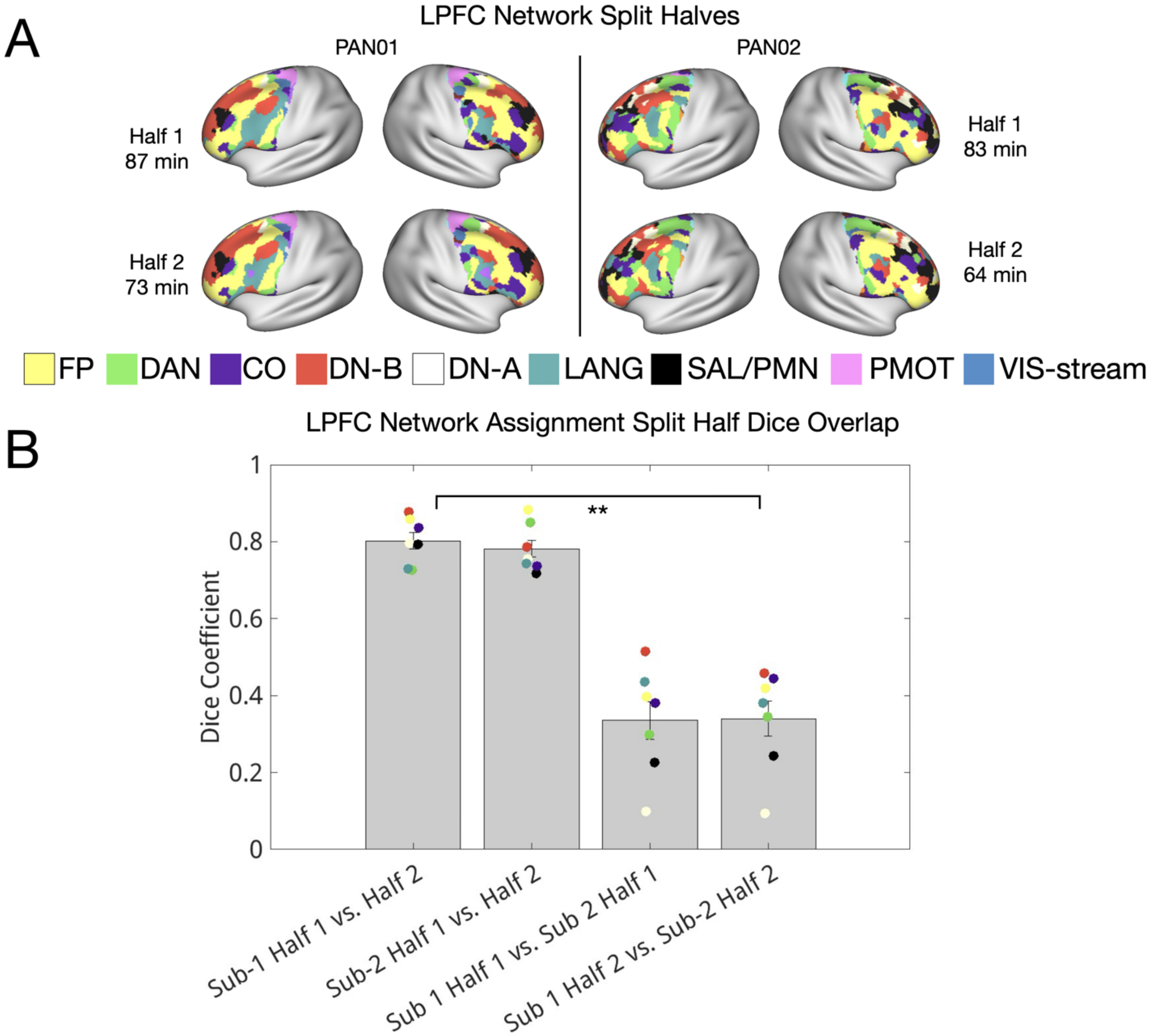


**Supplemental Figure S4. LPFC network assignments are reliable within individuals across sessions and distinct across individuals.** (A) For two high-data individuals (PAN01 and PAN02), resting-state fMRI data were divided into two independent split-halves based on scanning day (odd vs. even sessions). Network parcellations were generated separately for each half using the same identification protocol. We visually observed LPFC network topographies were highly similar across halves within individuals but different across individuals. (B) This was quantified with the dice coefficient per network (mean within-individual Dice coefficient = 0.79 ± 0.05, mean between-individual Dice coefficient = 0.34 ± 0.13) and confirmed with a paired-samples t-test ($t(6) = 9.8$, $p < 0.0001$), indicating robust within-subject reliability and pronounced individual specificity.

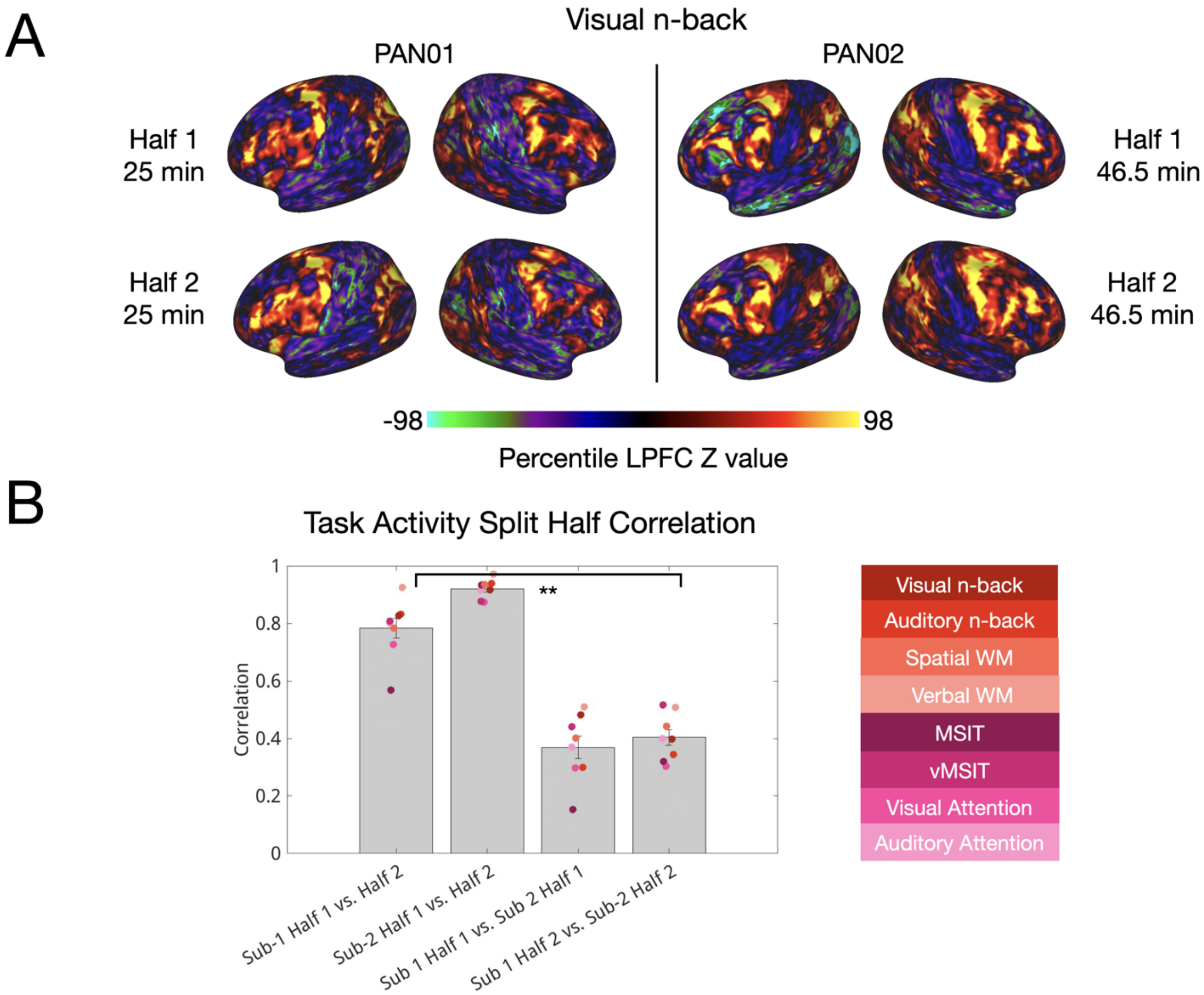


**Supplemental Figure S5. LPFC task activation patterns are reliable within individuals and distinct across individuals.** (A) For two high-data individuals (PAN01 and PAN02), task fMRI data were divided into two independent split-halves based on scanning day (odd vs. even sessions). Z-statistic maps were computed separately for each split-half. We found that these maps were visually similar across split halves within an individual and different between individuals. Visual n-back maps are shown as an example. (B) Overall, these task-evoked activation patterns in the lateral prefrontal cortex (LPFC) were more similar within individuals than across individuals (mean within-individual correlation = 0.85 ± 0.06, mean between-individual correlation = 0.39 ± 0.09, confirmed with a paired-samples t-test $t(7) = 21.6$, $p < 0.000001$). This highlighted the reliability and idiosyncrasy of individual activation profiles.

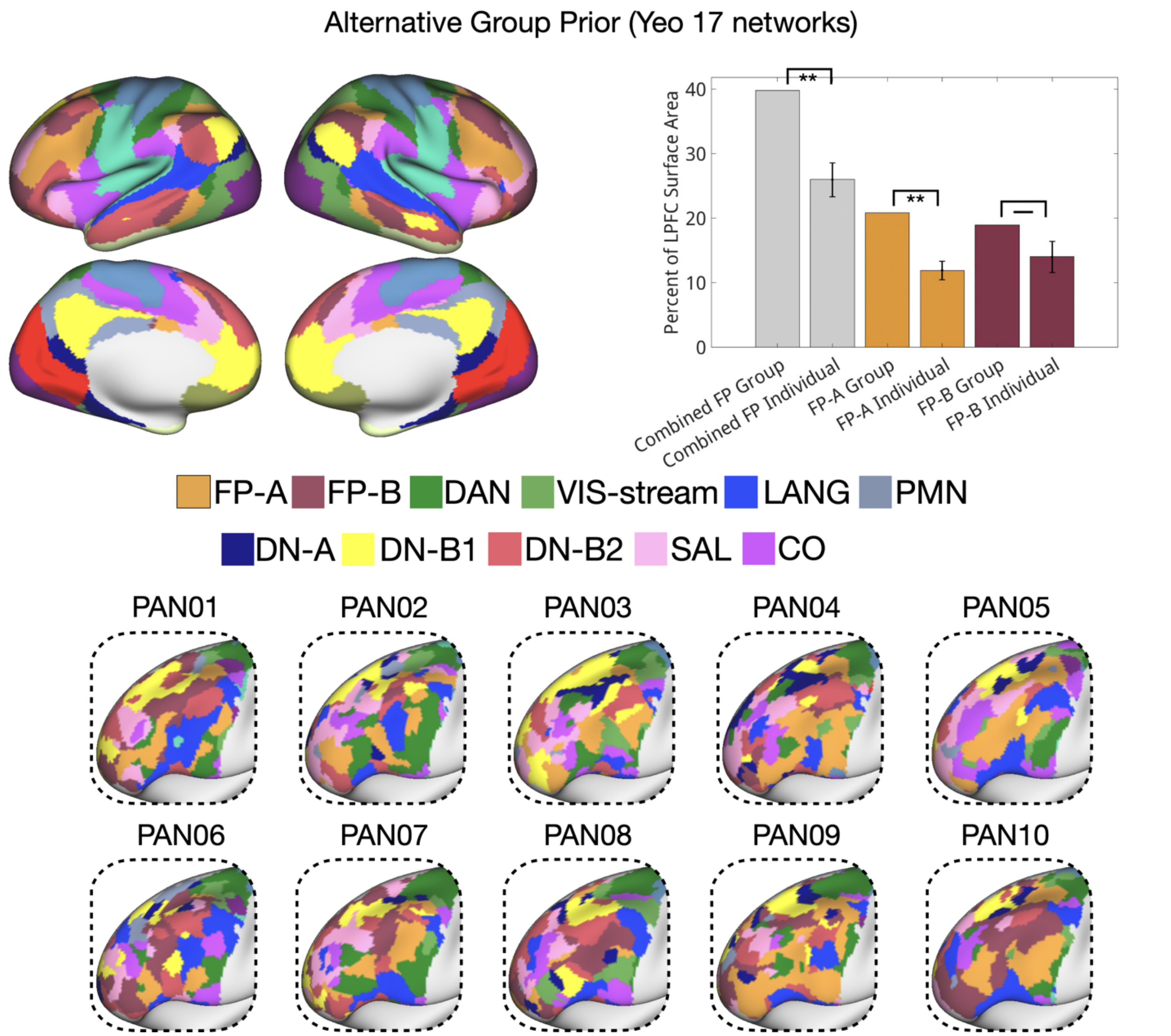


**Supplemental Figure S6. Replication of network composition results using an alternative group prior with two frontoparietal networks.** To assess the robustness of our findings, individual-specific LPFC parcellations were re-derived using an alternative group prior that included two distinct frontoparietal networks (Yeo 17 networks, FP-A and FP-B; Yeo et al., 2011). The combined frontoparietal territory (FP-A + FP-B) occupied significantly more LPFC surface area in the group prior than in individual-specific parcellations (one-sample t-test: $t(9) = 5.3$, $p = 0.0005$), replicating the primary finding of inflated frontoparietal representation in group-level maps. When examined separately, FP-A was significantly larger in the group prior (one-sample t-test: $t(9) = 6.20$, $p = 0.0001$), while FP-B showed a non-significant trend in the same direction (one-sample t-test: $t(9) = 2.04$, $p = 0.07$). Significant ($p < 0.05$) differences are indicated with asterisks.

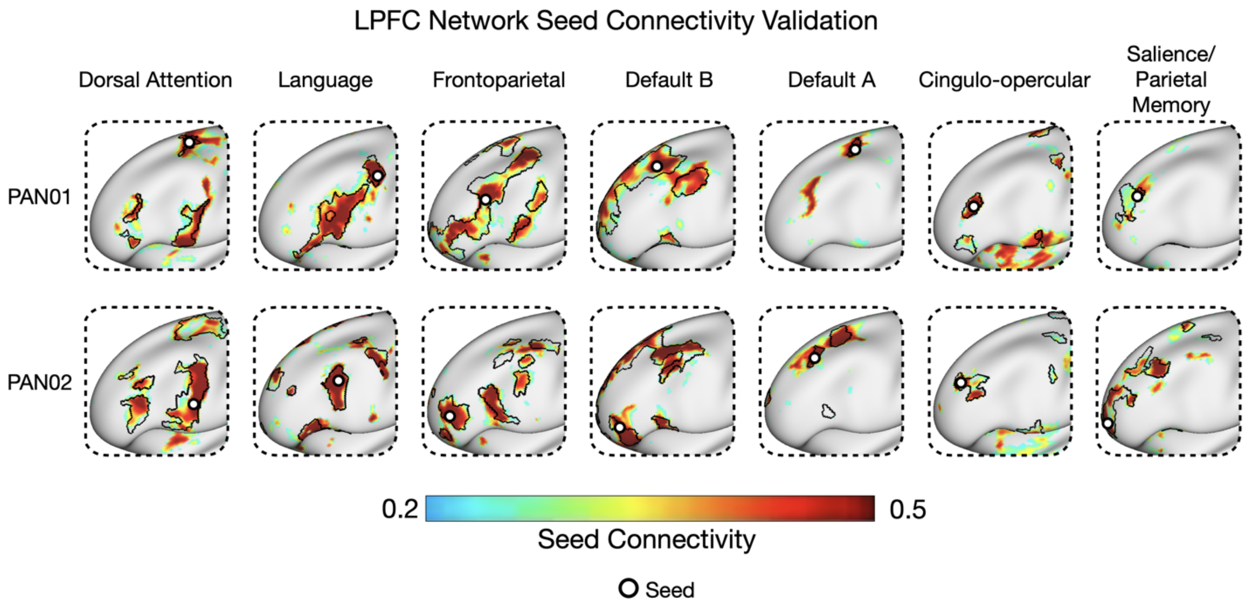


**Supplemental Figure S7. LPFC network parcellations are validated by seed-based functional connectivity.** To validate the fine-grained organization observed in LPFC network parcellations, seed-based connectivity maps were generated for two exemplar individuals (PAN01 and PAN02). Seeds were manually placed within the boundaries of each association network in the LPFC. The resulting connectivity maps closely matched the parcellated network territories, including the spatial patchiness observed relative to group-average maps, providing converging evidence for the reliability of individual-specific parcellations.

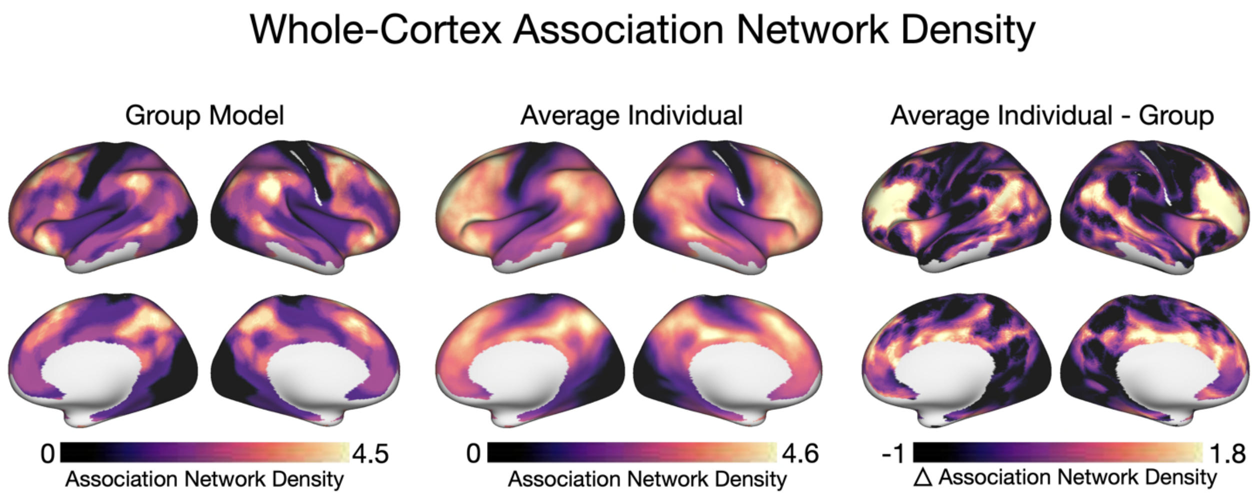


**Supplemental Figure S8. Association network density is underestimated by group averaging.** Association network density is shown across the cortical surface for the group-average map (left), the across-individual average (middle), and the difference between the two (right). While broad patterns are generally preserved, group averaging underestimates association network density in the rostral LPFC.

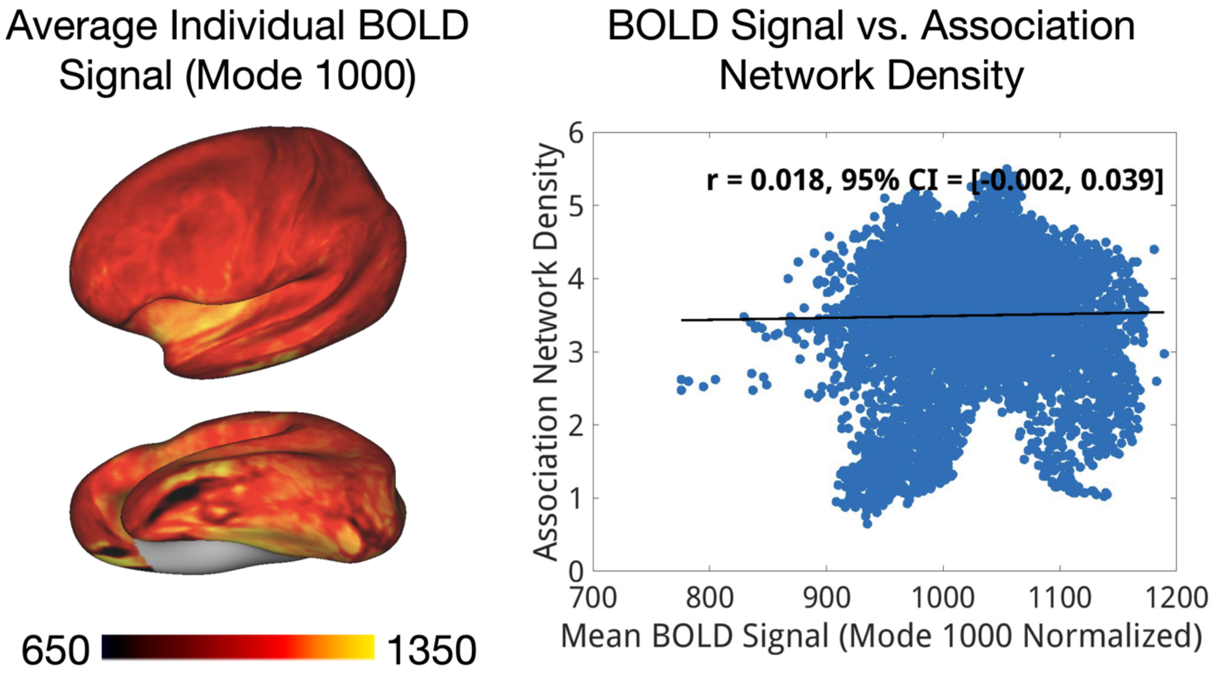


**Supplemental Figure S9. There is no meaningful relationship between association network density and BOLD signal in the LPFC.** A measure of BOLD signal (after mode 1000 normalization) across individuals is shown across the left hemisphere cortical surface. Visually, there is no obvious pattern that corresponds with the high rostral association network density shown in Figure 2D and 2E. This is quantified (right) – the correlation between BOLD signal and association network density in the LPFC was near zero (r = 0.018, 95% CI: [–0.002, 0.039]), suggesting no meaningful relationship.

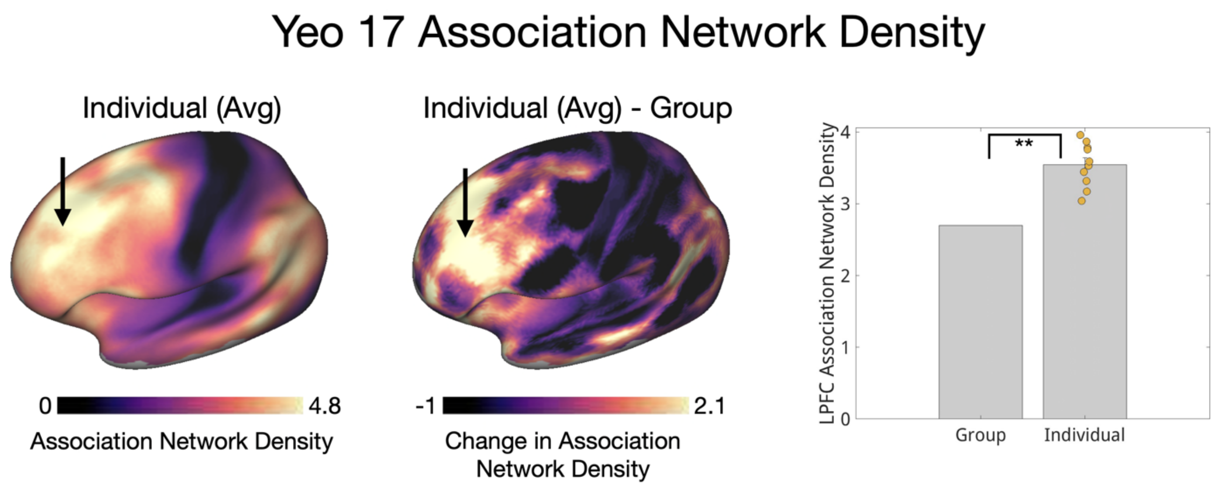


**Supplemental Figure S10. Replication of association network density using Yeo 17 group average atlas.** Association network density is shown across the cortical surface for the across-individual average (left) and the difference between group-average and individual average (middle) using the Yeo 17 group average. Marked by an asterisk, individuals showed significantly higher association network density compared to the group average (one-sample t-test: $t(9) = 8.82$, $p = 0.00001$)

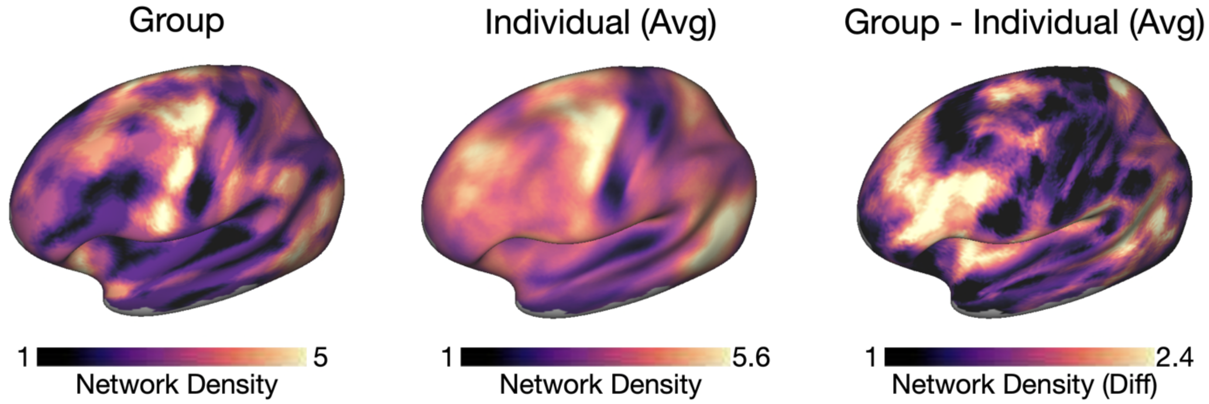


**Supplemental Figure S11. Cortical maps of network density including all functional networks.** Network density maps were created for each individual using the same method as for association network density but including all functional networks rather than only association networks. These maps resemble association network density maps but also highlight regions where association networks sit close to local networks (e.g. caudal LPFC). However, these regions tended to show high density in both the group and individual maps, so the difference map closely resembles the association network density map.

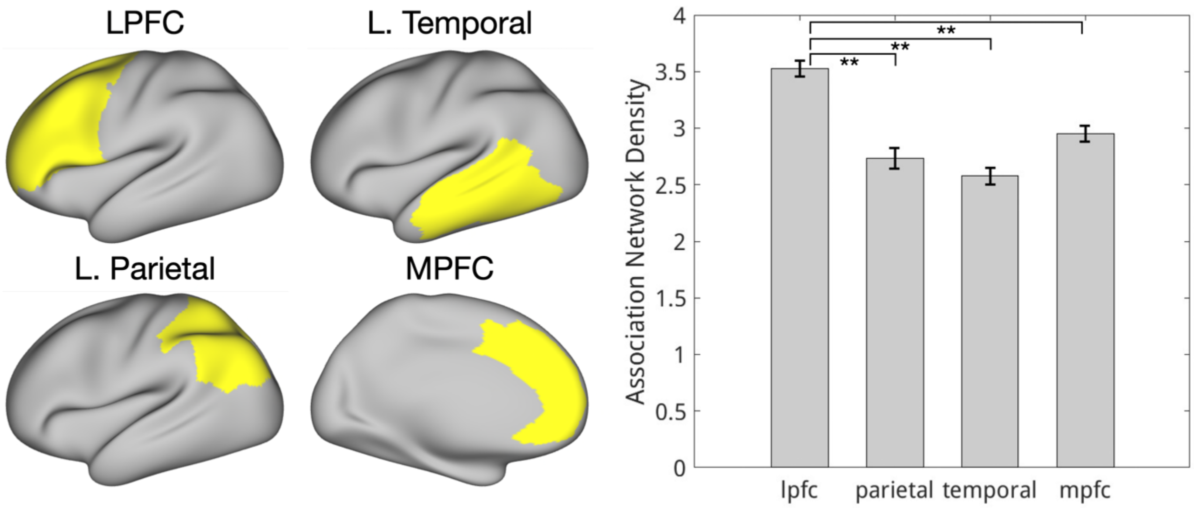


**Supplemental Figure S12. The lateral prefrontal cortex exhibits higher association network density than other association regions.** ROIs were defined for three other association regions of the brain (lateral temporal, lateral parietal, and medial prefrontal cortex). The average association network density within those regions was calculated for each individual. The lateral prefrontal cortex exhibited higher association network density than the lateral parietal cortex (paired t-test: $t(9) = 6.0$, $p < 0.001$), the lateral temporal cortex (paired t-test: $t(9) = 10.0$, $p < 0.001$), and the medial prefrontal cortex (paired t-test: $t(9) = 8.1$, $p < 0.001$).

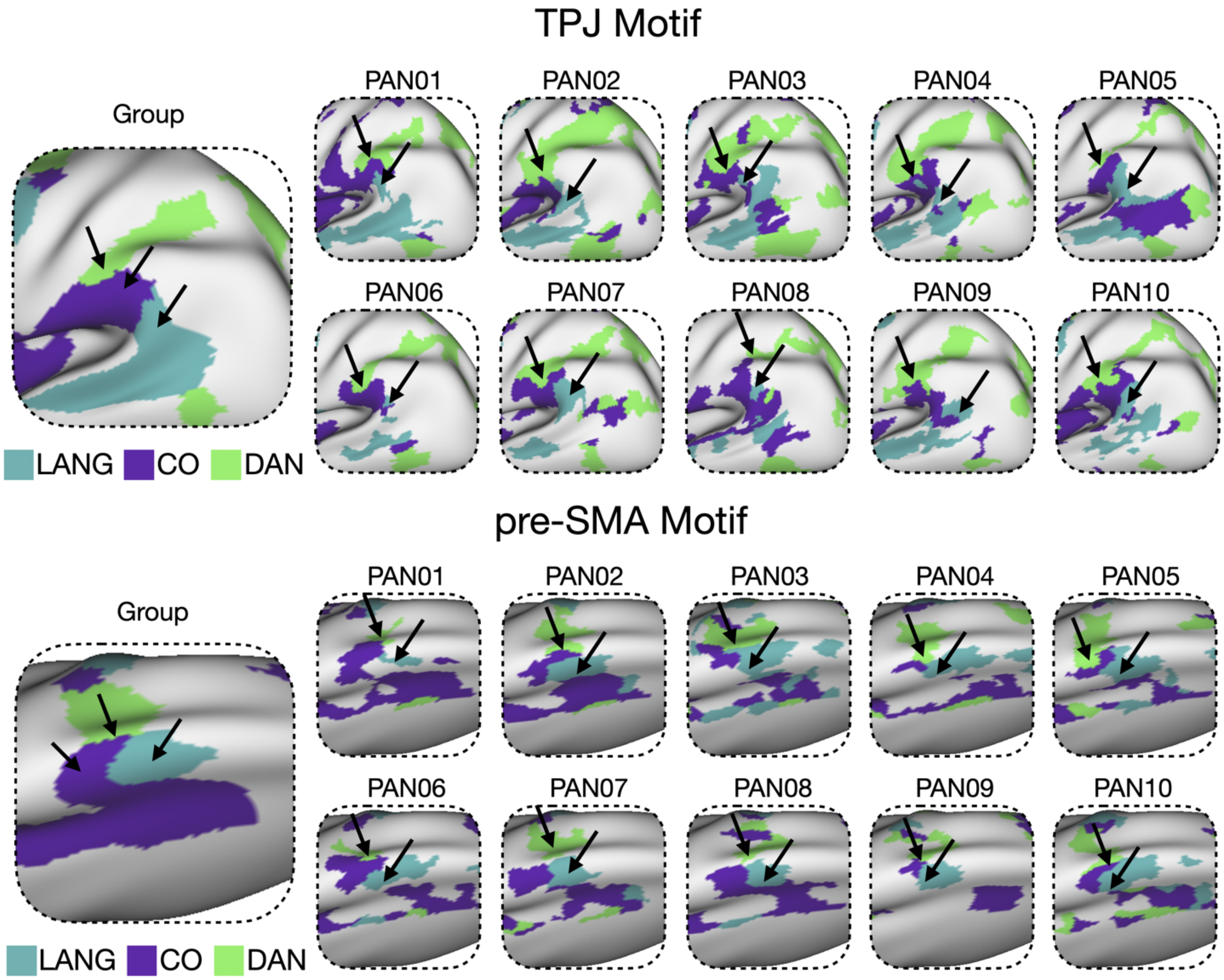


**Supplemental Figure S13. The DAN–CO–LANG motif recurs in both TPJ and pre-SMA across individuals.** Network parcellations for each individual were manually examined for evidence of a repeating three-network motif involving the dorsal attention network (DAN), cingulo-opercular network (CO), and language network (LANG). This motif was observed in all ten individuals in both the temporoparietal junction (TPJ) and the pre-supplementary motor area (pre-SMA). In both regions, the spatial arrangement of the motif was sufficiently consistent across individuals that it is also evident in the group-average map.

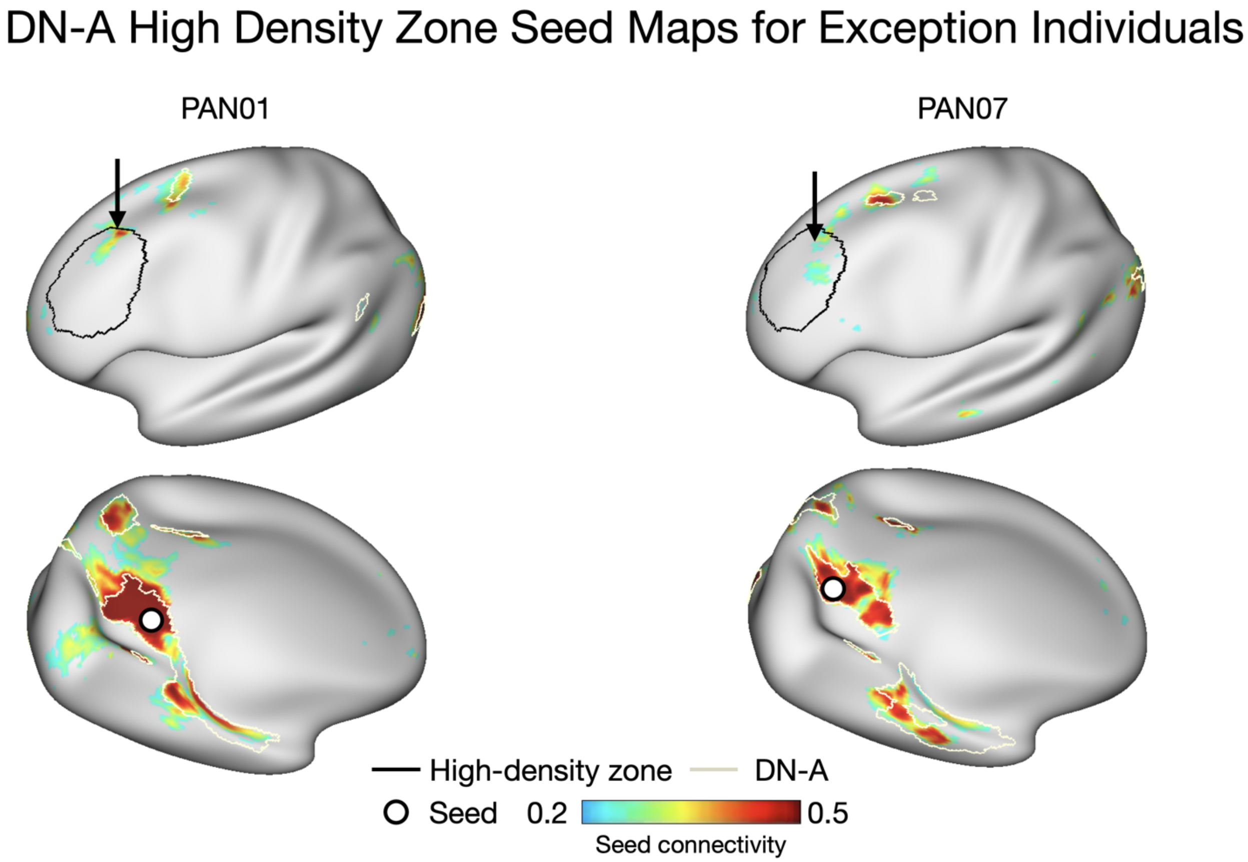


**Supplemental Figure S14. In exception cases, DN-A shows seed-based connectivity to the rostral high-density zone.** PAN01 and PAN07 did not exhibit DN-A network territory within the rostral LPFC high-density zone based on their individual parcellations. However, manually selected seeds within DN-A in these individuals showed functional connectivity to this location. This suggests the possible presence of DN-A–related signal in this region that was not captured by the discrete parcellation boundaries. The high-density zone shown is the same as in Figure 4 – a 15mm radius around the rostral CO region.

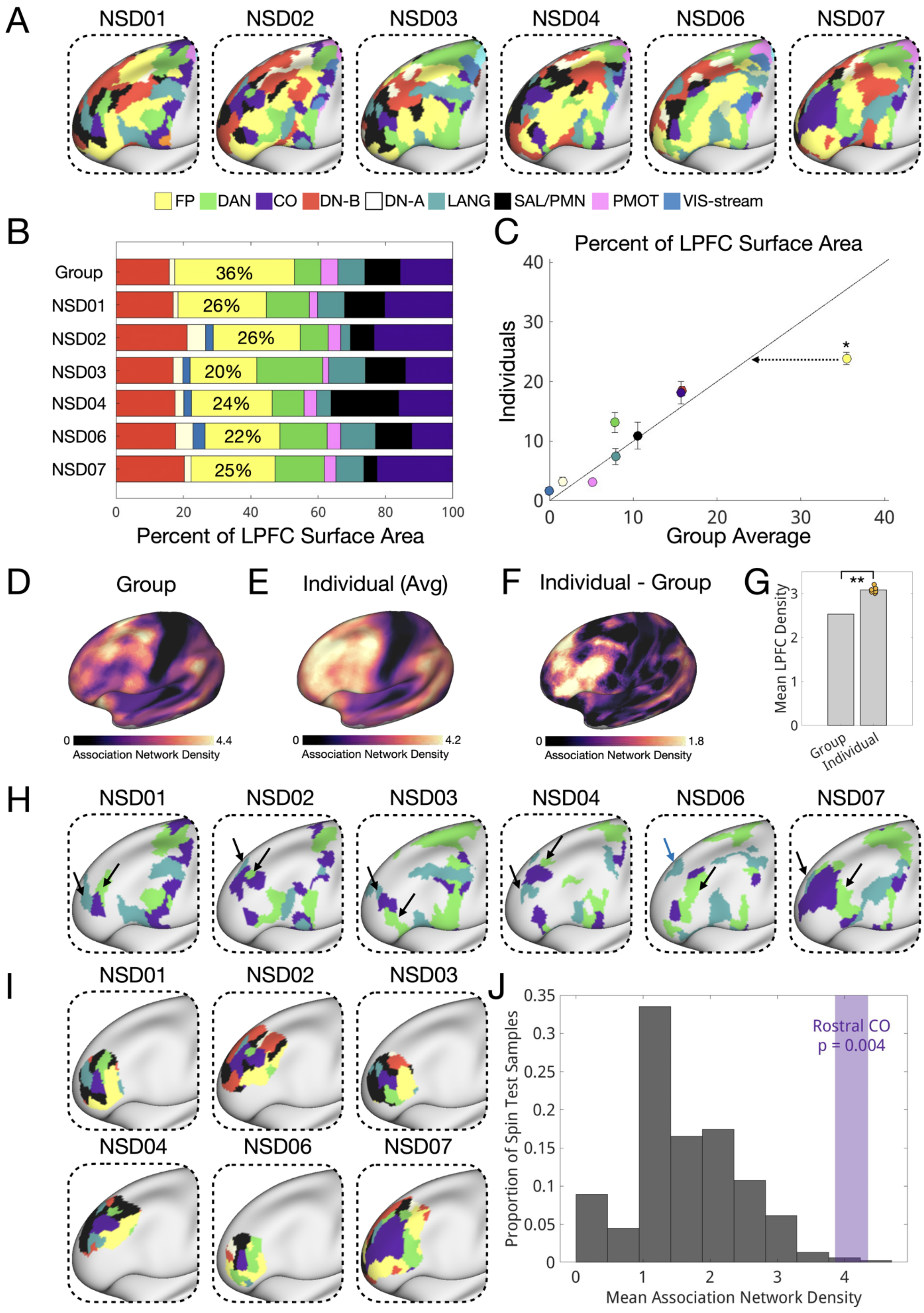

A
NSD01
NSD02
NSD03
NSD04
NSD06
NSD07
FP
DAN
CO
DN-B
DN-A
LANG
SAL/PMN
PMOT
VIS-stream
B
Group
NSD01
NSD02
NSD03
NSD04
NSD06
NSD07
36%
26%
26%
20%
24%
22%
25%
0
20
40
60
80
100
Percent of LPFC Surface Area
C
Percent of LPFC Surface Area
Individuals
Group Average
0
10
20
30
40
*
D
Group
E
Individual (Avg)
F
Individual - Group
0
4.4
4.2
1.8
Association Network Density
G
Mean LPFC Density
**
Group
Individual
H
NSD01
NSD02
NSD03
NSD04
NSD06
NSD07
I
NSD01
NSD02
NSD03
NSD04
NSD06
NSD07
J
Proportion of Spin Test Samples
0
0.05
0.1
0.15
0.2
0.25
0.3
0.35
Rostral CO
p = 0.004
Mean Association Network Density

**Supplemental Figure S15. Resting-state results replicate in an independent publicly available precision fMRI dataset.** (A) LPFC networks were generated for 6 individuals from the Natural Scenes Dataset (NSD). (B,C) They exhibited smaller frontoparietal networks versus the group average (Group Average: 35.5%, Individual Average = 23.9% ± 2.5%, one-sample t-test: $t(5) = 11.6$, $p < 0.0001$). (D-G) They exhibited higher LPFC association network density versus the group average (Group Average: 2.53, Individual Average=3.08 ± 0.02, one-sample t-test: $t(5) = 19.1$, $p < 0.00001$). (H) 5 of 6 subjects exhibited the anterior LANG-CO-DAN motif. (I,J) They exhibited higher than expected association network density at rostral CO (spin test $p = 0.004$).

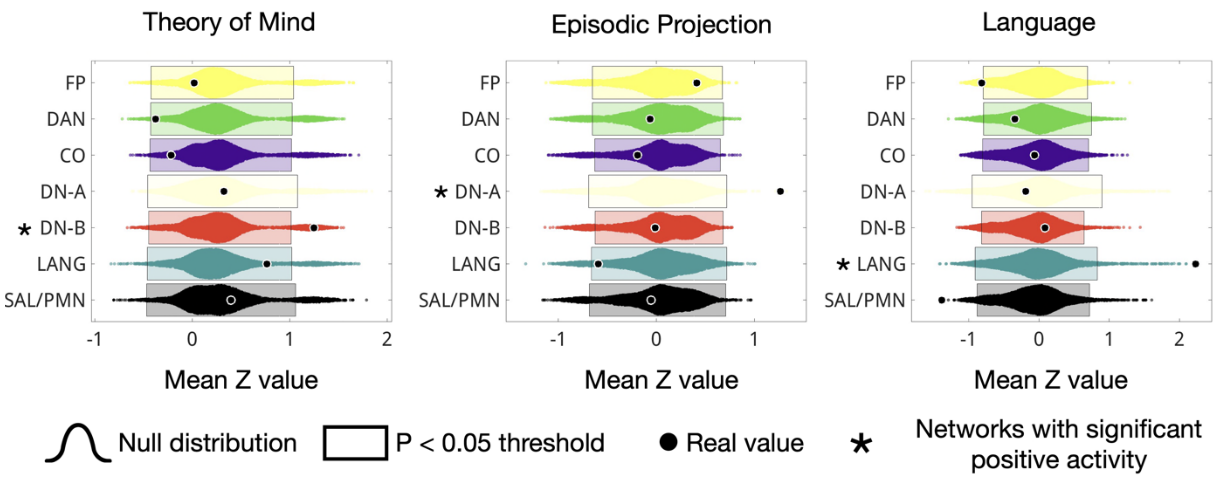


**Supplemental Figure S16. Theory of mind, episodic projection, and language demands preferentially recruit distinct individual-specific networks versus a spatially permutation null distribution.** Unthresholded task activation maps for theory of mind, episodic projection, and language demands were randomly rotated around the surface 10,000 times and their network-level activations calculated to create a null model of network activity based on the shape and size of the networks. This activity distribution is shown for each network for each task, along with the thresholds for significant activation ($p < 0.05$) and the real value. For theory of mind, only DN-B showed significant activity, for episodic projection, only DN-A showed significant activity, and for language, only LANG showed significant activity.

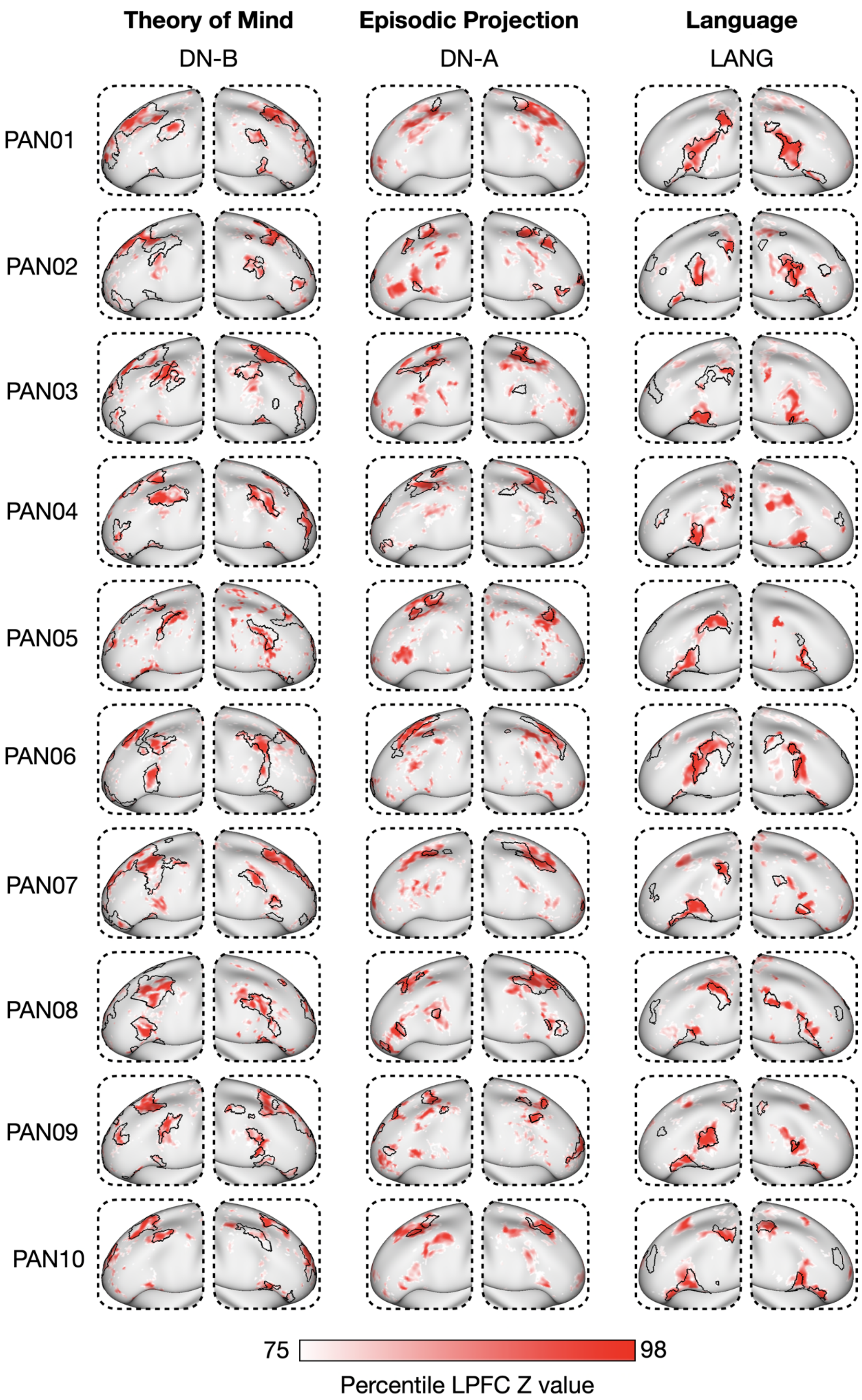
Theory of Mind
DN-B
Episodic Projection
DN-A
Language
LANG
PAN01
PAN02
PAN03
PAN04
PAN05
PAN06
PAN07
PAN08
PAN09
PAN10
75
98
Percentile LPFC Z value

**Supplemental Figure S17. Theory of mind, episodic projection, and language task activation maps for all individuals.**

The top 25% task-active LPFC vertices (ranked by z-statistic) are shown for all individuals for theory of mind, episodic projection, and language tasks, overlaid with individual-specific borders for the relevant network (DN-B, DN-A, LANG).

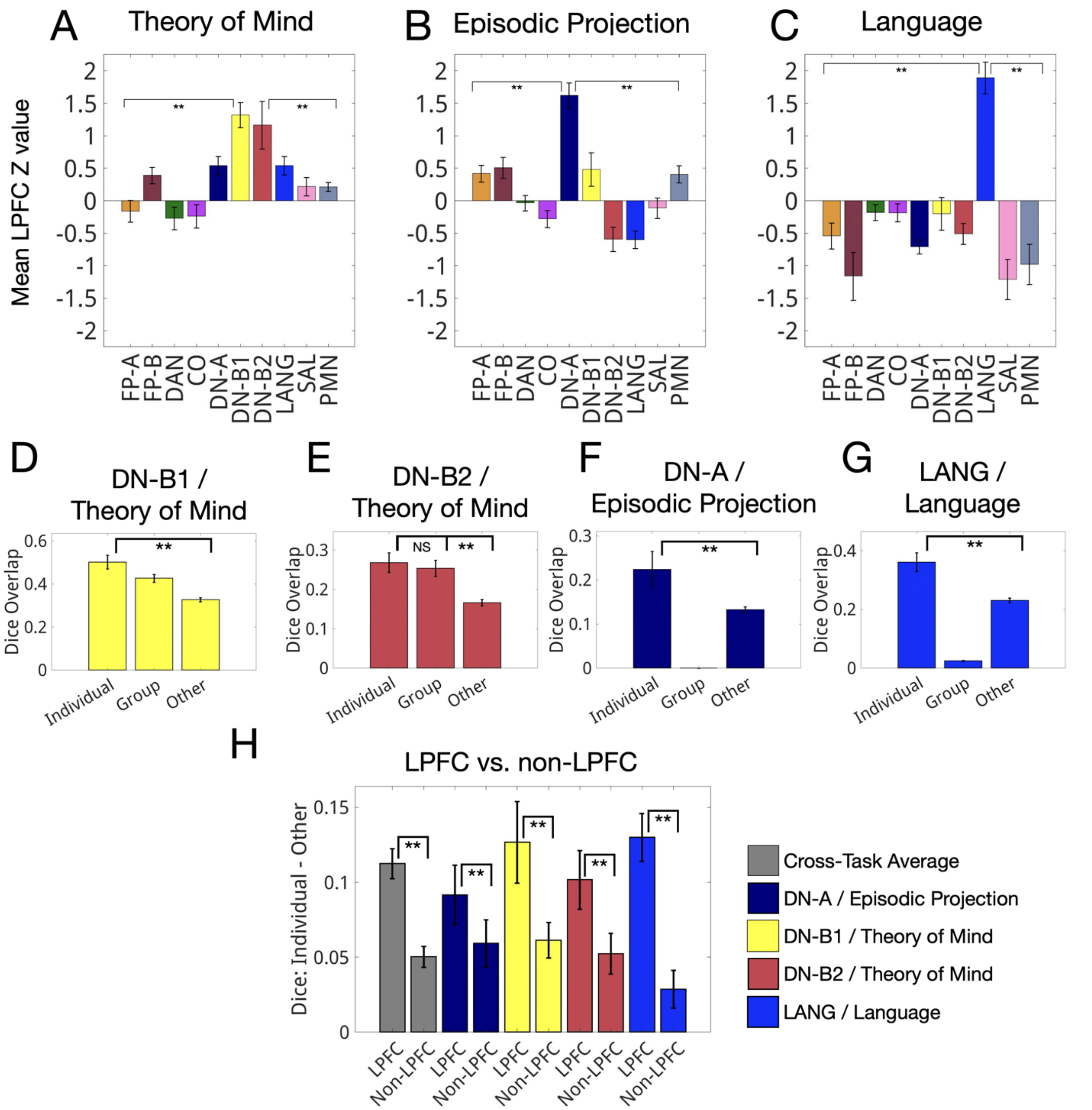


**Supplemental Figure S18. Domain-specific network-level task preferences replicate with an alternative parcellation.** An alternative network parcellation prior (Yeo 17) was used to identify LPFC networks in individuals and their network-level task preferences were evaluated. (A) Theory of mind preferentially activated default B1 and B2 (all comparisons, corrected $p < 0.01$). (B) Episodic projection preferentially activated default A (all comparisons, corrected $p < 0.01$). (C) Language processing preferentially activated the language network (all comparisons, corrected $p < 0.01$). (D-G) Task activations showed significantly greater overlap with individual-specific LPFC networks than with either group-average networks or non-specific individual networks (corrected $p < 0.05$ for all comparisons). (H) This effect was larger in the LPFC than in non-LPFC cortical regions ($p < 0.01$).

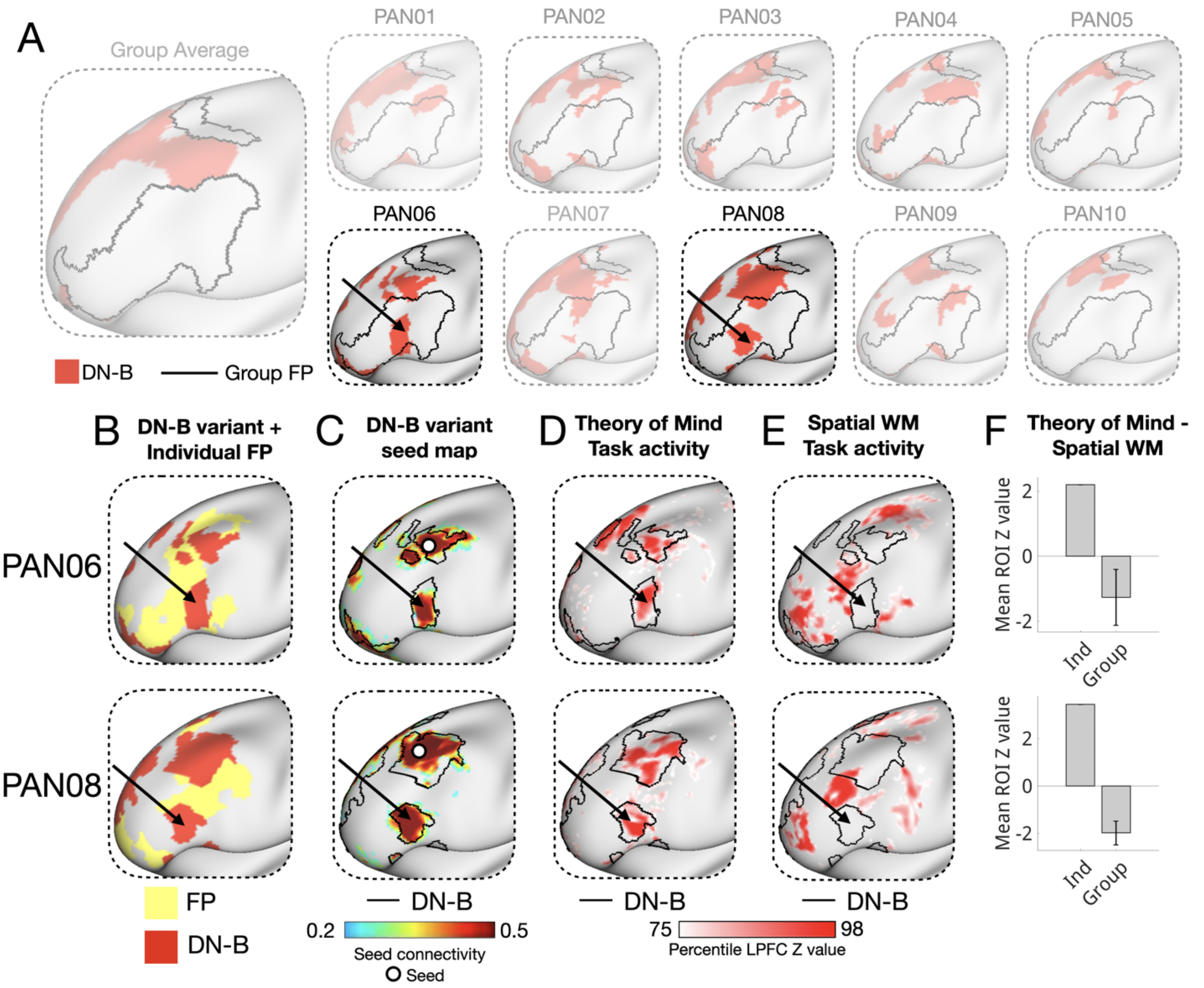


**Supplemental Figure S19. Idiosyncratic default B regions are embedded within canonical frontoparietal territory in a subset of individuals and are validated through multiple measures.** (A) Default B network parcellations for 10 individuals and the group average, overlaid with the group-average frontoparietal (FP) network. Two individuals (PAN06, PAN08) showed large default B regions in mid-LPFC, well outside typical default B territory. (B) These variant default B regions were interdigitated with individual-specific FP regions. (C) Seed-based connectivity from the variant regions showed coupling with canonical LPFC default B network regions, supporting their network identity. (D) Theory of mind task activations showed positive responses in the variant default B regions but not in adjacent FP regions. (E) Spatial working memory activations showed the opposite pattern: activation in adjacent FP regions but not in the variant default B regions. (F) Theory of mind > spatial working memory *z*-values are shown for the variant individuals and for the same ROI locations averaged across the eight non-variant individuals. Variant-region ROIs in non-variant individuals did not show theory of mind > spatial WM responses, confirming their individual-specific nature.

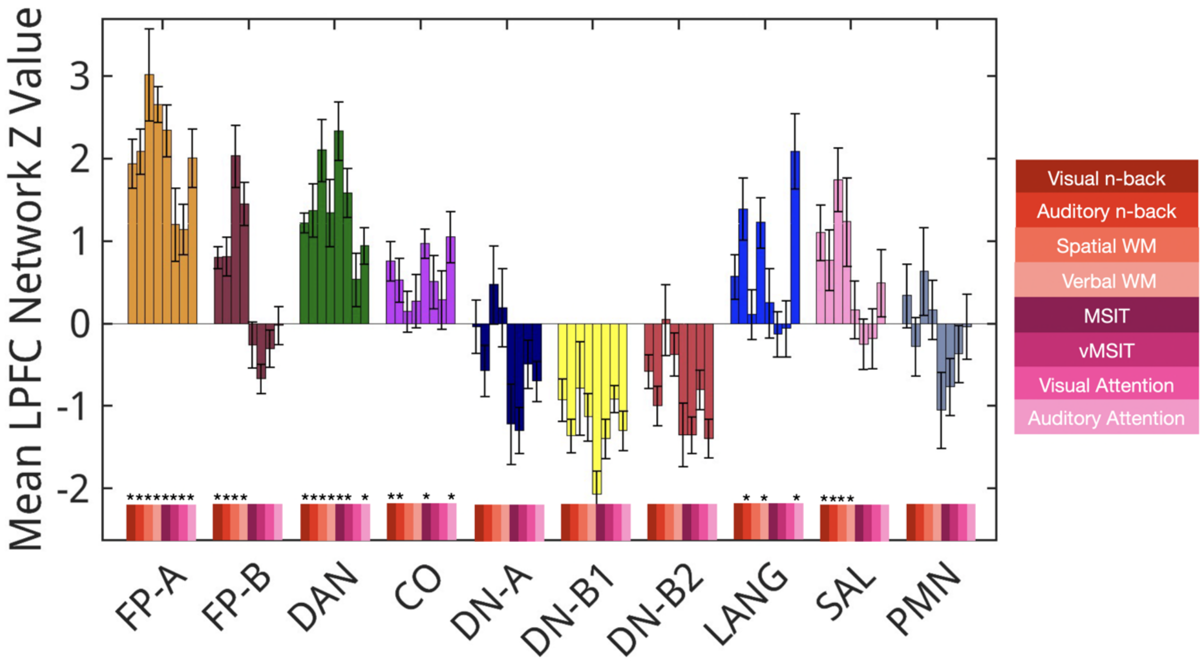


**Supplemental Figure S20. Cognitive control network-level task preferences are similar with an alternative parcellation**. Using the Yeo 17 network prior, mean network-level LPFC activations (z value) were computed for all eight cognitive control tasks. The dorsal attention (DAN), frontoparietal-A (FP-A), frontoparietal-B (FP-B) and cingulo-opercular (CO) networks were the most consistently engaged across tasks (8/8, 8/8, 4/8 and 4/8 tasks, respectively). Language and salience networks were also selectively engaged (3/8 and 4/8 tasks). Significant activations (corrected $p < 0.05$) are marked with asterisks.

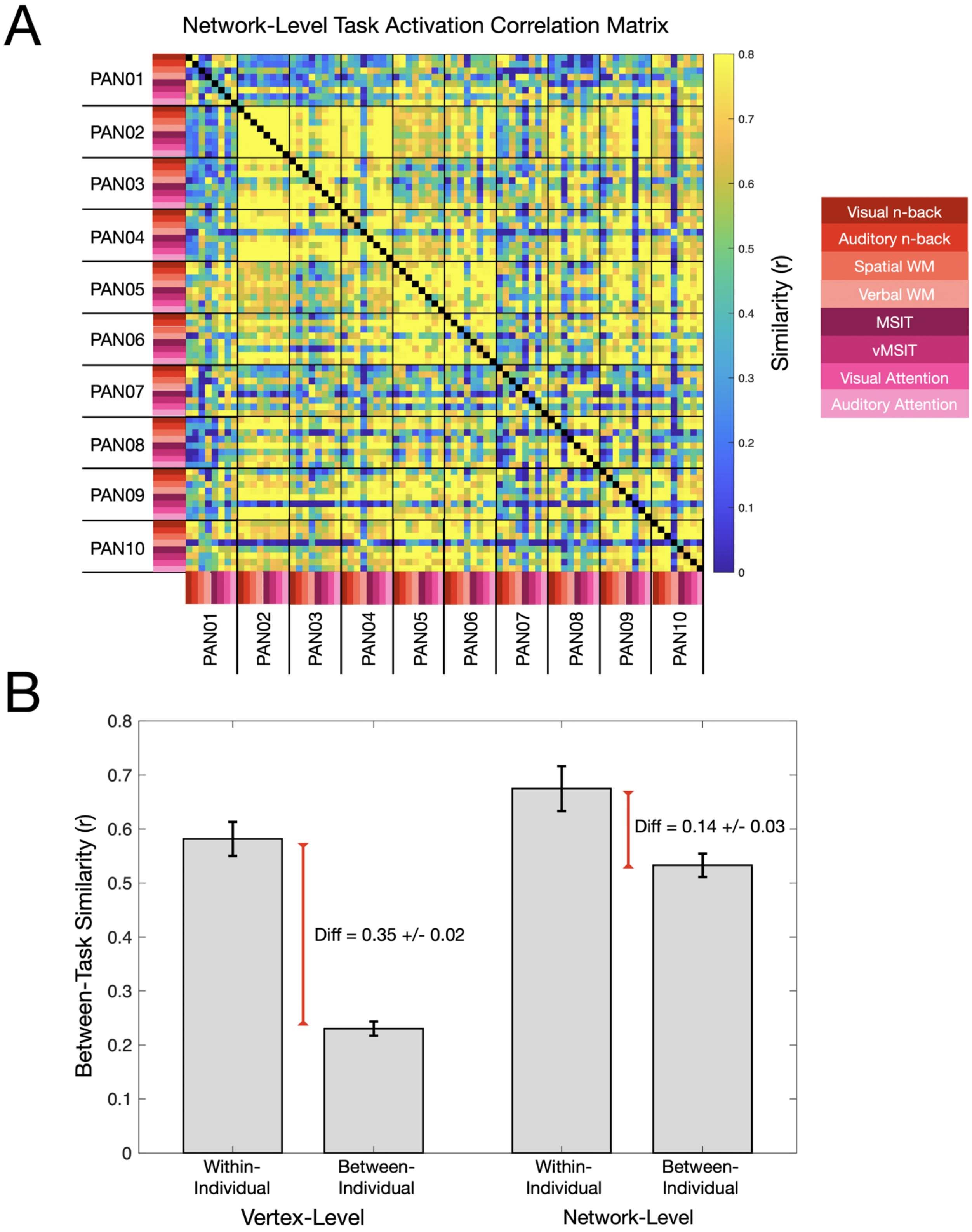


**Supplemental Figure S21. Network-level task activations are more similar between individuals than vertex-level task activations.** (A) Average LPFC network activations were calculated for each network for each individual and their patterns were correlated. Unlike for vertex-wise patterns, different individuals look similar to one another with no clear block structure. (B) Vertex-level task activation maps are much more similar across tasks for a single individual than between individuals (between-task: within-subject $r = 0.58 \pm 0.03$, between-subject $r = 0.23 \pm 0.01$, difference = $0.35 \pm 0.03$). Network-level task activations were also more similar within versus between subjects, but the difference was much smaller than in vertex-wise comparisons (within-subject $r = 0.67 \pm 0.04$, between-subject $r = 0.53 \pm 0.02$, difference = $0.14 \pm 0.01$, vertex-wise difference > network-wise difference paired t-test $t(9) = 16.0$, $p < 0.000001$). This suggests that the interindividual differences observed in LPFC activation patterns to the same task are driven more by interindividual differences in network topography than by differences in network recruitment.

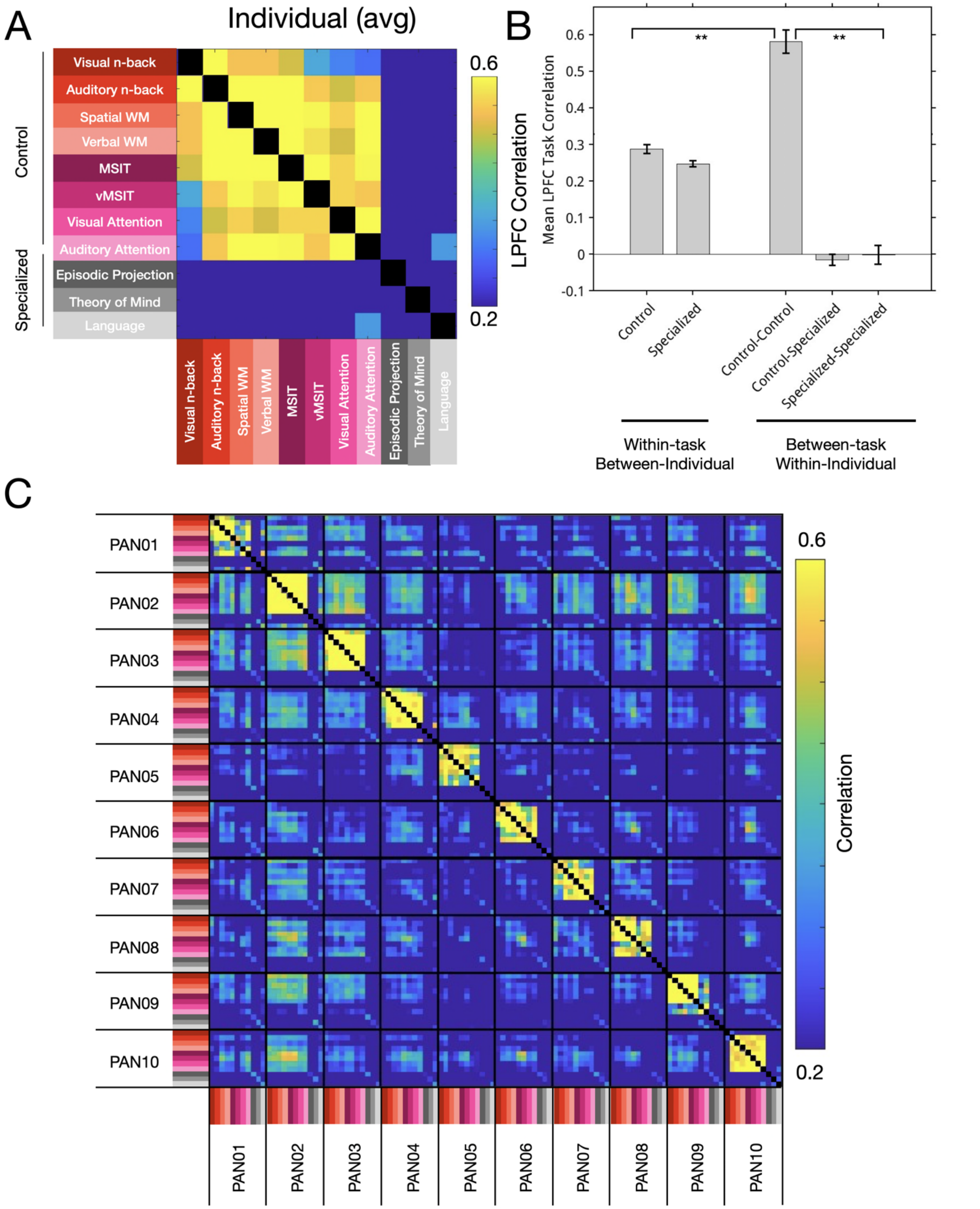


**Supplemental Figure S22. Within-individual cross-task cognitive control activation maps were more similar than cross-individual within-task control activation maps, within-individual control/domain-specific activation maps, and within-individual domain-specific activation maps.** (A) An average correlation matrix of LPFC activation similarity for all tasks within all individuals. (B) Cognitive control maps were quite similar to one another ($r = 0.58 \pm 0.03$), more so than they were to domain-specific activation maps within the same individual ($r = -0.02 \pm 0.02$, paired t-test $t(9) = 27.2$, $p < 0.00001$), or than domain-specific maps were to one another within the same individual ($r = 0.00 \pm 0.03$, paired t-test $t(9) = 11.4$, $p < 0.00001$). Across individuals and within-task, both domain-specific activation maps and cognitive control activations showed modest correlations ($r = 0.25 \pm 0.01$, $r = 0.27 \pm 0.03$, respectively). (C) A correlation matrix is shown for all individuals and tasks.

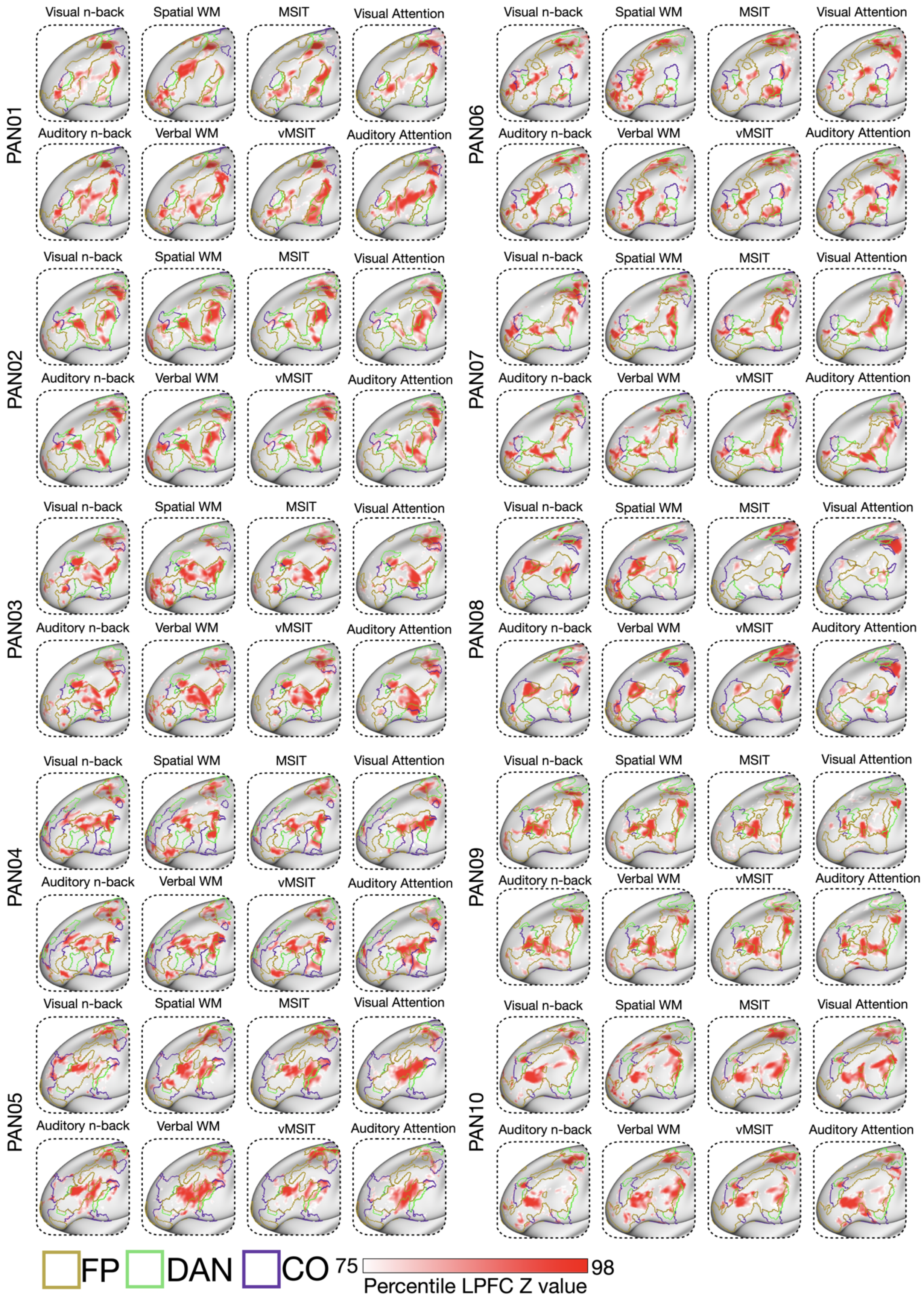
PAN01
PAN02
PAN03
PAN04
PAN05
PAN06
PAN07
PAN08
PAN09
PAN10
Visual n-back
Spatial WM
MSIT
Visual Attention
Auditory n-back
Verbal WM
vMSIT
Auditory Attention
FP
DAN
CO
75
98
Percentile LPFC Z value

**Supplemental Figure S23. Left hemisphere cognitive control task activation maps for all individuals.** Left hemisphere cognitive control task activation maps showing the top 25% activated LPFC vertices along with the outlines of the FP, CO, and DAN networks. As described in the main text, activation maps revealed a distributed set of highly active regions which were similar across the 8 different tasks, but unique between people. These regions tended to be located near or crossing the borders of the FP, CO, and DAN networks.

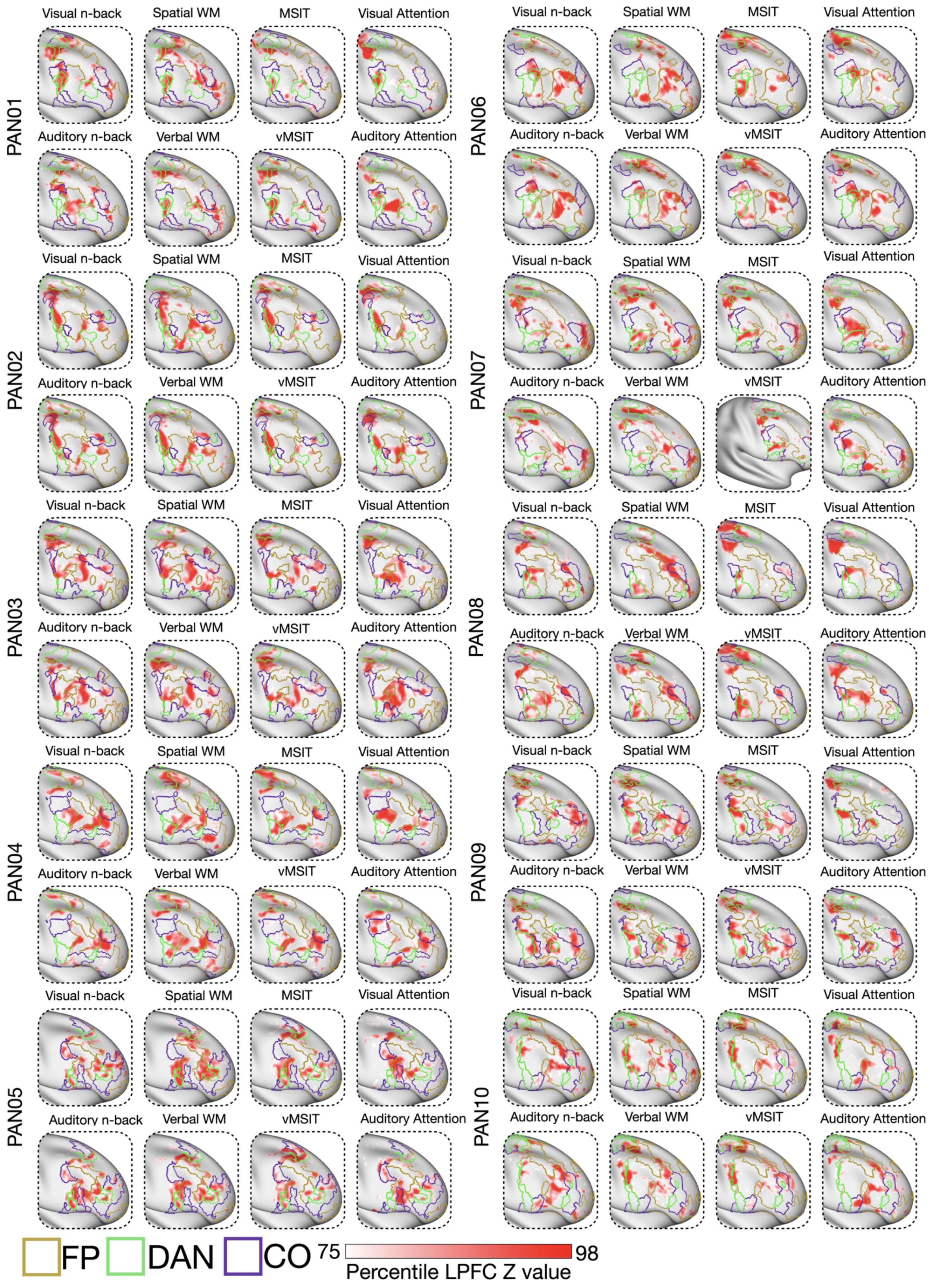
PAN01
PAN02
PAN03
PAN04
PAN05
PAN06
PAN07
PAN08
PAN09
PAN10
Visual n-back
Spatial WM
MSIT
Visual Attention
Auditory n-back
Verbal WM
vMSIT
Auditory Attention
FP
DAN
CO
75
98
Percentile LPFC Z value

**Supplemental Figure S24. Right hemisphere cognitive control task activation maps for all individuals.** Maps showing the top 25% activated LPFC right hemisphere vertices along with the outlines of the FP, CO, and DAN networks. As described in the main text, activation maps revealed a distributed set of highly active regions which were similar across the 8 different tasks, but unique between people. These regions tended to be located near or crossing the borders of the FP, CO, and DAN networks.

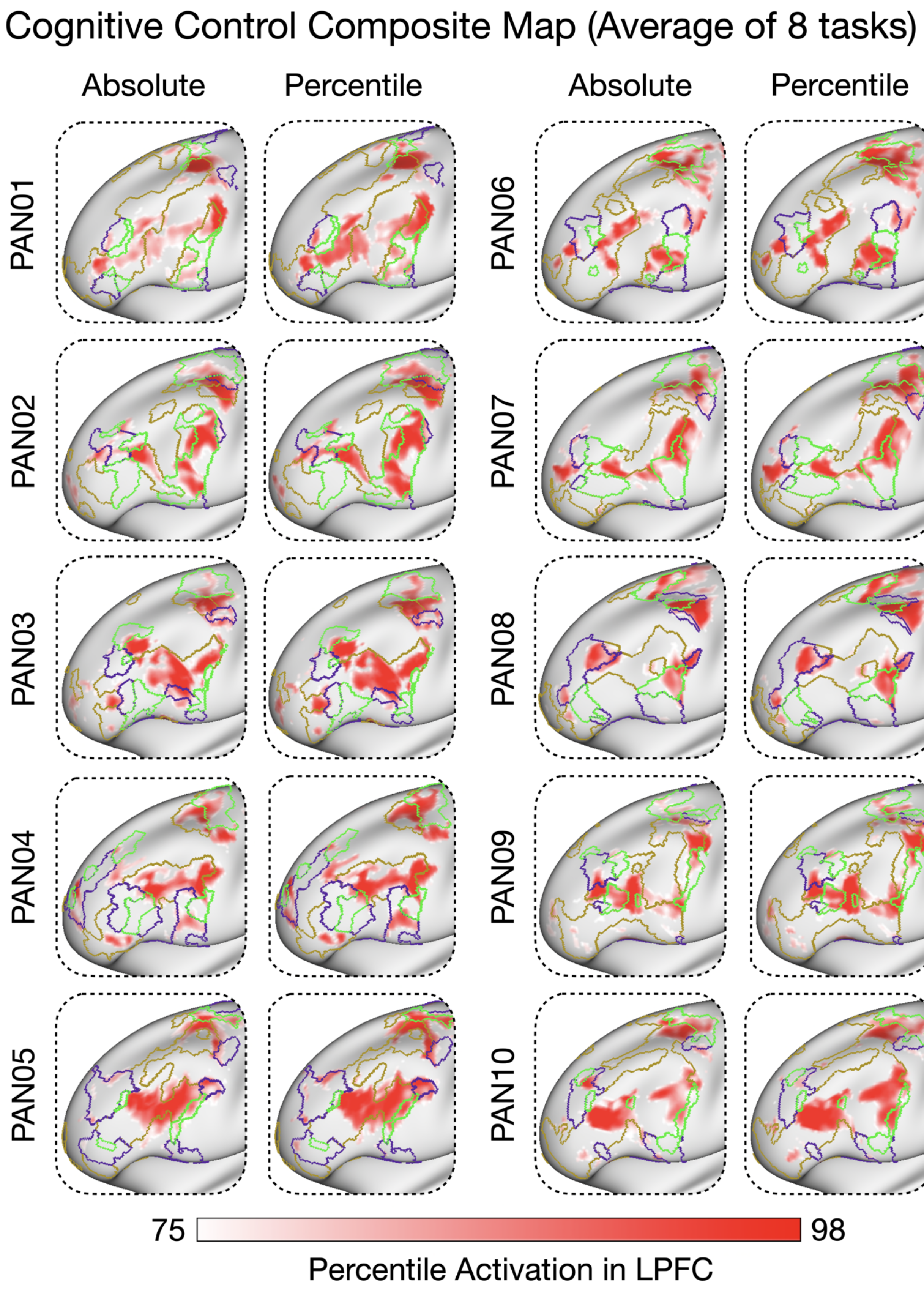
Cognitive Control Composite Map (Average of 8 tasks)
Absolute
Percentile
Absolute
Percentile
PAN01
PAN02
PAN03
PAN04
PAN05
PAN06
PAN07
PAN08
PAN09
PAN10
75
98
Percentile Activation in LPFC

**Supplemental Figure S25 - Composite cognitive control task maps.** Given the similarity of the cognitive control activations within an individual, composite control task maps were created by averaging the task activation z-statistic maps across all eight control tasks. Maps were made both averaging z-values on an absolute and percentile basis. The maps are quite similar to one another and also make obvious the individual differences between people in their control activation patterns as well as provide an easier visualization of border activity.

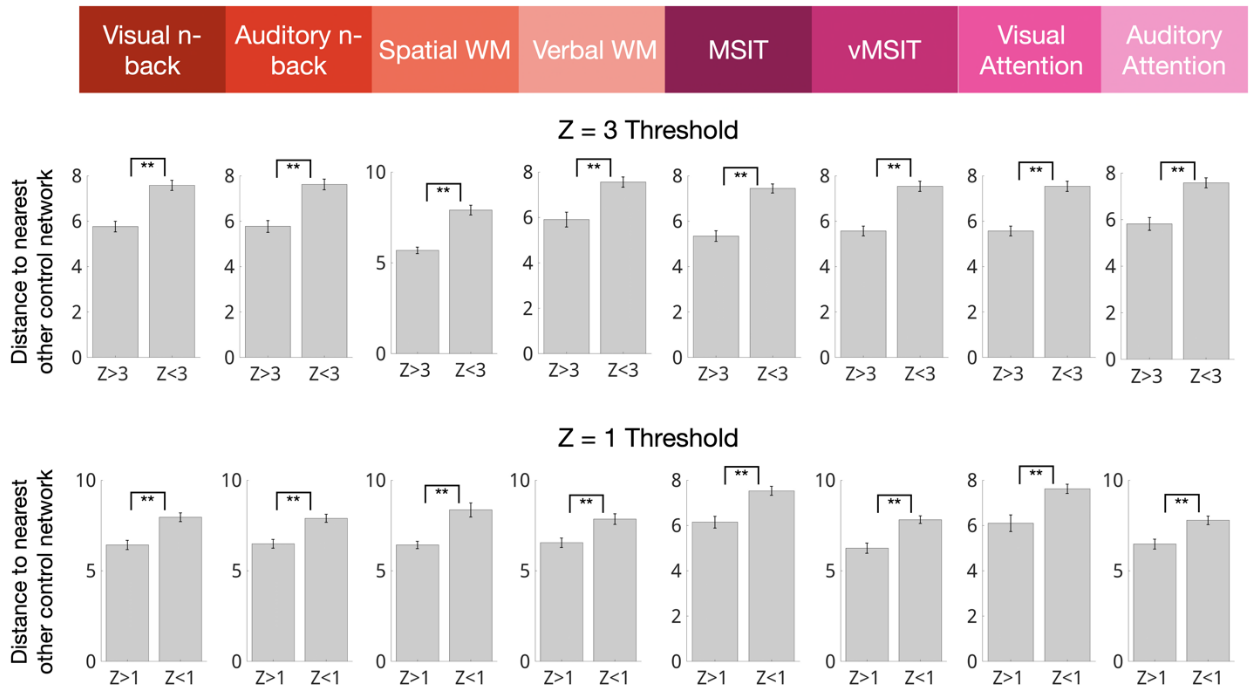


**Supplemental Figure S26. Cognitive control activations within control networks (FP, DAN, CO) tended to be near the borders where these networks meet.** For each task, LPFC vertices in the FP, CO, and DAN networks were split into "active" ($z > 3$, $z > 2$, $z > 1$) or less-active ($z < 3$, $z < 2$, $z < 1$) groups, and the mean geodesic distance from those vertices to the other two control networks was calculated. Indicated by asterisks, the more active group was closer to the other two control networks (all $p < 0.01$) across all 8 tasks across all 3 thresholds. Z > 2 is shown in the main text.

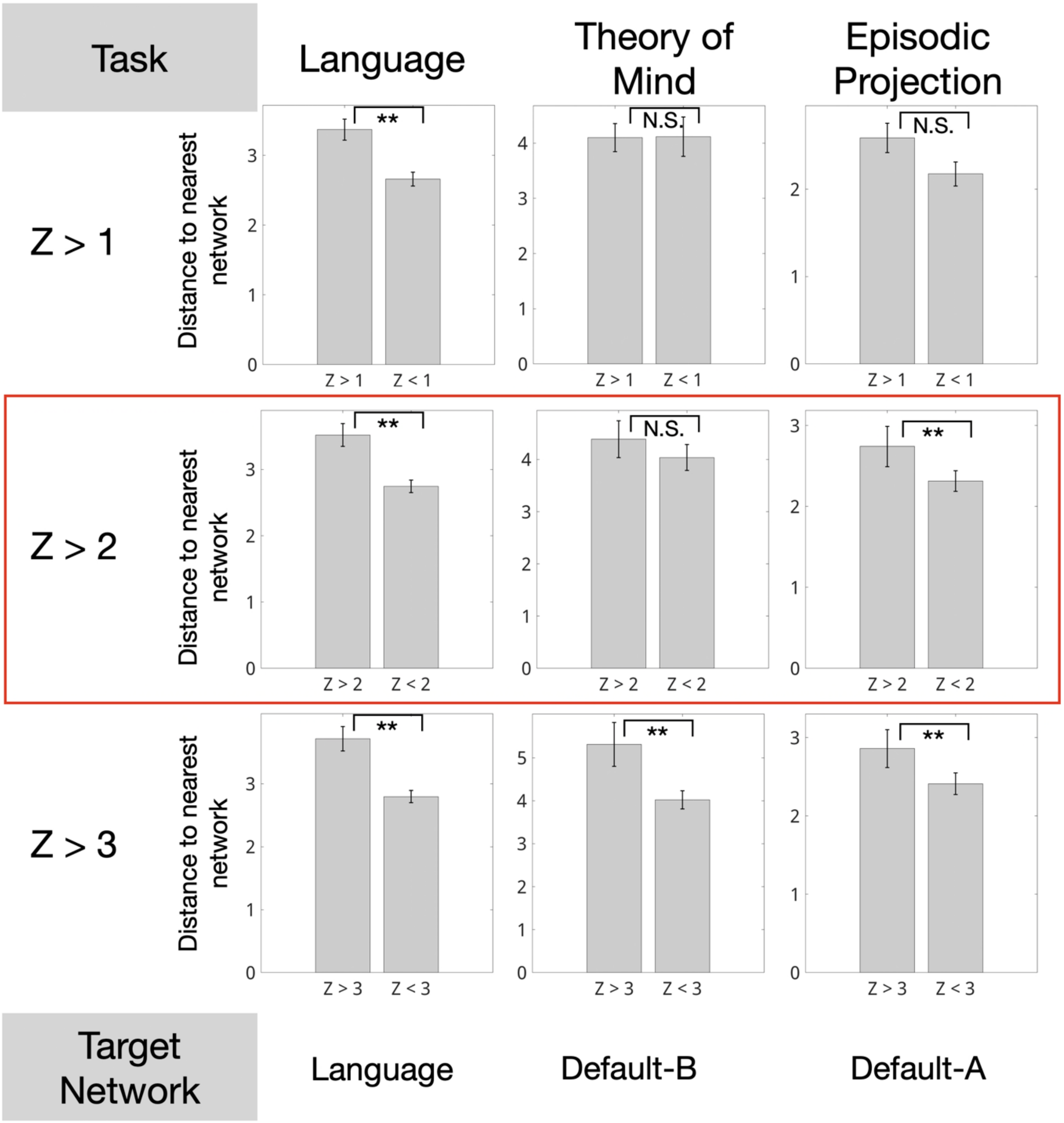


**Supplemental Figure S27. For domain-specific demands, highly active vertices within the target network are farther from network borders than less active vertices.**
We computed the geodesic distance from each LPFC vertex in the target networks (Language: Language, Episodic Projection: Default A, Theory of Mind: Default B) to the nearest other network. For language and episodic projection demands, more active vertices within the target network were significantly farther from other networks than less active vertices ($p < 0.01$). For theory of mind demands there was no significant relationship at $z > 1$ and $z > 2$, with a significant reversed effect only at $z > 3$. Stars mark $p < 0.01$.

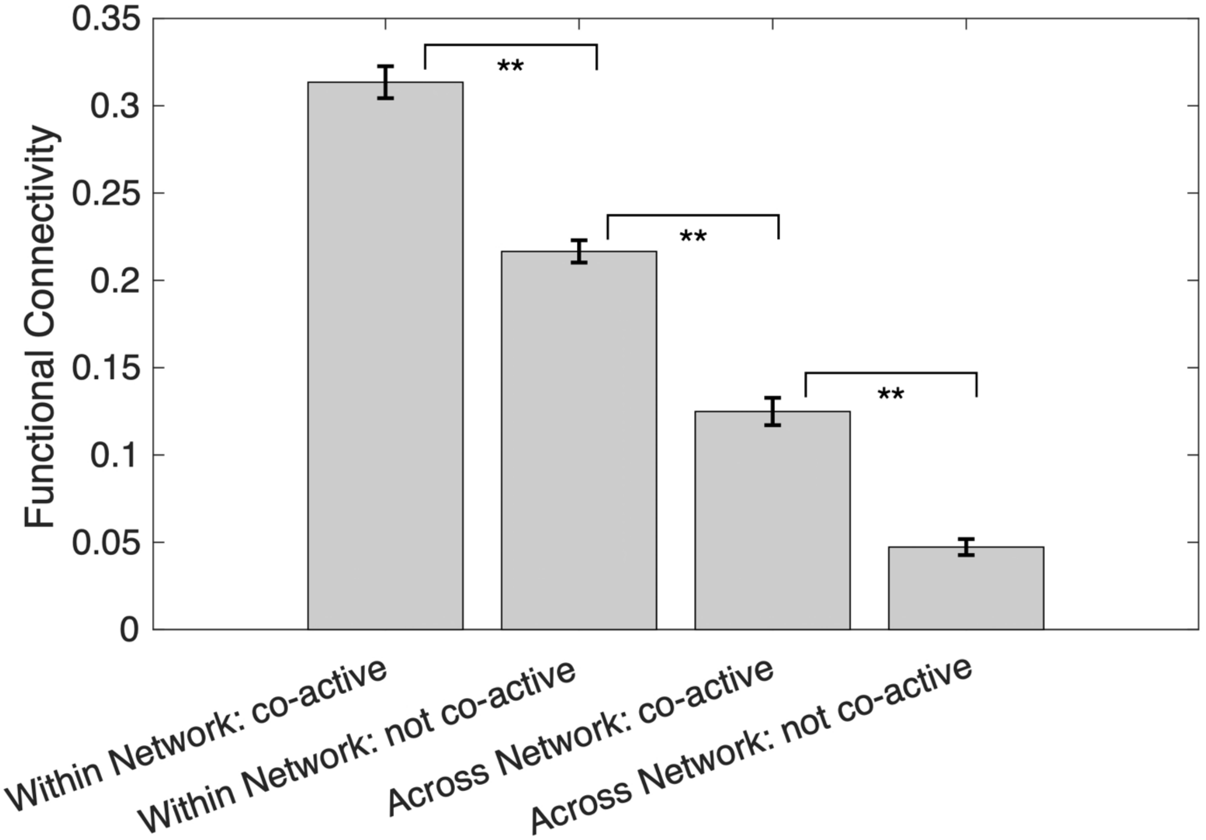


**Supplemental Figure S28. Co-active vertices from different control networks show higher FC than not co-active vertices, but lower than within-network FC, regardless of co-activity.** For each cognitive control task, vertices within FP, CO, and DAN were classified as active ($z > 2$) or not active ($z < 2$). Resting-state functional connectivity was measured between all vertex pairs, grouped according to (1) shared network membership and/or (2) co-activation status. A graded pattern of functional connectivity emerged: (1) co-active and in the same network > (2) not co-active but in the same network > (3) co-active in different networks > (4) not co-active in different networks. Starred comparisons indicate $p < 0.01$.

Theory of Mind Comparisons (Target Network DN-B > Comparison Network)

| Comparison Network | Raw p-value | Corrected p-value | T-stats | Cohen's d |
|---|---|---|---|---|
| FP | 3.337e-05 | **0.0002002** | 6.9515 | 2.1983 |
| DAN | 9.2e-07 | **5.54e-06** | 10.822 | 3.4222 |
| CO | 5.58e-06 | **3.349e-05** | 8.7088 | 2.754 |
| DN-A | 0.00013899 | **0.00083392** | 5.7452 | 1.8168 |
| LANG | 0.00147005 | **0.00882029** | 4.0374 | 1.2767 |
| SAL/PMN | 7.23e-06 | **4.336e-05** | 8.4358 | 2.6676 |

Episodic Projection Comparisons (Target Network DN-A > Comparison Network)

| Comparison Network | Raw p-value | Corrected p-value | T-stats | Cohen's d |
|---|---|---|---|---|
| FP | 6.41e-06 | **3.845e-05** | 8.5619 | 2.7075 |
| DAN | 5.9e-07 | **3.57e-06** | 11.4015 | 3.6055 |
| CO | 1.9e-07 | **1.17e-06** | 12.9986 | 4.1105 |
| DN-B | 1.51e-06 | **9.07e-06** | 10.2045 | 3.2269 |
| LANG | 1.9e-07 | **1.12e-06** | 13.0632 | 4.131 |
| SAL/PMN | 1.6e-07 | **9.4e-07** | 13.3251 | 4.2138 |

Language Processing Comparisons (Target Network LANG > Comparison Network)

| Comparison | Raw p-value | Corrected p-value | T-stats | Cohen's d |
|---|---|---|---|---|
| FP | 8.05e-06 | **4.833e-05** | 8.3232 | 2.632 |
| DAN | 3.971e-05 | **0.00023826** | 6.7959 | 2.1491 |
| CO | 0.00011666 | **0.00069994** | 5.8852 | 1.8611 |
| DN-A | 8.18e-06 | **4.907e-05** | 8.3075 | 2.6271 |
| DN-B | 4.48e-06 | **2.689e-05** | 8.9465 | 2.8291 |
| SAL/PMN | 5.59e-06 | **3.355e-05** | 8.7069 | 2.7534 |

**Supplemental Table S1: Corrected and uncorrected p-values, t-stats and Cohen's d for all network task activation comparisons to high level task domains.** Bolded corrected p-values are significant (corrected $p < 0.05$).

**Language Network/Language Task Overlap (Individual versus Comparison)**

| Comparison | Raw p-value | Corrected p-value | T-stats | Cohen’s d |
|---|---|---|---|---|
| Group | 0.00603 | **0.0121** | 3.13 | 1.04 |
| Others | 0.000675 | **0.00135** | 4.57 | 1.52 |

**Default Network A /Episodic Projection Task Overlap**

| Comparison | Raw p-value | Corrected p-value | T-stats | Cohen’s d |
|---|---|---|---|---|
| Group | 0.000865 | **0.00173** | 4.4 | 1.47 |
| Others | 0.0178 | **0.0357** | 2.47 | 0.823 |

**Default Network B /Theory of Mind Task Overlap**

| Comparison | Raw p-value | Corrected p-value | T-stats | Cohen’s d |
|---|---|---|---|---|
| Group | 0.00068 | **0.00136** | 4.56 | 1.52 |
| Others | 1.9e-05 | **3.8e-05** | 7.47 | 2.49 |

**LPFC vs. Non-LPFC Network/Task Overlap**

| Network/Task | Raw p-value | Corrected p-value | T-stats | Cohen’s d |
|---|---|---|---|---|
| Language/Language | 0.000148 | **0.000148** | 5.7 | 1.9 |
| DN-A/Episodic Projection | 0.0271 | **0.0271** | 2.21 | 0.738 |
| DN-B/Theory of Mind | 0.000307 | **0.000307** | 5.14 | 1.71 |
| Cross Task Average | 2.68e-05 | **2.68e-05** | 7.15 | 2.38 |

**Supplemental Table S2: Corrected and uncorrected p-values, t-stats and Cohen’s d for all network task activation overlap comparisons.** Bolded corrected p-values are significant (corrected $p < 0.05$).

| Task | Network | Mean Z Value | T-stat | Raw p-value | Cohen's d | Corrected p-value |
|---|---|---|---|---|---|---|
| Visual N-back | **FP** | 1.41 | 7.25 | 0 | 2.29 | **0.0002** |
| | **DAN** | 1.58 | 10.34 | 0 | 3.27 | **0** |
| | **CO** | 1.05 | 6.77 | 0 | 2.14 | **0.0003** |
| | DN-A | 0.18 | 0.47 | 0.326 | 0.15 | 1 |
| | DN-B | -0.76 | -4.26 | 0.9989 | -1.35 | 1 |
| | LANG | 0.47 | 2.16 | 0.0297 | 0.68 | 0.2081 |
| | **SAL/PMN** | 1.03 | 4.98 | 0.0004 | 1.57 | **0.0027** |
| Auditory N-back | **FP** | 1.55 | 9.5 | 0 | 3 | **0** |
| | **DAN** | 1.55 | 5.05 | 0.0003 | 1.6 | **0.0024** |
| | **CO** | 0.93 | 4.15 | 0.0013 | 1.31 | **0.0088** |
| | DN-A | -0.42 | -1.2 | 0.8692 | -0.38 | 1 |
| | DN-B | -1 | -6.46 | 0.9999 | -2.04 | 1 |
| | **LANG** | 1.2 | 3.79 | 0.0021 | 1.2 | **0.015** |
| | SAL/PMN | 0.34 | 1.24 | 0.1226 | 0.39 | 0.8582 |
| Spatial WM | **FP** | 2.81 | 6.31 | 0.0001 | 1.99 | **0.0005** |
| | **DAN** | 2.78 | 6.74 | 0 | 2.13 | **0.0003** |
| | CO | 0.61 | 2.36 | 0.0212 | 0.75 | 0.1483 |
| | DN-A | 0.91 | 1.56 | 0.0765 | 0.49 | 0.5356 |
| | DN-B | -0.28 | -0.64 | 0.732 | -0.2 | 1 |
| | LANG | 0.2 | 0.6 | 0.2803 | 0.19 | 1 |
| | **SAL/PMN** | 1.64 | 4.53 | 0.0007 | 1.43 | **0.005** |
| Verbal WM | **FP** | 2.27 | 9.11 | 0 | 2.88 | **0** |
| | **DAN** | 2 | 5.41 | 0.0002 | 1.71 | **0.0015** |
| | CO | 0.67 | 1.96 | 0.0407 | 0.62 | 0.2849 |
| | DN-A | 0.17 | 0.4 | 0.3504 | 0.13 | 1 |
| | DN-B | -0.7 | -3.46 | 0.9964 | -1.09 | 1 |
| | **LANG** | 1.14 | 5.24 | 0.0003 | 1.66 | **0.0019** |
| | SAL/PMN | 1.19 | 2.4 | 0.02 | 0.76 | 0.1402 |
| MSIT | FP | 0.31 | 1.02 | 0.1667 | 0.32 | 1 |
| | **DAN** | 1.68 | 4.3 | 0.001 | 1.36 | **0.007** |
| | CO | 0.55 | 1.79 | 0.0537 | 0.57 | 0.3757 |
| | DN-A | -1.23 | -4.27 | 0.999 | -1.35 | 1 |
| | DN-B | -1.39 | -6.76 | 1 | -2.14 | 1 |
| | LANG | -0.24 | -1.18 | 0.8652 | -0.37 | 1 |
| | SAL/PMN | -0.24 | -0.73 | 0.7591 | -0.23 | 1 |
| vMSIT | **FP** | 1.09 | 4.32 | 0.001 | 1.37 | **0.0068** |
| | **DAN** | 2.68 | 13.76 | 0 | 4.35 | **0** |

| | | | | | | |
|---|---|---|---|---|---|---|
| | **CO** | 1.1 | 7.96 | 0 | 2.52 | **0.0001** |
| | DN-A | -1.15 | -2.45 | 0.9817 | -0.78 | 1 |
| | DN-B | -1.8 | -5.9 | 0.9999 | -1.87 | 1 |
| | LANG | 0.15 | 0.44 | 0.3352 | 0.14 | 1 |
| | SAL/PMN | -0.35 | -1.05 | 0.8395 | -0.33 | 1 |
| Visual Attention | FP | 0.47 | 1.41 | 0.0954 | 0.45 | 0.668 |
| | **DAN** | 0.94 | 3.64 | 0.0027 | 1.15 | **0.0189** |
| | CO | 0.26 | 0.94 | 0.1857 | 0.3 | 1 |
| | DN-A | -0.65 | -2.17 | 0.9711 | -0.69 | 1 |
| | DN-B | -0.79 | -4.66 | 0.9994 | -1.47 | 1 |
| | LANG | -0.24 | -0.74 | 0.7606 | -0.23 | 1 |
| | SAL/PMN | -0.25 | -0.68 | 0.7445 | -0.22 | 1 |
| Auditory Attention | FP | 0.98 | 2.94 | 0.0083 | 0.93 | 0.058 |
| | **DAN** | 1.37 | 6.45 | 0.0001 | 2.04 | **0.0004** |
| | **CO** | 1.31 | 5.97 | 0.0001 | 1.89 | **0.0007** |
| | DN-A | -0.67 | -2.82 | 0.99 | -0.89 | 1 |
| | DN-B | -1.17 | -6.44 | 0.9999 | -2.04 | 1 |
| | **LANG** | 1.55 | 3.57 | 0.003 | 1.13 | **0.0211** |
| | SAL/PMN | 0.09 | 0.25 | 0.4046 | 0.08 | 1 |

**Supplemental Table S3: Corrected and uncorrected p-values, t-stats and Cohen's d for all LPFC network task activations to individual cognitive control tasks.** Bolded corrected p-values are significant (corrected $p < 0.05$).
